# A method of comprehensive sequencing analysis of the small RNA fragmentome (RiboMarker^®^)

**DOI:** 10.1101/2025.11.12.688114

**Authors:** Rachel C. Clark, Aidan C. Manning, Jonathan M. Howard, Sergio Barberán-Soler, Sergei A. Kazakov

## Abstract

Circulating cell-free nucleic acids (cfDNA and cfRNA) found in blood and other biofluids are promising biomarkers for cancer. However, current methods exploiting tumor-derived cfDNA (ctDNA) are not sensitive enough in detecting minimal residual disease and early stages of cancer when it is more treatable. Small RNAs and RNA fragments (sRNA) can potentially provide higher detection sensitivity and specificity than ctDNA. Sequencing analysis of the variety of sRNAs representing the entire RNA fragmentome would improve our understanding of their roles in cancer development and help to discover novel sRNA biomarkers for cancer diagnostics and personalized treatments. However, conventional methods of sRNA-Seq library preparation are limited to detection of sRNA with 5’-P and 3’-OH ends that represent only less than 10% of the whole RNA fragmentome, whereas sRNAs having different phosphorylation statuses (P or OH) of their termini are hidden. Although recently developed sRNA-Seq methods allow detection of most sRNA (including the hidden ones) simultaneously, these methods cannot both detect and distinguish among the individual RNAs with differing termini combinations (RNA Types). Here we describe the RiboMarker^®^ platform for preparation of sRNA sequencing libraries that addresses these shortcomings. It uses distinctive enzymatic pretreatment(s) of RNA samples that can both detect all and enrich for individual sRNA Types upfront of sequencing library preparation. The RiboMarker^®^ platform has the potential capability to both identify and detect with enhanced sensitivity low abundance sRNA biomarkers of specific RNA classes and their termini.

## INTRODUCTION

Circulating cell-free DNA (cfDNA) found in human plasma and other biofluids is considered one of the most potent liquid biopsy biomarkers for cancer (Song et al. 2022; Brito-Rocha et al. 2023). However, current methods for the detection of tumor-derived cfDNA (ctDNA) are not sensitive enough to avoid often false-positive or negative results in detecting cancer, especially at early stages when it is more treatable (Fiala and Diamandis 2020; Pons-Belda et al. 2021). For example, ctDNA represents only 0.01–0.0001% against the background of non-cancerous cfDNA at low tumor burden (Siravegna et al. 2017). Meanwhile, circulating cell-free RNA (cfRNA) is emerging as a novel class of diagnostic, prognostic, and predictive biomarkers that can provide higher sensitivity and specificity for cancer detection compared to ctDNA (Fish et al. 2018; Pardini et al. 2019; Ning et al. 2023; Tao et al. 2023). Unlike ctDNA, tumor-associated cfRNAs (ctRNA) are released not only from dead tumor cells, but also from both alive cancerous and non-cancerous cells, which surround the tumor and provide immune response to cancer (Cabús et al. 2022; Liu et al. 2023). This and the fact that RNA is transcribed in multiple copies from various genomic DNA regions (including the ones that are only produced by cancer) contribute to the higher ctRNA abundance than ctDNA both in cells and in circulation (Larson et al. 2021; Vibert et al. 2022). In addition, dramatic changes in the RNA expression profile in tumors, overexpression of specific ctRNA, and dysregulated RNA post-transcriptional events (including alternative splicing and formation of chimeric RNAs that are detectable only in the transcriptome) contribute to the higher abundance and complexity of the ctRNA landscape (Cabús et al. 2022; Ning et al. 2023; Wang et al. 2024).

Most of the cfRNA molecules (∼95%) found in human plasma are small processed RNAs and RNA fragments of <45 nucleotides (nt) in length, hereinafter together referred to as sRNA (Akat et al. 2019; Galvanin et al. 2019; Giraldez et al. 2019; Wang et al. 2024). These sRNAs represent the primary RNA transcripts and precursor RNAs of various RNA classes (Tosar et al. 2020). Main RNA classes considered in this study include ribosomal RNA (rRNA), transfer RNA (tRNA), microRNA (miRNA), small nuclear RNA (snRNA), small nucleolar RNA (snoRNA), messenger RNA (mRNA) (including exons and UTRs), long non-coding RNA (lncRNA) and Piwi interacting RNA (piRNA) (Vickers et al. 2015; Hulstaert et al. 2020). Until recently, the analysis of sRNAs has been primarily focused on miRNA (Wang et al. 2018; Pardini et al. 2019; Wu et al. 2021). However, there are other sRNA classes with greater diversity and abundance than miRNA and, therefore, have greater biomarker potential that could reliably and with higher specificity and sensitivity assess the state of a disease (Vickers et al. 2015; Akat et al. 2019; Yao et al. 2020; Cabús et al. 2022; Liu et al. 2022). Furthermore, certain sRNAs derived from mRNA and lncRNA, which are mostly represented by multiple overlapping sequences, contain information about biological phenotypes that inform about tissues of cancer origin and cancer subtypes (Akat et al. 2019; Giraldez et al. 2019; Larson et al. 2021; Chen et al. 2022; De Sota et al. 2024). sRNAs with unique (defined) sequences aligned to tRNAs (Wang et al. 2022; Chen and Zhou 2023; Di Fazio and Gullerova 2023), snRNAs (Pardini et al. 2019; Su et al. 2022), and snoRNAs (Gao et al. 2024) have also been identified as potential cancer biomarkers.

sRNAs are processed from the parent RNA transcripts by one or more intracellular and/or extracellular ribonucleases (RNases) through one or more RNA cuts (Shi et al. 2022; Shigematsu and Kirino 2022; Chen and Zhou 2023; Lai et al. 2023; Tosar et al. 2024). Protective RNA-protein complexes, encapsulation into naturally occurring lipid extracellular vesicles (EVs), and/or RNA secondary structures help prevent further cleavage of processed sRNAs released from cells into circulation (De Sota et al. 2024; Crocker et al. 2022; Shi et al. 2022; Tosar et al. 2024). A major proportion of extracellular sRNAs are found outside EVs (Tosar et al. 2020; Jia et al. 2021). Depending on the RNases involved in the RNA cleavage and naturally occurring RNA termini, the produced sRNAs may feature specific combinations of RNA 5’ and 3’ ends (hereafter called RNA Types) having different phosphorylation states (Fig. 1).

**FIGURE 1.**
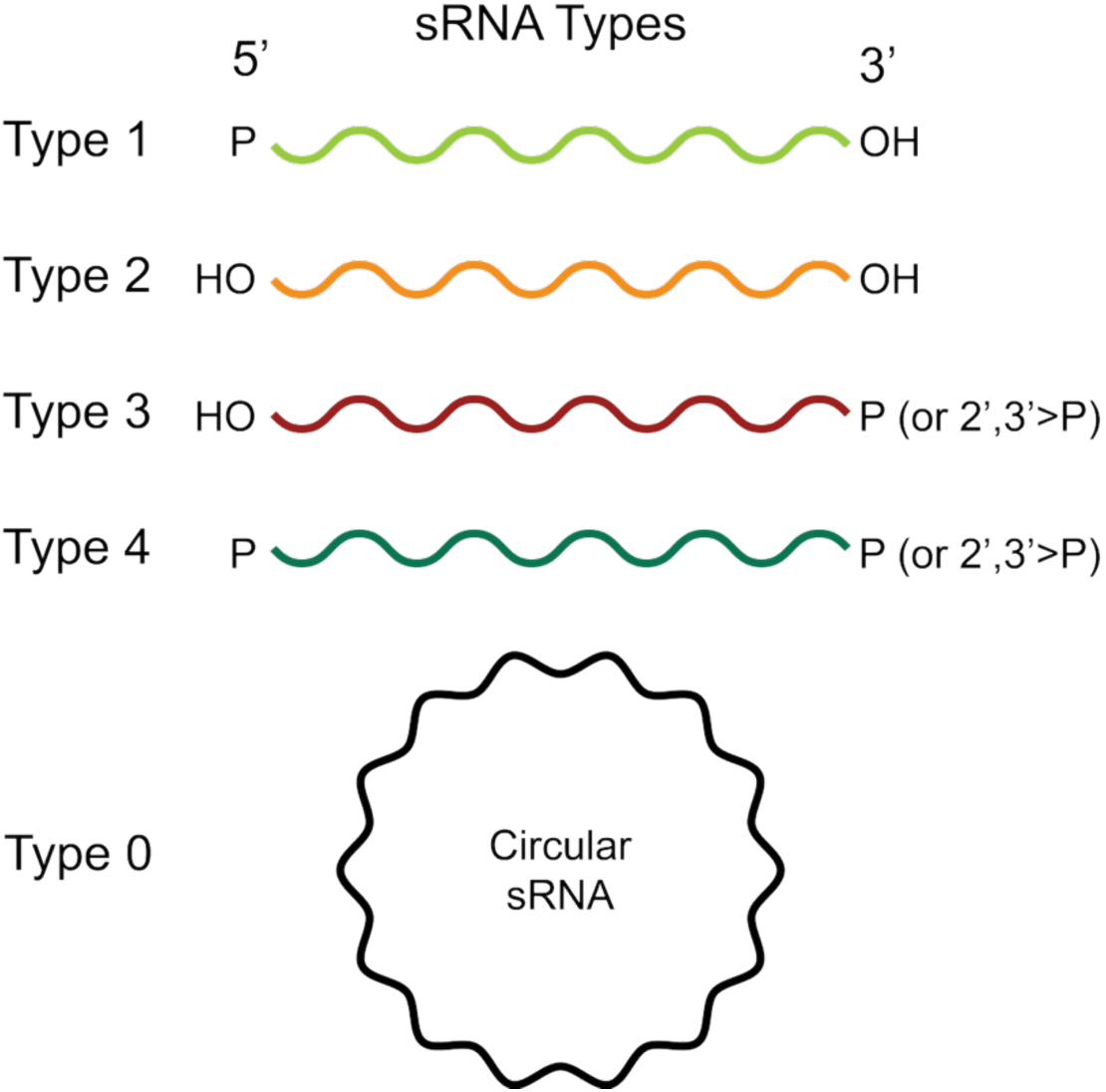
RNA Types representing small RNA molecules and RNA fragments (sRNAs) having different phosphorylation states of their ends (Types 1 through 4) or their circular form (Type 0) with no ends.

Original methods for sRNA-Seq library preparation, which have been used to discover sRNAs and are still commonly used for their analysis, capture only RNA Type 1 [5’-P and 3’-OH] including miRNAs and some other sRNAs having the same ends, whereas RNA Type 2 [5’-OH and 3’-OH], RNA Type 3 [5’-OH and 3’-P] and RNA Type 4 [5’-P and 3’-P] are not incorporated into sequencing libraries (Crocker et al. 2022; Shi et al. 2022; Shigematsu and Kirino 2022). However, the sRNAs with 3’-OH ends (RNA Types 1 and 2) account for only ∼10% of the human cellular RNA fragmentomes, while sRNAs having 3’-phosphorylated ends (RNA Types 3 and 4) are “hidden” and cannot be detected by these methods (Lai et al. 2023). Moreover, some sRNAs (e.g., tRNA fragments) contain modified 5’-end (e.g., 5’ cap) or internal nucleotide(s) that can interfere with reverse transcription, which prevents or allows only partial incorporation of these sRNA sequences into libraries (Wang et al. 2024; Crocker et al. 2022; Shi et al. 2022; Verwilt et al. 2023).

Current methods for detecting these hidden sRNAs by sequencing could be divided into two groups. The first group uses wild-type T4 polynucleotide kinase (PNK), which has both phosphatase and kinase activity in the presence of ATP, for treating sRNAs to erase differences between the phosphorylation states of the sRNA ends by converting them to the RNA Type 1 form (Akat et al. 2019; Galvanin et al. 2019; Giraldez et al. 2019; Crocker et al. 2022; Shi et al. 2022; Liu et al. 2022; Shigematsu and Kirino 2022; Solaguren-Beascoa et al. 2023). Although this approach allows simultaneous analysis of all RNA Types and RNA classes, the generated sRNA sequencing libraries contain very high proportions of rRNA fragments that overshadow the other RNA classes. While deep sequencing and/or commonly used rRNA depletion prior to sequencing library preparation could assist in the discovery of biomarker candidates within all other RNA classes, such approaches significantly decrease the sensitivity of their detection. The second group focuses exclusively on sRNAs with specific RNA ends such as 5’-OH; 3’-OH; 3’-P or 2’,3’>P and cannot distinguish between the phosphorylation state at the opposite 5’ or 3’ end (Peach et al. 2015; Honda et al. 2016; Jia et al. 2021; Kugelberg et al. 2021; Crocker et al. 2022; Del Piano et al. 2022; Shi et al. 2022; Shigematsu and Kirino 2022; Lai et al. 2023). Except for RNA Type 1, none of these library preparation methods can specifically detect and compare individual sRNAs of Types 2, 3, and 4, representing the absolute majority of sRNAs as indicated above. Moreover, these known methods cannot detect all sRNA Types simultaneously and distinguish each individual Type specifically while using the same protocol for the preparation of sequencing libraries. Because of these method-specific biases (Zhuang et al. 2012a; Wright et al. 2019; Shi et al. 2021a), relative abundances of the different sRNA Types determined by different methods of library preparation cannot be accurately compared. Furthermore, all these reported methods require purification of intermediate reaction products, e.g. by gel-electrophoresis, phenol-chloroform or TRIzol extraction and ethanol precipitation (Akat et al. 2019; Giraldez et al. 2019; Solaguren-Beascoa et al. 2023; Kugelberg et al. 2021; Del Piano et al. 2022; Shi et al. 2021a; Lai et al. 2023), which leads to unavoidable loss of RNA during the purification, limiting sensitivity and reproducibility of sRNA quantification by sequencing.

To address these shortcomings, we developed the RiboMarker^®^ platform that enables comprehensive profiling of total sRNA content in biological samples, including the detection of all RNA Types simultaneously and discrimination of the individual Types. RiboMarker^®^ uses distinctive enzymatic pretreatment(s) of RNA samples, determining the sRNA Type(s) to be sequenced upfront of a universal (the same for all different pretreatment protocols) library preparation protocol. It has the potential capability to both identify and then detect with enhanced sensitivity low-abundance sRNA biomarkers for minimal residual disease or cancer at early stages.

## RESULTS AND DISCUSSIONS

### Design of RiboMarker^®^ platform

The RiboMarker^®^ platform has two key components: 1) RNA Type-specific pretreatments of RNA samples to be sequenced; and 2) a universal library preparation protocol for the pretreated RNA samples.

The first key component is a customizable group of RNA enzymatic pretreatment(s) that enriches for the specific sRNA Type(s) of interest while depleting other Type(s) in sequencing libraries. The RNA Type(s)-specific pretreatment protocols are schematically presented in Table 1. We applied column-based purification after each pretreatment step to concentrate reaction products, to exchange buffers, and to remove undesirable enzymes, which may interfere with the next reaction step.

**TABLE 1.** Selected RNA Type-specific protocols for the preparation of RiboMarker^®^ sequencing libraries.

| RiboMarker® Protocols | T[1+2] | T[2] | T[1+2+3+4] | T[3+4] | T[3] | T[4] |
| --- | --- | --- | --- | --- | --- | --- |
| RNA Types in the sample | 1, 2, 3, 4 | 1, 2, 3, 4 | 1, 2, 3, 4 | 1, 2, 3, 4 | 1, 2, 3, 4 | 1, 2, 3, 4 |
| Pretreatment # 1 | – | Rnl1 + Rnl2 ligases (+ATP) | PNK (-ATP) pH 6.0 | PNK, 3'-minus (+ATP) pH 7.6 | Terminator 5'-p - dependent exonuclease | RtcB ligase |
| RF Types conversion | – | 1 → 0 | 3 → 2<br>4 → 1 | 2 → 1<br>3 → 4 | Degrade 1, 4 | 3 → 0 |
| Pretreatment # 2 | – | – | – | Rnl1 + Rnl2 ligases (+ATP) | PNK, 3'-minus (+ATP) pH 7.6 | PNK, 3'-minus (+ATP) pH 7.6 |
| RNA Types conversion | – | – | – | 1 → 0 | 2 → 1<br>3 → 4 | 2 → 1 |
| Pretreatment # 3 | – | – | – | PNK (-ATP) pH 6.0 | Rnl1 + Rnl2 ligases (+ATP) | Rnl1 + Rnl2 ligases (+ATP) |
| RNA Types conversion | – | – | – | 4 → 1 | 1 → 0 | 1 → 0 |
| Pretreatment # 4 | – | – | – | – | PNK (-ATP) pH 6.0 | PNK (-ATP) pH 6.0 |
| RNA Types conversion | – | – | – | – | 4 → 1 | 4 → 1 |
| Library preparation Protocol | T[1+2] | T[1+2] | T[1+2] | T[1+2] | T[1+2] | T[1+2] |
| RNA Types depleted from library | 3, 4 | 1, 3, 4 | None | 1, 2 | 1, 2, 4 | 1, 2, 3 |
| RNA Types enriched in library | 1, 2 (3'-OH Types) | 2 | 1, 2, 3, 4 (All Types) | 3, 4 (3'-P Types) | 3 | 4 |

Protocol T[1+2] provides the detection of RNA Types 1 and 2 simultaneously. This protocol is also used as a universal library preparation method in combination with all the other pretreatment steps described below.

Protocol T[2] has been designed to enrich for and detect RNA Type 2 with a single pretreatment by a mix of T4 RNA ligase 1 (Rnl1) and T4 RNA ligase 2 (Rnl2). This pretreatment results in the circularization of RNA Type 1 by ligating their 5’-P and 3’-OH ends and preventing incorporation of such RNA into the sequencing libraries.

Protocol T[1+2+3+4] has been designed to detect all RNA Types (Types 1, 2, 3, and 4) with a single pretreatment using T4 polynucleotide kinase (PNK) at pH 6.0 in the absence of ATP. Such pretreatment results in specific dephosphorylation of 3’-P or 2’,3’-cyclic phosphate (2’,3’>P) ends in RNA Types 3 and 4 without affecting the 5’ ends of RNA Types 1 and 2.

Protocol T[3+4] has been designed to enrich for and detect RNA Types 3 and 4 with two sequential pretreatment steps, including: (i) simultaneously run in one mixture reaction with PNK mutant having no 3’-end phosphatase activity (PNK, 3’-minus) in the presence of ATP at pH 7.6 to phosphorylate 5’ ends of RNA Types 2 and 3, converting them to RNA Types 1 and 4, respectively, and with Rnl1+Rnl2 mix to circularize RNA Type 1 excluding them from the sequencing library; and (ii) by PNK at pH 6.0 in the absence of ATP to dephosphorylate 3’ ends of former RNA Type 3 (converted to RNA Type 4) and original RNA Type 4 converting both of them to RNA Type 1.

Protocol T[3] has been designed to enrich for and detect RNA Type 3 with three sequential treatment steps, including: (i) by Terminator 5’-Phosphate-dependent exonuclease, which specifically digests RNA Types 1 and 4 having 5’-P ends excluding them from sequencing libraries; (ii) simultaneously run in one mixture reaction with PNK, 3’-minus in the presence of ATP at pH 7.6 to convert RNA Types 2 and 3, to RNA Types 1 and 4, respectively and Rnl1+Rnl2 mix to circularize RNA Type 1 excluding them from the sequencing library; and (iii) by PNK at pH 6.0 in the absence of ATP to dephosphorylate 3’ ends of former RNA Type 3 (converted to RNA Type 4) converting these to RNA Type 1.

Protocol T[4] has been designed to enrich for and detect RNA Type 4 with three sequential pretreatment steps, including: (i) by RtcB ligase to circularize RNA Type 3 through ligation of its 5’-OH and 3’-phosphorylated ends and preventing their incorporation in the sequencing libraries; (ii) simultaneously run in one mixture reaction with PNK, 3’-minus in the presence of ATP at pH 7.6 to phosphorylate 5’ ends of RNA Type 2, and Rnl1+Rnl2 mix to circularize RNA Type 1 and former RNA Type 2 converted to RNA Type 1; and (iii) by PNK at pH 6.0 in the absence of ATP to dephosphorylate of 3’ ends of RNA Type 4.

The second key component of the RiboMarker^®^ platform is the universal library preparation protocol T[1+2], compatible with all the pretreatment steps described above. This protocol is based on the original RealSeq platform technology for preparation of small RNA sequencing libraries (Barberán-Soler et al. 2018) that is commercially available as RealSeq^®^-Biofluids. The protocol T[1+2] workflow is schematically shown in supplementary Fig. S1, and its experimental details are described in MATERIALS and METHODS. A recent systematic evaluation of commercially available small RNA-Seq library preparation kits selected the RealSeq platform for further development of the small RNA sequencing pipeline for absolute quantitation based on the sensitivity, consistency of detection of miRNAs, low bias towards different miRNAs, as well as usability criteria (Khamina et al. 2022). This protocol can simultaneously capture RNA Types 1 and 2 (Shigematsu and Kirino 2022) which allows detection of a higher variety of specific small RNAs than the standard methods of small RNA-Seq library preparation (specific to RNA Type 1). The RiboMarker^®^ approach could also be combined with these standard methods using alternative pretreatment protocols suggested in supplementary Table S1.

To optimize and monitor the efficacy of conversion between the different RNA Types, we leveraged a pool of synthetic spike-in sRNAs corresponding to <u>s</u>pecific <u>T</u>ypes of <u>sRNAs</u> (stsRNAs). The stsRNA sequences are shown in Table 2. The stsRNAs feature (i) different lengths (20, 30, 40, 50, and 60 nt), (ii) 6-nt internal RNA Type-specific bar-codes, (iii) 4-nt randomized nucleotides at both ends, and (iv) different termini corresponding to sRNA Types 1 through 4 shown in Fig. 1. The core sequences of these stsRNAs (excluding the random and barcode sequences), which do not have homology to the human genome, were adapted from Locati et al. (2015). A placement of four randomized nucleotides on both the 5′ and 3′ ends of stsRNAs were adapted from Lutzmayer et al. (2017). Equimolar pools of these 20 (4 Types x 5 lengths) individual stsRNAs were spiked in the analyzed RNA samples at different inputs before pretreatments and/or preparation of sequencing libraries. We determined that the optimal inputs of the stsRNA pool were from 1 to 5% relative to total brain RNA samples, which allowed robust identification and quantification of stsRNAs along with the analysis of natural sRNA sequencing profiles.

**TABLE 2.**
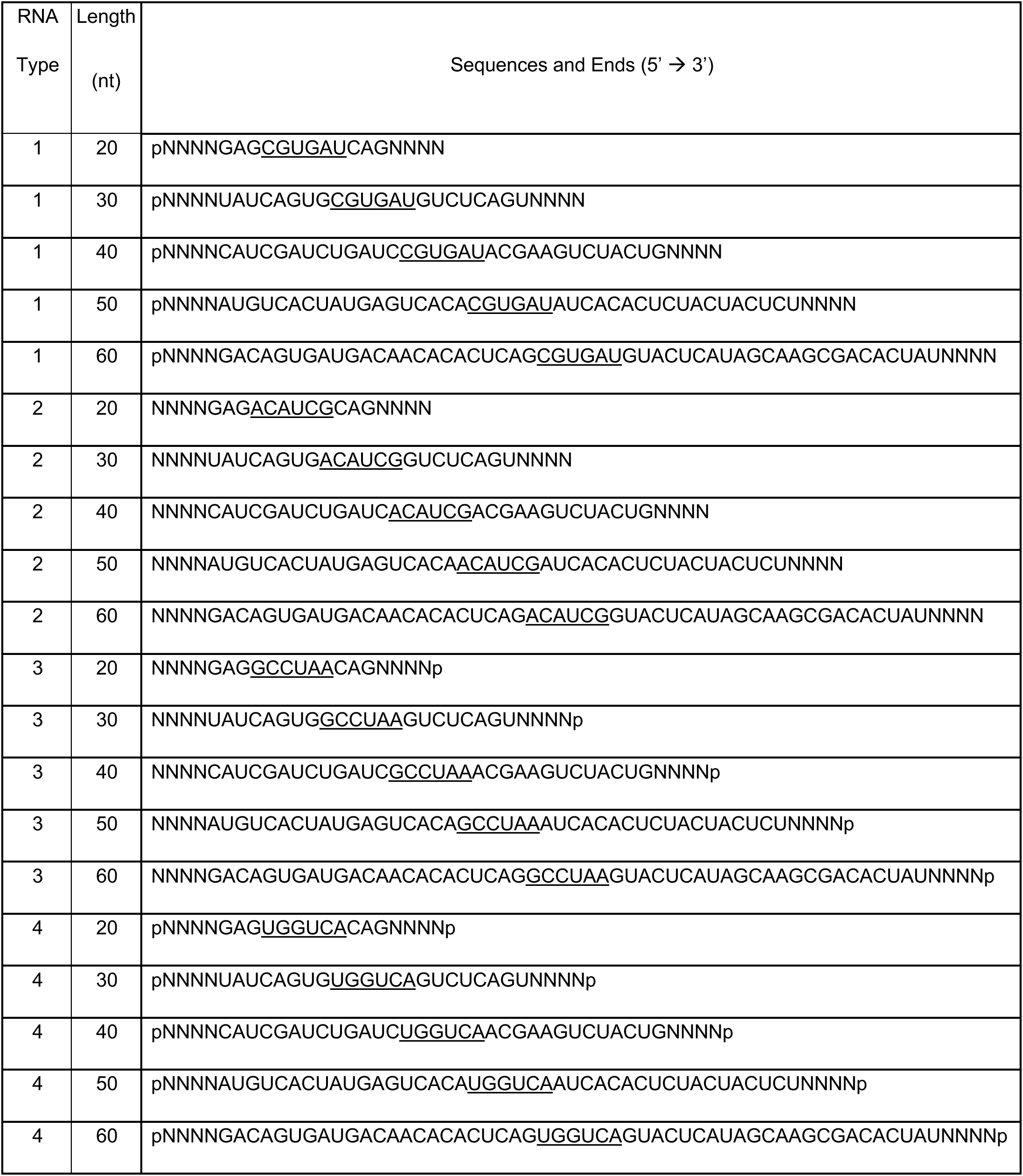
Sequences of spike-in RNAs (stsRNAs) having barcodes (underlined) specific to each RNA Type.

### Implementation of RNA circularization in the RiboMarker^®^ platform

The RiboMarker^®^ approach exploits circularization reactions catalyzed by an RNA ligase for two different purposes. First, it is applied in the sequencing library preparation protocol T[1+2] (supplementary Fig. S1), where sRNAs are first ligated to the sequencing <u>c</u>ombo <u>ad</u>apter (CAD) and then sRNA-CAD ligation products are circularized. In contrast to similar methods of sequencing library preparations based on circularization of cDNAs (Kwon 2011; Jackson et al. 2014; Heyer et al. 2015), our approach allows sequencing of full-length sRNAs rather than their RT cDNA products. The RNA circularization steps are also used in the pretreatment protocols (Table 1), where sRNAs are circularized to block the sRNA ends preventing adapter ligation and incorporation of “unwanted” RNA into the sequencing libraries.

Although rates of RNA *inter*molecular ligation reactions catalyzed by Rnl1 are strongly dependent on the identity of terminal nucleosides at 5’-P and 3’-OH ends involved in these reactions (England and Uhlenbeck 1978; Ohtsuka et al. 1980; Romaniuk et al. 1982), the RNA circularization (*intra*molecular ligation) reactions exhibit less bias to the identity of terminal nucleosides and a presence of ribose 2’-O-Methyl modification (2’-OMe) at the RNA 3’-end, depending mostly on RNA length and folding (Silber et al. 1972; Cranston et al. 1974; Kaufmann et al. 1974; Kumar et al. 2011; Del Piano et al. 2022). For example, maximal rates were obtained with poly(A) of an average chain length between 34–40 nt, while this rate was about twice that observed with substrates with an average chain length of 70–100 nt (Silber et al. 1972). In another study, the maximum circularization rate for (pA)_n_ was observed for (pA)_10-16_, but then it declined for (pA)_30_ and (pA)_100_ by ∼1.5- and 3-folds, respectively (Kaufmann et al. 1974). Moreover, there are reports describing efficient circularization of synthetic and naturally occurring RNAs of more than 300 nt long (Petkovic and Müller 2015; Del Piano et al. 2022; Bradley et al. 2024). Although the generation of concatenated ligation products through intermolecular reactions may also occur concurrently with RNA circularization (Hafner et al. 2011; Chu et al. 2015), the circularization reaction greatly predominates for RNAs shorter than 60 nt (Uhlenbeck and Gumport 1982; Harrison and Zimmerman 1984) and can be additionally promoted by dilution of the ligation reaction mixtures (Obi and Chen 2021). Similar tendencies were observed for sRNA-Seq library preparations. In comparison to the intermolecular ligation of adapters to 5’-p ends of sRNAs in dual-adapter ligation methods, which has significant biases (Hafner et al. 2011; McCormick et al. 2011), the single adapter and circularization-based approach used in the RealSeq platform (supplementary Fig. S1) significantly reduces this bias (Barberán-Soler et al. 2018). Furthermore, the possible concatemerization of sRNAs and/or sRNA-adapter ligation products are not an issue for the circularization-based methods of sRNA-seq library preparation (Chu et al. 2015; Gout et al. 2017; Barberán-Soler et al. 2018; Bradley et al. 2024).

### Evaluation of circularization-based RiboMarker^®^ sRNA sequencing library preparation method

We started development of the RiboMarker^®^ platform with an assessment of the universal protocol T[1+2] and its capability to capture stsRNAs of different lengths by analyzing sequencing length profiles (sequencing read count vs length of sequences) while the stsRNA Types were identified by sequences of stsRNA barcodes.

Separate stsRNA libraries were prepared by protocol T[1+2] for stsRNAs in the absence of naturally occurring RNAs (mock controls) (Fig. 2A, left panel and supplementary Fig. S2A) or for stsRNAs spiked into total RNA isolated from either reference human brain (Fig. 2B, left panel) or human plasma (Figs. 2C–D, left panels) samples. Our data demonstrated that protocol T[1+2] captured similar percentages of stsRNA Type 1 and Type 2, whereas only marginal percentages of stsRNA Type 3 and Type 4 were included in the sequencing libraries (Figs. 2A-D, left panels and supplementary Table S2). Triplicate sequencing length profiles demonstrated no significant variations between the ratios of the RNA Types for the stsRNAs of the same lengths when the stsRNAs were spiked either into human brain RNA samples (supplementary Fig. S3A) or into two groups (named here as H and D) of human plasma RNA samples (supplementary Fig. S4). However, we noticed two phenomena related to differences of the sRNA profiles among the tested samples. The first phenomenon was differing average ratios of RNA Type 1 and Type 2 for stsRNAs of all lengths in the mock controls performed in the absence of naturally occurring RNA or when they were spiked into total RNA from human brain (RNA Type 1 > RNA Type 2) and plasma (RNA Type 2 > RNA Type 1) samples (Figs. 2A–D, left panels and supplementary Table S2). In principle, a competition between naturally occurring sRNAs and synthetic stsRNAs of the same RNA Types present at different molecular ratios in the samples could explain the Types ratio phenomenon. Nevertheless, it does not explain variations of the ratios between RNA Type 1 and Type 2 in the stsRNAs of different lengths. The second phenomenon was related to the differential capture of the stsRNAs of different lengths spiked in human brain (20 ≥ 30 > 40 > 50 >> 60 nt) and plasma (40 > 20 ∼ 30 > 50 > 60 nt) RNA samples (Figs. 2B–D, left panels). Although the latter differences were minimal between the mock control and human brain samples, the differences in the ratios of stsRNA of different lengths between human brain and plasma RNA samples were pronounced. Also, we found that the average intracellular sRNAs in human brain samples (Fig. 2E left panel) were longer as compared to the extracellular sRNA molecules found in the human plasma RNA samples (Figs. 2F–G, left panels). Furthermore, we hypothesized that opportunistic folding and co-folding of stsRNAs having randomized ends with natural sRNAs, which is length, structure, and concentration dependent, may affect enzymatic reactions of RNA molecules at their ends as previously proposed (Zhuang et al. 2012a; Maguire et al. 2020). Regardless of the mechanism of these phenomena, protocol T[1+2] provides the capability for robust quantification of sRNAs ≤50 nt in human plasma samples, where 95% of extracellular sRNAs are less than 45 nt in length (Akat et al. 2019; Galvanin et al. 2019; Giraldez et al. 2019; Wang et al. 2024).

**FIGURE 2.**
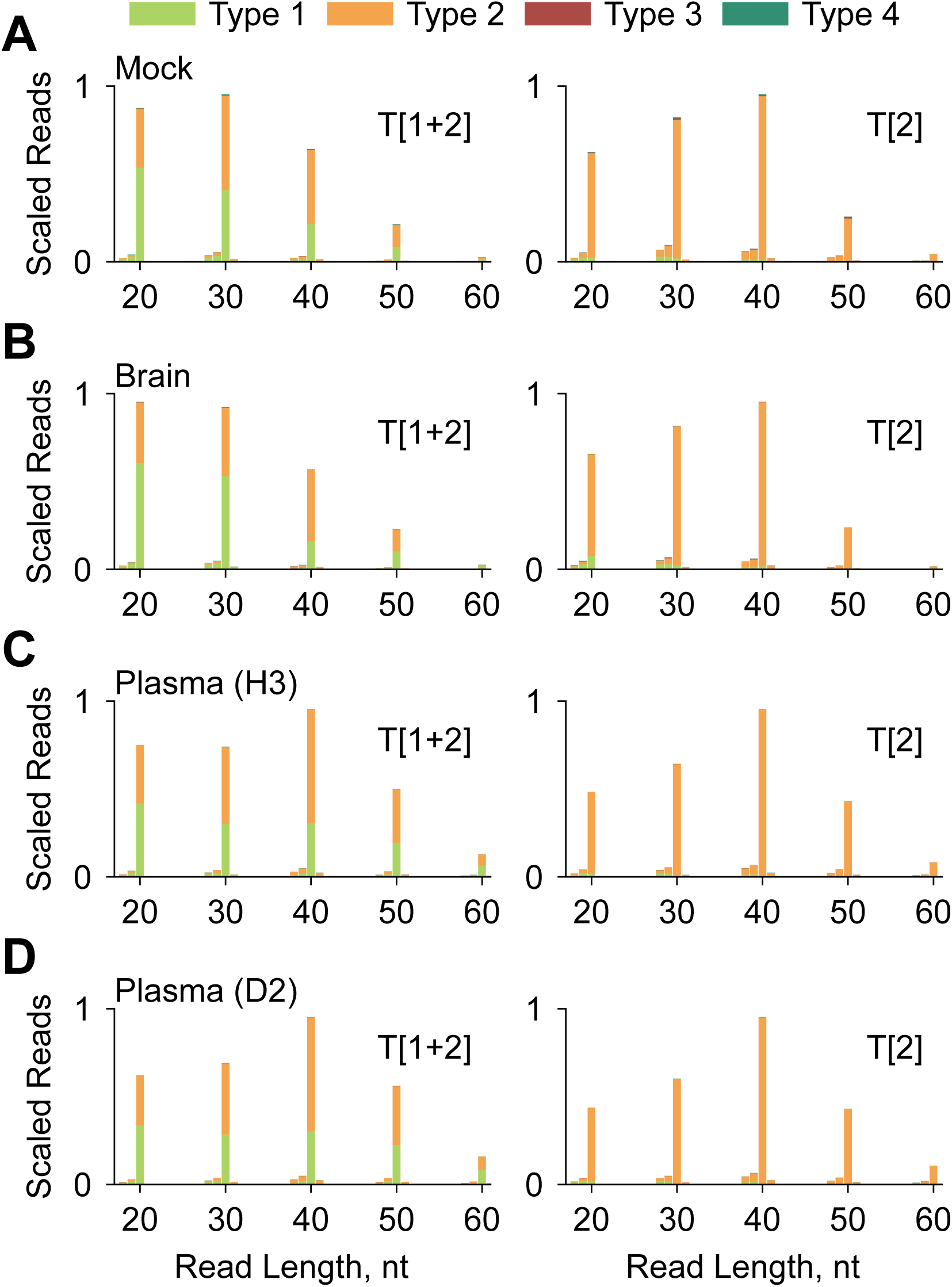
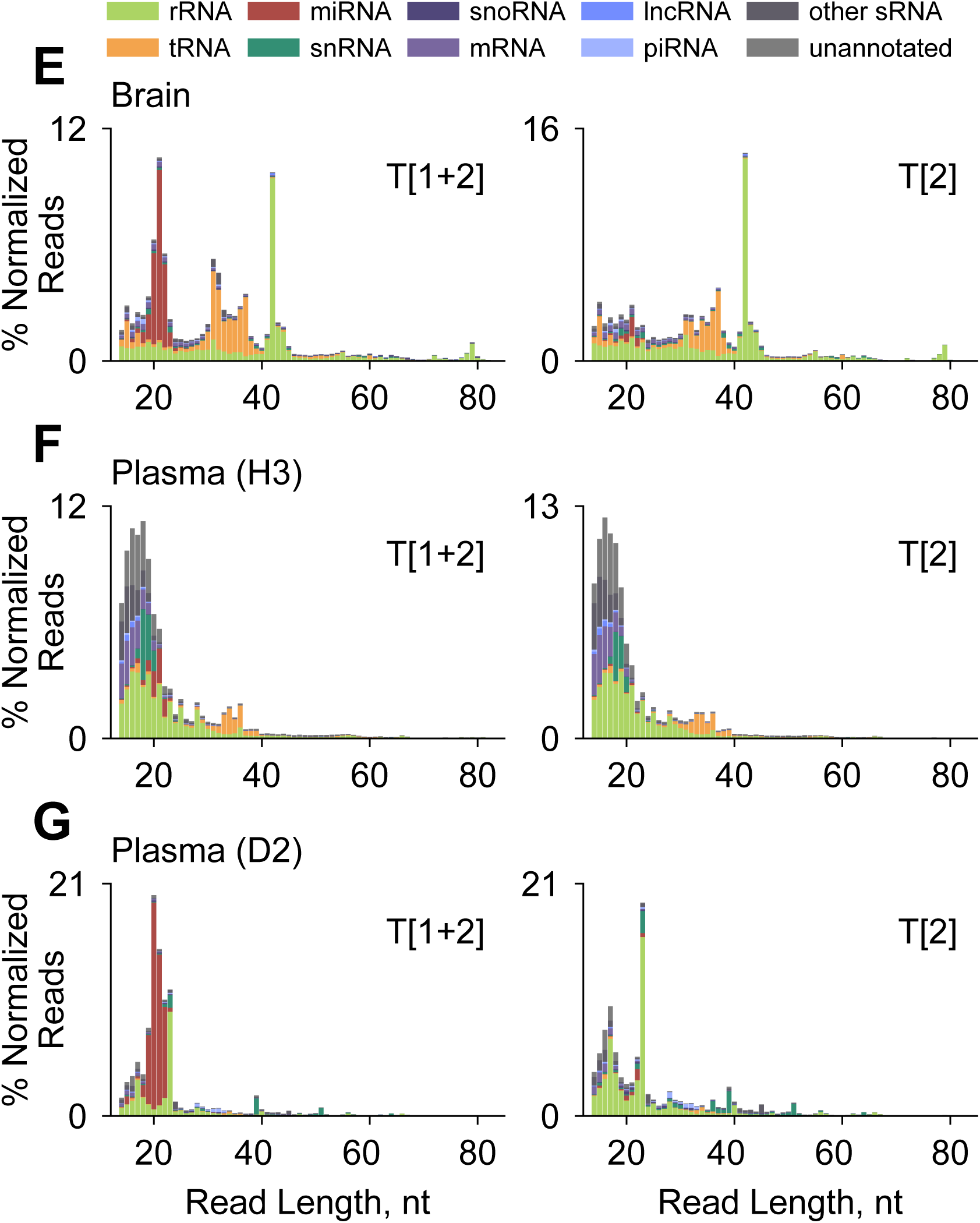
Sequencing length and RNA Type profiles for (**A**) stsRNAs in the absence of naturally occurring RNAs (mock controls); stsRNAs spiked in total RNA from (**B**) human brain and (**C**–**D**) indicated human plasma RNA samples; naturally occurring sRNAs detected along with stsRNAs spiked into the (**E**) human brain sample, or (**F**–**G**) indicated human plasma RNA samples for the libraries prepared by protocols T[1+2] (left panels) and T[2] (right panels).

### Circularization of small RNAs prevents their incorporation into sequencing libraries

sRNA circularization has been previously used for the detection of sRNAs by RT-qPCR (Kumar et al. 2011) and sequencing (Chu et al. 2015; Gout et al. 2017; Barberán-Soler et al. 2018; Bradley et al. 2024). In this study, we applied it for the exclusion of selected sRNA Types (Fig. 1) from sequencing libraries. Following the circularization of sRNA Type 1, the termini of selected non-circularized sRNA Types are converted into Types 1 and/or 2 in the next pretreatment step(s), and then these converted sRNAs are either also excluded by circularization or incorporated into sequencing libraries using the universal protocol. In doing so, the exclusion of highly abundant sRNA Types from the libraries allows for a more sensitive detection of rare sRNA of other Type(s). This new exclusion-by-circularization approach could also be used along with other methods of sequencing library preparation that rely on the ligation of adapter(s) to RNA end(s) or on 3’-end extension by a ribonucleotidyl transferase described elsewhere (Crocker et al. 2022; Shi et al. 2022; Shigematsu and Kirino 2022; Wang et al. 2024).

Among all pretreatment protocols (Table 1) featuring the RNA circularization step, protocol T[2] with circularization by RNA ligase, in which the abundance of RNA Type 1 is decreased while RNA Type 2 is increased, provides the best estimation of this efficiency. To find the most efficient conditions for the circularization step, we used the stsRNA pool spiked into human brain RNA samples at 37°C in standard T4 RNA ligase reaction buffer supplemented with 10% PEG 8000, 20 U Murine RNase Inhibitor (all NEB) and 100 µM ATP with the following variables: 10 U Rnl1 for 1 hour (Rnl1_1h); 20 U Rnl1 for 1 hour (2xRnl1_1h); 20 U Rnl1 for 2 hours (2xRnl1_2h); and 10 U Rnl1 + 10 U Rnl2 for 1 hour (Rnl1+Rnl2_1hr). The Rnl1+Rnl2 combination was tested because of the ability of both Rnl1 and Rnl2 to circularize RNA Type 1 (Ho and Shuman 2002; Ho et al. 2004; Yin et al. 2004), and a potential synergy between these two enzymes having different structural and end-nucleotide biases for ligation of co-folded RNA ends (Yoshinari et al. 2017; Chen et al. 2020). Sequencing analysis of these different versions of protocol T[2] for the efficiency of RNA Type 2 enrichment revealed that the “Rnl1+Rnl2_1h” reaction condition provided 90.91±0.19% yield of stsRNA Type 2 for the human brain RNA samples and 93.50±0.51% for the H and 93.90±0.24% for the D human plasma RNA samples (supplementary Table S3). The corresponding stsRNA length profiles demonstrated high reproducibility for the human brain (supplementary Fig. S3B) and plasma (supplementary Fig. S5) RNA samples. The mock controls performed in the absence of naturally occurring RNA resulted in a 91.53±0.23% yield of stsRNA Type 2 (supplementary Table S4).

Although two other circularization reaction conditions “Rnl1_1h” and “2xRnl1_1h” provided only marginally smaller yields (90.59±0.28% and 89.71±0.02%, respectively) of stsRNA Type 2 in the brain RNA samples in comparison to the selected condition, these conditions also produced higher percentages of side products of stsRNA Type 3 and Type 4 than the “Rnl1+Rnl2_1h” condition (supplementary Table S3 and supplementary Fig. S6). The most prominent amounts of stsRNA Type 3 products (8.25%±0.03%) were produced under the “2xRnl1_2h” conditions (supplementary Table S3). We concluded that the generated stsRNA Types 3 and 4 were products of the T4 RNA ligase reverse reaction, causing removal of single mononucleotide diphosphates (5’-P-N-3’-P or 5’-P-N-2’,3’>P) from the 3’-end-phosphorylated RNA molecules that have been previously described only for Rnl1 (Krug and Uhlenbeck 1982; Uhlenbeck and Gumport 1982). The sequencing length profiles confirmed that this side reaction indeed removes a single terminal nucleotide from 3’ ends of stsRNA Types 3 and 4 that makes them shorter by 1 nt and converts them to stsRNA Types 2 and 1, respectively, allowing their incorporation into sequencing libraries (supplementary Fig. S6). The higher percentage of sequencing reads for stsRNA Type 3 (converted by Rnl1 to a 1 nt shorter Type 2) in comparison to stsRNA Type 4 (converted to a 1 nt shorter Type 1) is because the converted stsRNA Type 1 can then be circularized by Rnl1 and, therefore, excluded from the library.

Based on the above results, we selected the “Rnl1+Rnl2_1h” reaction condition to be used for the RNA Type 1 circularization in protocols T[2], T[3+4], T[3] and [T4]. In contrast to protocol T[1+2], the stsRNA sequencing length profiles for protocol T[2] were similar in the tested samples, including the mock controls of stsRNAs performed in the absence of naturally occurring RNA (Fig. 2A, right panel and supplementary Fig. S2B), human brain (Fig. 2B, right panel) and plasma RNA samples (Fig. 2C–D, right panels). Furthermore, we analyzed the stsRNA Type distributions for the individual length groups of 20-, 30-, 40-, 50- and 60-nt stsRNAs spiked into three different human H (H1, H2, H3) plasma RNA samples for protocol T[1+2] (supplementary Fig. S4A and Table S5) and protocol T[2] (supplementary Fig. S5A and Table S6). As indicated in supplementary Table S6, the average percentage of stsRNA Type 2 spiked into the human H plasma RNA samples were found to slightly increase with increasing stsRNA lengths from 88.48±1.32% (20-nt) to 92.78±0.43% (30-nt) and 96.42±0.22% (40-nt), held at 95.21±0.44% (50-nt), and then dropped to 86.39±1.00% (60-nt). By analyzing the percentage of stsRNA Type 1 at the different lengths in the human H plasma RNA samples using protocols T[1+2] (supplementary Table S5) and T[2] (supplementary Table S6), we were able to evaluate both the extent of stsRNA Type 1 reduction from sequencing libraries and the corresponding efficiency of their circularization. We calculated the reduction for each stsRNA length for the human H plasma samples by subtracting the corresponding stsRNA Type 1 percentages found with protocol T[2] from those with protocol T[1+2] and then dividing the differences by the Type 1 percentages for protocol T[1+2] as shown in supplementary Table S7. These data revealed a reduction of stsRNA Type 1 of about: 79.85% (20-nt), 83.30% (30-nt), 89.05% (40-nt), 87.90% (50-nt), and 74.02% (60-nt). These results indicated that we could achieve high efficiency of stsRNA Type 1 reduction through circularization regardless of stsRNA lengths under the selected circularization reaction conditions. The efficiency of the sRNA circularization reaction could be further increased to get closer to 100% by adjusting the reaction conditions considering that Rnl1 and Rnl2 have different optimal concentrations of Mg^2+^ and reaction buffer pH (Ho et al. 2004; Yin et al. 2004; Viollet et al. 2011).

### Detection of naturally occurring RNA Types 1 and 2

After our assessment of the synthetic stsRNAs, we analyzed sequencing length profiles for the major RNA classes of naturally occurring sRNAs in the same human brain and plasma RNA samples processed using protocols T[1+2] and T[2]. For the human brain RNA samples, a comparison of (a) the sRNA profiles of major RNA classes generated using either protocol T[1+2] (Fig. 2E, left panel) or protocol T[2] (Fig. 2E, right panel) confirmed the efficiency of depletion of sRNA Type 1 as shown by the reduction of the miRNA fraction in the protocol T[2] profiles. A subtraction of the normalized sequencing length profiles for the same natural sRNAs samples processed by protocol T[2], which is specific for RNA Type 2, from the profile for protocol T[1+2], which is specific for RNA Types 1 and 2, could enable the identification of sequencing length profiles for the depleted sRNA Type 1 that are detected directly by sequencing of the standard sRNA-Seq libraries. For example, using this approach, we found that most of the sRNA Type 1 in the human brain sample were represented by miRNAs as well as fragments of tRNAs and rRNAs (supplementary Fig. S7). In comparison to the human brain sRNA profiles (Fig. 2E), the corresponding human plasma sRNA profiles were comprised of a larger fraction of ultrashort fragments (< 20 nt) and had more sRNAs derived from rRNA for both protocols T[1+2] and T[2] (Figs. 2F–G). For protocol T[1+2], we also noticed significant biological variations of sRNA sequencing length profiles among the different H and D human plasma samples, especially in the tRNA and miRNA classes (supplementary Fig. S8). A comparison of the length profiles for protocols T[1+2] and T[2] revealed that generally sRNA Type 2 are significantly longer than the sRNA Type 1 molecules for both the human brain (Fig. 2E) and, especially, in the human plasma (Figs. 2F–G) samples, where on average about 93% of sRNA reads in protocol T[1+2] sequencing libraries can be classified as sRNA Type 2.

Sequencing length profiles for sRNA Type 1 using standard library preparation methods have been previously described for both cellular (Kawaji et al. 2008; Jackowiak et al. 2017; Shi et al. 2021b) and plasma (Max et al. 2018; Wang et al. 2024) RNA samples. To the best of our knowledge, sRNA Type 2 sequencing profiles have not been previously analyzed. Here we present the representative sequencing profiles (selected from corresponding replicates) for sRNA Type 2 of all major sRNA classes detected in human brain (Fig. 3) or plasma (Fig. 4) RNA samples by protocol T[2]. Looking at these same sRNA classes between the human brain and plasma RNA samples confirmed that plasma sRNA profiles comprised a larger portion of shorter fragments than the brain profiles. While length profiles for tRNA and mRNA-derived sRNAs were relatively consistent across the representative H and D human plasma samples (Figs. 4B and 4E), the profiles for rRNA, snRNA, snoRNA, and lncRNA-derived sRNAs showed significant differences with the appearance of distinctive longer RNA fragments (Figs. 4A, 4C, 4D, and 4F, respectively). Additionally, we noted significant variations in the percentage of sequencing reads for these RNA classes between the H and D human plasma samples.

**FIGURE 3.**
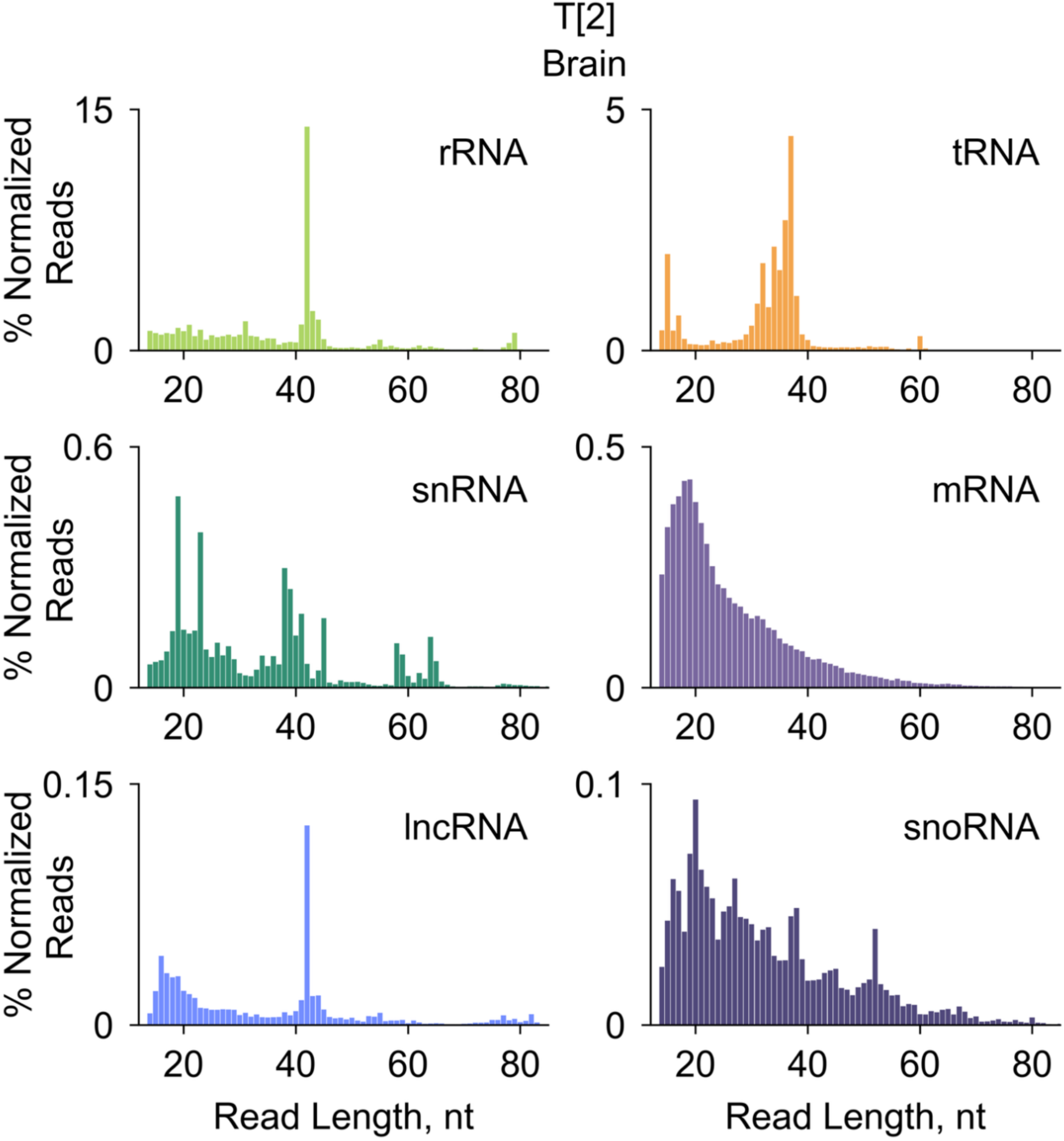
Representative sequencing length profiles for indicated RNA classes of naturally occurring sRNAs found in a human brain RNA sample for libraries prepared by protocol T[2].

**FIGURE 4.**
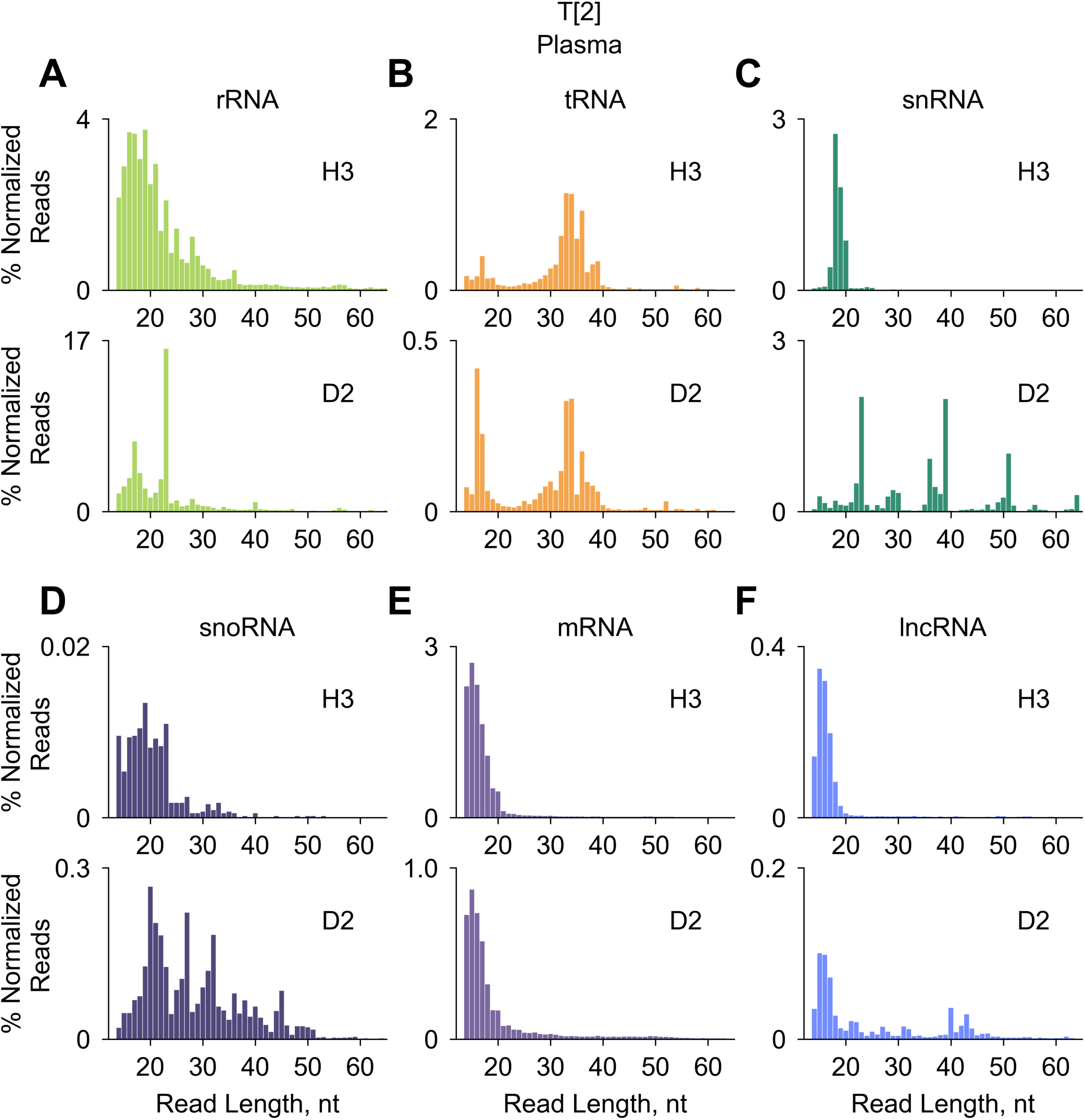
Comparison of sequencing length profiles for naturally occurring sRNAs from representative human H (H3) and D (D2) plasma RNA samples between libraries prepared by protocol T[2] for the indicated RNA classes.

Furthermore, protocol T[2] revealed intriguing features of piRNA-related sequences. Like miRNAs, mature (processed) piRNAs could be classified as RNA Type 1; however, these contain a 2’-OMe modification at their 3’-end. It has been previously shown that this modification could reduce the circularization rate of sRNAs having 5’-P and 2’-OMe/3’-OH ends by Rnl1 (Kumar et al. 2011). Using sequencing data for technical triplicates from the human brain RNA samples, we calculated that protocol T[2] reduced the percentage of the miRNA-related reads in comparison to protocol T[1+2] by 11.24±2.80 fold, whereas percentages of piRNA-related reads were reduced by only 1.30±0.05 fold (supplementary Tables S8A–B). In similar experiments using the human H plasma samples, the observed reduction for miRNA- and piRNA-related reads were 27.38±7.50 and 1.17±0.05 fold, respectively (supplementary Table S8C–D). Based on these experiments, it would be reasonable to suggest that the 2’-OMe modification inhibited circularization of piRNA-related sRNAs in protocol T[2] and, therefore, prevented their depletion from sequencing libraries. However, we also found protocol-specific differences in sequencing length profiles of piRNA-related sRNAs between all other RiboMarker^®^ protocols (as described below) in human plasma samples that implied the presence of sRNA Types 2, 3 and 4 in piRNA-related sequencing reads. The latter results agree with previously suggested piRNA biogenesis pathway information that indicates these RNA Types are generated from precursor piRNAs (Shigematsu et al. 2021).

### Detection of all four stsRNA Types after PNK pretreatment

PNK is commonly used to erase the differences between the phosphorylation states of sRNA ends by converting them simultaneously to RNA Type 1, followed by standard sRNA-Seq library preparation methods (Akat et al. 2019; Giraldez et al. 2019; Crocker et al. 2022; Shi et al. 2022, Shigematsu and Kirino 2022; Solaguren-Beascoa et al. 2023). This approach, also known as Phospho-RNA-Seq, uses the standard, one-step PNK treatment of RNA performed in Tris-HCl buffer at pH 7.6 in the presence of 1 mM ATP under conditions adopted from earlier studies (Pan et al. 1991; Busch et al. 2000; Schürer et al. 2002; Brown and Bevilacqua 2005). Although these reaction conditions are optimal for the phosphorylation of 5’-OH ends of RNA Types 2 and 3 (Cameron and Uhlenbeck 1977), the maximum rate of dephosphorylation of 3’-P and 2’, 3’>P ends of RNA Types 3 and 4 is ∼pH 5.9, whereas at pH 7.6 it decreases by ∼10-fold (Cameron and Uhlenbeck 1977; Zhuang et al. 2012b). Also, PNK has significant 5’-P reverse (dephosphorylation and exchange) activities catalyzed by ATP (van de Sande et al.1973). Additional anticipated problems include known bias of PNK to preferentially phosphorylate nucleic acids with certain 5’- and 3’-end nucleotides, sequences, lengths, and secondary structures (Székely and Sanger 1969; Lillehaug and Kleppe, 1975; van Houten et al. 1998; Lee et al. 2013).

In this study, we sought to optimize desirable and minimize unwanted PNK activities through an examination of the effects of different PNK pretreatment conditions on the sequencing output of our pool of stsRNA spiked into human brain RNA samples. In contrast to the Phospho-RNA-Seq method, we aimed to perform the 3’-end dephosphorylation and 5’-end phosphorylation by PNK in two separate steps (rather than simultaneously) under the optimal conditions for each step. In the first step, 3’-dephosphorylation was performed as an RNA pretreatment as described in protocol T[1+2+3+4] (Table 1). For this purpose, we tested three buffer conditions adapted from a study by Cameron and Uhlenbeck (1977), including 100 mM Tris-HCl at pH 7.6 in the presence of 1 mM ATP; and 100 mM Imidazole-HCl at pH 6.5 or 100 mM MES-NaOH at pH 6.0 (both in the absence of ATP) along with 10mM MgCl_2_, and other common buffer components and reaction conditions described in MATERIALS and METHODS. In the second step, the 5’-phosphorylation of RNA Types 2 and 3 was carried out in Step 2b of the library preparation by PNK in standard NEB kinase reaction buffer at pH 7.6 in the presence of 100 µM ATP which minimized the 5’-end dephosphorylation of RNA Types 1 and 4 (supplementary Fig. S1A). Although Cameron and Uhlenbeck (1977) demonstrated that the rate of 3’-P dephosphorylation depends on the buffer pH, we did not observe significant differences among the RNA Type distributions in our sequencing libraries prepared for stsRNAs spiked in different naturally occurring RNA samples, which were treated under the described above 3’-end dephosphorylation conditions (supplementary Table S9). Despite the different stsRNA Types having been pre-mixed at an equimolar ratio of 25% per Type, we found that protocol T[1+2+3+4] applied to stsRNAs spiked into either human brain or plasma RNA samples did not produce the expected equal distribution for the stsRNA Types (supplementary Table S9). While the percentage of stsRNA Types 1 and 2 were slightly overrepresented (relative to the expected 25%), both Type 4 and (especially) Type 3 were underrepresented (supplementary Table S9).

For this study, we selected the PNK pretreatment using MES buffer for two reasons. The first reason was the slightly higher percentage of stsRNA Type 3 molecules included as compared to the other tested conditions (supplementary Table S9). Since stsRNA Types 3 and 4 both contain 3’-P ends, we hypothesize that the de-phosphorylation of 3’-P ends might be incomplete. Therefore, we compared the output of reactions containing either 10 or 20 Units of PNK for the same human plasma samples and found that the use of 2-fold increase of PNK concentration did not increase the percentage of stsRNA Type 3 inclusion (supplementary Table S9). Since the stsRNA Type 3 underrepresentation was also found with the mock stsRNA experiment under the selected PNK pretreatment conditions while the other three stsRNA Types were evenly represented (supplementary Table S9), we considered it as a minor bias specific to the composition of the stsRNAs and RiboMarker^®^ protocols. The second reason was related to a consideration between using either heat-inactivation of PNK or column purification to remove PNK prior to sequencing library preparation. It has been established that the heating of RNA molecules in select buffers can lead to their fragmentation, which is heavily influenced by RNA length, sequence, presence of divalent cations, buffer composition and pH (Chheda et al. 2024). In the MES buffer at pH 6.0, RNA molecules were expected to be more resistant to cleavage than in Tris buffer at pH 7.6 (Chheda et al. 2024). Also, both Imidazole and Tris can catalyze RNA cleavage in the presence of Mg^2+^ cations (Breslow and Huang 1991; AbouHaidar and Ivanov 1999).

To test whether the column-based removal of PNK or heat-inactivation in solution affected the resulting sequencing length profiles, we performed two sets of experiments using our human plasma samples in triplicate. In the first set of experiments, we either passed untreated stsRNAs spiked into human plasma RNA samples through the Zymo RNA clean and concentrate columns or used these as input for library preparation without a column clean-up (supplementary Fig. S9). We found that passing the sample through the column (supplementary Fig. S9B) did not significantly alter the resulting stsRNA length profiles as compared to the samples not passed through the columns (supplementary Fig. S9A). In the second set of experiments, we treated the stsRNA with PNK in the MES pH 6.0 buffer, and then PNK was either heat-inactivated at 65°C for 20 min in the presence of 20 mM EDTA pH 8.0 or removed by the columns without heating. In this case, we observed drastic differences in the stsRNA profiles for these two experiments (supplementary Fig. S10). The heat-inactivation of PNK resulted in less robust capture of longer stsRNAs, with a significant increase in the relative abundance of shorter molecules (supplementary Fig. S10A) as compared to the column-purified stsRNAs (supplementary Fig. S10B). Given our previous observation that passing through the columns does not alter the inclusion ratio of different lengths and, therefore, does not artificially enrich for longer stsRNAs (supplementary Fig. S9), we concluded that the heat-inactivation of PNK leads to the partial fragmentation of our stsRNAs that evidently affects longer molecules more readily. Additionally, heat-inactivation seemed to lead to a reduced inclusion (13.23±0.42%) of stsRNA Type 3 of all lengths as compared to the column-purified samples (19.29±0.43%) (supplementary Fig. S10).

We evaluated the selected protocol T[1+2+3+4] using stsRNA mock controls and stsRNAs spiked into human brain and plasma RNA samples by analyzing the distribution of stsRNA Types in corresponding sequencing length profiles. In this evaluation, we also included a comparison of the sequencing outputs between protocol T[1+2+3+4] and Phospho-RNA-Seq. Phospho-RNA-Seq includes a PNK pretreatment in buffer comprising 10 mM MgCl_2_ and Tris-HCl pH 7.6 followed by NEBNext Small RNA library preparation, which specifically detect RNA Type 1 (Giraldez et al. 2019; Solaguren-Beascoa et al. 2023). For the later comparison, we spiked stsRNAs into a human plasma sample (selected from the CL cohort) and skipped the PNK heat-inactivation step, which is custom for the Phospho-RNA-Seq, but kept the column cleanup step which was common for protocol T[1+2+3+4] and Phospho-RNA-Seq.

A comparison of stsRNA sequencing length profiles for the mock controls and stsRNAs spiked into the human brain and different human plasma RNA samples (Figs. 5A-E) indicates that, similarly as for protocols T[1+2] and T[2], the natural sample RNA compositions can significantly affect the ratios between stsRNA Types for protocol T[1+2+3+4]. Furthermore, the relative capture of stsRNAs of different lengths were also affected. Although these phenomena are not fully understood, the putative competition and/or co-folding between naturally occurring sRNAs (whose contents can vary among different samples) and synthetic stsRNAs might be responsible for these effects. Besides obvious differences between the mock control (lacking a naturally occurring sRNA background) and other sample types, the human brain and plasma samples (and human plasma samples from different sources) differ in abundance of naturally occurring sRNA Types and lengths.

**FIGURE 5.**
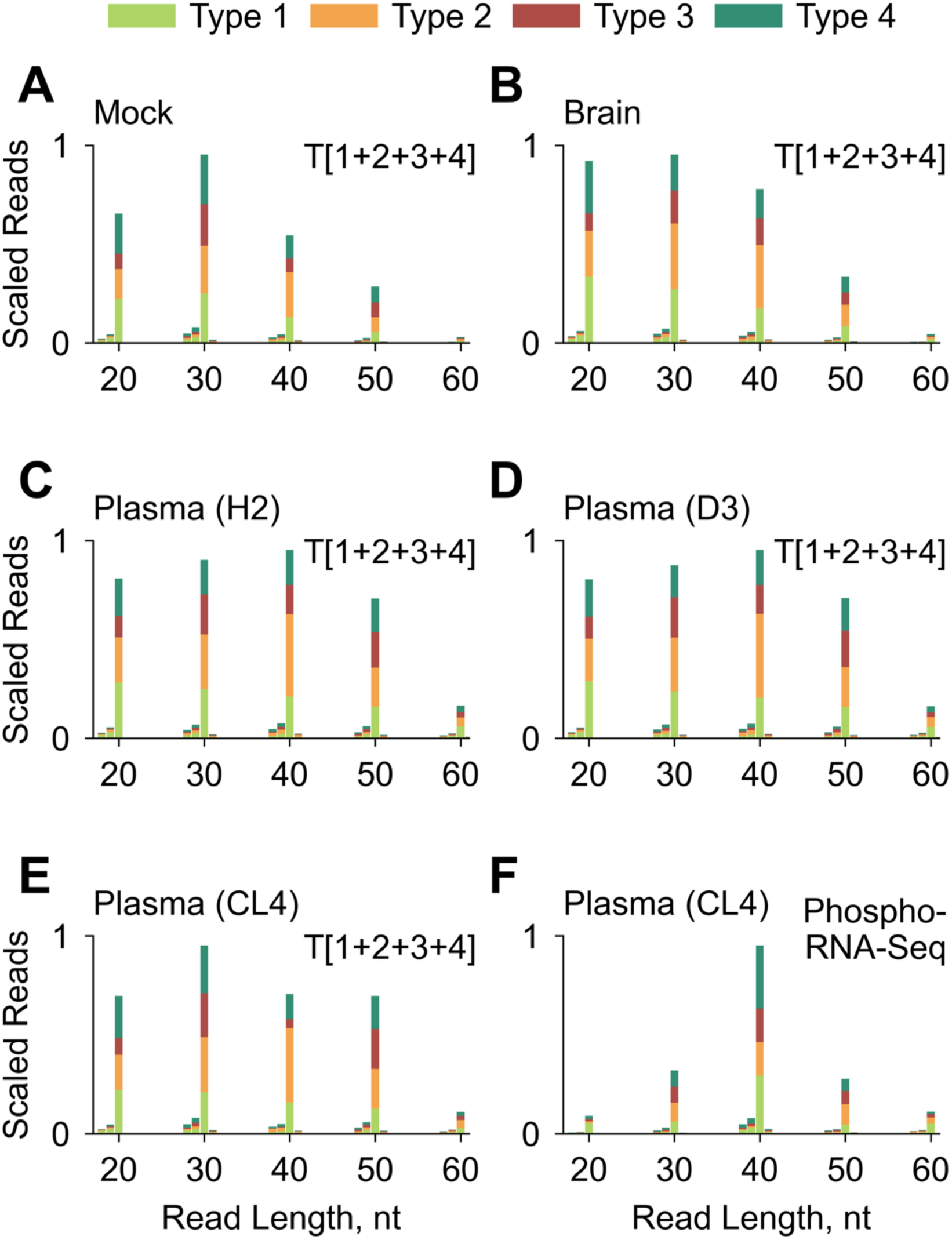
Representative sequencing length and RNA Type profiles for different stsRNAs contexts, including: (**A**) mock control lacking a naturally occurring sRNA background; and stsRNAs spiked into either (**B**) human brain RNA or (**C** – **F**) indicated human plasma RNA samples. The libraries were prepared either by (**A** – **E**) RiboMarker^®^ protocol T[1+2+3+4] or (**F**) Phospho-RNA-Seq.

Looking more deeply at the RNA Type distribution for stsRNAs of different lengths spiked in the human CL (CL1, CL2, CL4) plasma RNA samples for libraries prepared by protocol T[1+2+3+4] as compared to Phospho-RNA-Seq, revealed that they are different in two aspects. The first aspect is related to deviations from the intended 25% distribution among the four RNA Types at different lengths. On average for all stsRNA lengths, protocol T[1+2+3+4] includes more stsRNA Type 2 (35.15±1.51%) and less stsRNA Type 3 (15.8±1.75%) than Phospho-RNA-Seq (23.35±0.56% and 19.02±0.32%, respectively), whereas Phospho-RNA-Seq includes more stsRNA Type 1 (29.84±0.44%) and Type 4 (27.79±0.49%) than protocol T[1+2+3+4] (26.56±1.86% and 22.49±1.52%, respectively) as detected in experiments with stsRNAs spiked in three human plasma RNA samples (supplementary Tables S10 and S11). However, there were significant variations from the average stsRNA Type distributions of different lengths for both protocol T[1+2+3+4] and Phospho-RNA-Seq (supplementary Tables S10 and S11). The second aspect is related to the significant differences among the percent yield of stsRNAs of different lengths. Protocol T[1+2+3+4] provided almost even coverage of stsRNAs in the 20- to 50-nt range, with a slightly higher yield for 30-nt and a significantly lower yield for 60-nt species (Fig. 5E and supplementary Fig. S11A). In contrast, Phospho-RNA-Seq revealed a biased distribution of stsRNAs of different lengths, with a strong peak at 40-nt (10x yield) symmetrically surrounded by 30-nt and 50-nt (3x yield), and by 20-nt and 60-nt (1x yield) (Fig. 5F and supplementary Fig. S11B). The sequencing library preparation for Phospho-RNA-Seq with the NEBNext Small RNA library preparation kit for Illumina used a two-adapter ligation to RNA ends and (theoretically) should provide length-independent incorporation of the stsRNAs in the libraries. Even so, a co-folding between sRNAs and the adapters has been identified as a key determinant of ligation efficiency bias (Zhuang et al. 2012a; Maguire et al. 2020). A plausible explanation of this observed NEBNext bias would be an accidental co-folding between our stsRNAs and NEBNext sequencing adapter(s) that is either favorable or unfavorable to their ligation efficiency. However, profiling of stsRNAs (which in this study was primarily used to evaluate efficiency of conversions of their ends between different RNA Types) represents only an approximate outline of profiling naturally occurring sRNAs that might have different abundances, lengths and non-random nucleotides at their ends than the stsRNA molecules.

### Detection of all naturally occurring RNA Types simultaneously

After evaluation of protocol T[1+2+3+4] using the stsRNAs, we explored naturally occurring sRNAs present in the human brain and various plasma RNA samples. Representative sequencing length and RNA Type profiles, which were selected from the corresponding technical triplicates for human brain samples (supplementary Fig. S12) or biological triplicates for human H and D plasma samples (supplementary Fig. S13), are shown for all main sRNA classes in Fig. 6. The most distinctive differences between protocols T[1+2] and T[1+2+3+4] were increased rRNA and decreased miRNA contents (as indicated) for both the human brain (Fig. 6A) and plasma (Figs. 6B–C) samples. The observed differences in the rRNA and miRNA content in human plasma samples were more significant than in the human brain samples. The most striking was the complete domination of rRNA fragments in the human plasma protocol T[1+2+3+4] sequencing profiles. Still, these differences in the rRNA and miRNA content in these profiles between protocols T[1+2] and T[1+2+3+4] were less dramatic than differences between standard sRNA-Seq protocols detecting only sRNA Type 1 and Phospho-RNA-Seq detecting all four RNA Types simultaneously (Akat et al. 2019; Galvanin et al. 2019; Giraldez et al. 2019; Shi et al. 2021b; Solaguren-Beascoa et al. 2023).

**FIGURE 6.**
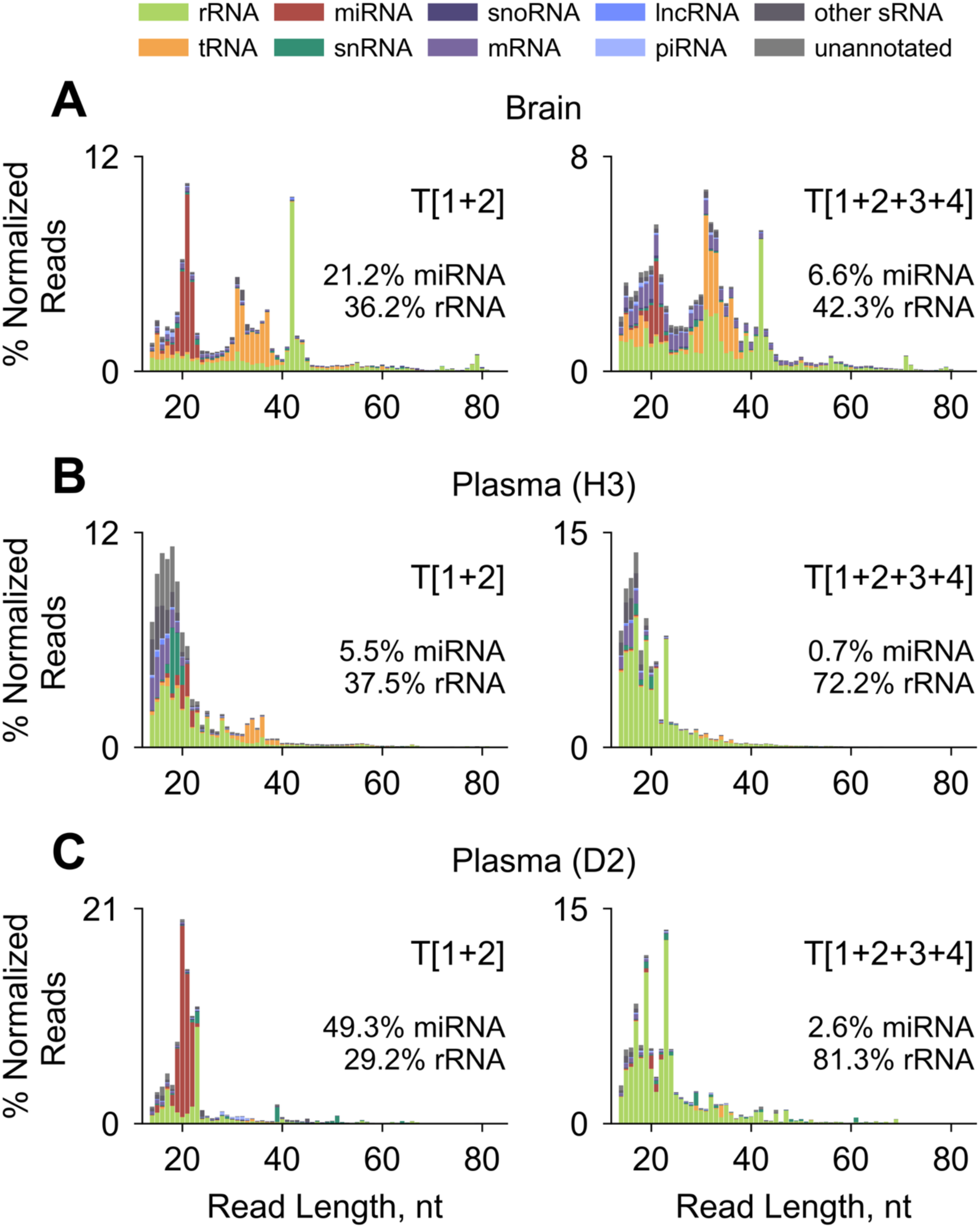
Representative sequencing length and RNA Type profiles for indicated RNA classes of naturally occurring sRNAs found in either (**A**) human brain or in (**B**–**C**) indicated human plasma RNA samples for libraries prepared by protocols T[1+2] (left panels) and T[1+2+3+4] (right panels).

In contrast to protocol T[2], which depletes RNA Type 1 (including miRNAs) from the sequencing libraries, protocol T[1+2+3+4] dilutes these reads by adding previously “hidden” sRNA species of RNA Types 3 and 4 to the libraries after the pretreatment. If the 5’-P ends of miRNAs are not significantly affected by the PNK pretreatment, the reduction in the capture of miRNAs will be inversely proportional to the increase of new sRNA species. A comparison between these two protocols showed that the average inclusion into the resulting libraries for miRNA was reduced by 3.34±0.09 fold in the human brain (supplementary Fig. S14A), 8.08±1.73 fold in the human H plasma, and 20.64±9.93 fold in the human D plasma samples (supplementary Fig. S14B). These results indicate that the “hidden” sRNA species in human D plasma samples can exceed 95% of total sRNA. Also, supplementary Figs. S14A-B show the changes in the abundance of other RNA classes, including tRNA, snRNA and snoRNA. These data indicate a high reproducibility for technical repeats regarding human brain samples and significant biological variation among the profiles for major RNA classes across different human plasma samples. In the human brain samples (supplementary Fig. S14A), the percentage of sequencing reads for sRNAs derived from mRNA showed the largest deviations between protocols T[2] and T[1+2+3+4], along with miRNAs (although in opposite directions), whereas sRNAs derived from the other RNA classes showed little or no change. These differences were largely related to a bigger proportion of mRNA fragments of Types 3 and 4 than Types 1 and 2, and vice versa for miRNA. The no or little change of the percentage of reads of other sRNAs suggested that a reduction of sRNA Types 1 and 2 was compensated by the appearance of an equivalent inclusion of sRNAs of Types 3 and 4. Despite the expectation that most of the reads which align to rRNA, mRNA, and lncRNA should be sRNA Type 3 due to the abundance of superfamily A RNases in human blood (Giraldez et al. 2019; Lee et al. 2019), we found that protocol T[1+2] detected almost as many of these RNA sequences as protocol T[1+2+3+4] both in the human brain (supplementary Fig. S14A) and plasma samples (supplementary Fig. S14B). This finding correlates with previous TGIRT-seq analyses, indicating that many of the mRNA fragments present in plasma have 3’-OH rather than 3’-P ends (Yao et al. 2020).

Subsequent analysis of the changes in the length profiles of the major RNA classes obtained by protocols T[1+2] and T[1+2+3+4] for sRNAs featured in Fig. 6 revealed more details. For the human brain samples (supplementary Fig. S15), we detected comparatively long (>30 nt) fragments of snRNAs, snoRNAs, and lncRNAs whose relative abundance were decreased in protocol T[1+2+3+4] as compared to protocol T[1+2], except for mRNA, where the abundance of longer fragments was moderately increased. An increase of short fragments (<30 nt) was the most noticeable for the snoRNA and lncRNA-derived sRNAs obtained by protocol T[1+2+3+4]. For a subset of the representative human H (H3) plasma sample (supplementary Fig. S16), nearly all sRNAs representing the main RNA classes were more fragmented (shorter in length) than in the human brain samples (supplementary Fig. S15), with rRNA and snRNA having ultrashort fragments (<20 nt). Also, we noticed two distinctive differences between these two protocols for the representative human D (D2) plasma RNA sample (supplementary Fig. S17). Firstly, longer tRNA fragments were captured in protocol T[1+2+3+4], suggesting their origin as RNA Types 3 and 4. Secondly, we detected longer distinctive fragments (>30 nt) of snRNA, snoRNA, and lncRNA in profiles for protocol T[1+2] than in profiles for protocol T[1+2+3+4], suggesting they are RNA Types 1 or 2.

In the RiboMarker^®^ approach, protocol T[1+2+3+4] is also incorporated into the workflows for protocols T[3+4], T[3], and T[4] (Table 1). Yet, it harbors the capacity to detect different sRNA Types simultaneously and could be used independently like Phospho-RNA-Seq. Using the latter protocol as a benchmark, we compared its capacity to capture sRNAs of different RNA classes with protocol T[1+2+3+4]. Although the corresponding sequencing length profiles of all sRNA classes performed in triplicate were in general similar (supplementary Figs. S18A–B), the profiles of certain individual sRNA classes for a representative human CL (CL4) plasma sample (selected from the biological triplicates) were different in fine details between protocol T[1+2+3+4] and Phospho-RNA-Seq (supplementary Fig. S19A–B). We then compared the percentage of sRNA sequencing reads uniquely mapped to the major RNA classes individually as captured by each protocol in the human CL plasma RNA samples (supplementary Fig. S20). Protocol T[1+2+3+4] provided the detection of more sRNA mapped to miRNA, piRNA and tRNA sequences, whereas Phospho-RNA-Seq detected slightly more mRNA and lncRNA fragments. Both protocols were similar in the detection of sRNAs mapped to rRNA, snoRNA and snRNA.

In essence, both protocol T[1+2+3+4] and Phospho-RNA-Seq have their strengths and weaknesses as standalone methods for the simultaneous detection of all sRNA Types. In theory, these methods should allow detection of higher diversity of sRNAs of all RNA Types as compared to the Type-specific protocols. However, an emergence of highly abundant sRNAs (like rRNA fragments) significantly reduce relative percentages of sequencing reads of other, less abundant sRNA classes, e.g., in comparison to sRNA Type 1 and Type 2 detected by protocol T[1+2] (supplementary Fig. S14B). Because potential sRNA biomarkers, especially for minimal residual disease or early disease detection, are presumably very rare and have a specific RNA Type, it would be logical to identify their Types (in addition to their sequences) first and then use a protocol focusing on this RNA Type to maximize the sensitivity of their detection.

### Detection of stsRNA Types 3 and 4

After establishing protocols T[1+2], T[2] and T[1+2+3+4], we developed protocols T[3+4], T[3] and T[4]. First, we evaluated protocol T[3+4] as schematically described in Table 1. In theory, a combination of data obtained by protocols T[1+2] and T[3+4] could provide much more sensitive detection of all Types than protocol T[1+2+3+4]. Like protocol T[1+2], the stsRNA sequencing length profiles for protocol T[3+4] significantly varied between the mock control (Fig. 7A, left panel and supplementary Fig. S2D) and stsRNAs spiked in a representative human D (D2) plasma RNA sample (Fig. 7A, right panel), while there were few differences in profiles of stsRNAs spiked in the human H and D plasma RNA samples (supplementary Fig. S21A). Notably, the presence of naturally occurring sRNAs in the plasma samples provided almost even coverage of stsRNAs in the 20–50 nt range. As anticipated, using protocol T[3+4], we successfully enriched specifically for stsRNA Type 3 and Type 4, making up ∼90% of our library, with similar average values for stsRNA Type 3 (H samples: 39.80±0.17% and D samples: 39.92±0.81%) and stsRNA Type 4 (H: 49.39±0.94% and D: 49.90±0.51%) for stsRNAs spiked in the human H and D plasma RNA samples, respectively (supplementary Tables S12A–B) with the ratio between stsRNA Type 3 to stsRNA Type 4 estimated as about 1 to 1.25.

**FIGURE 7.**
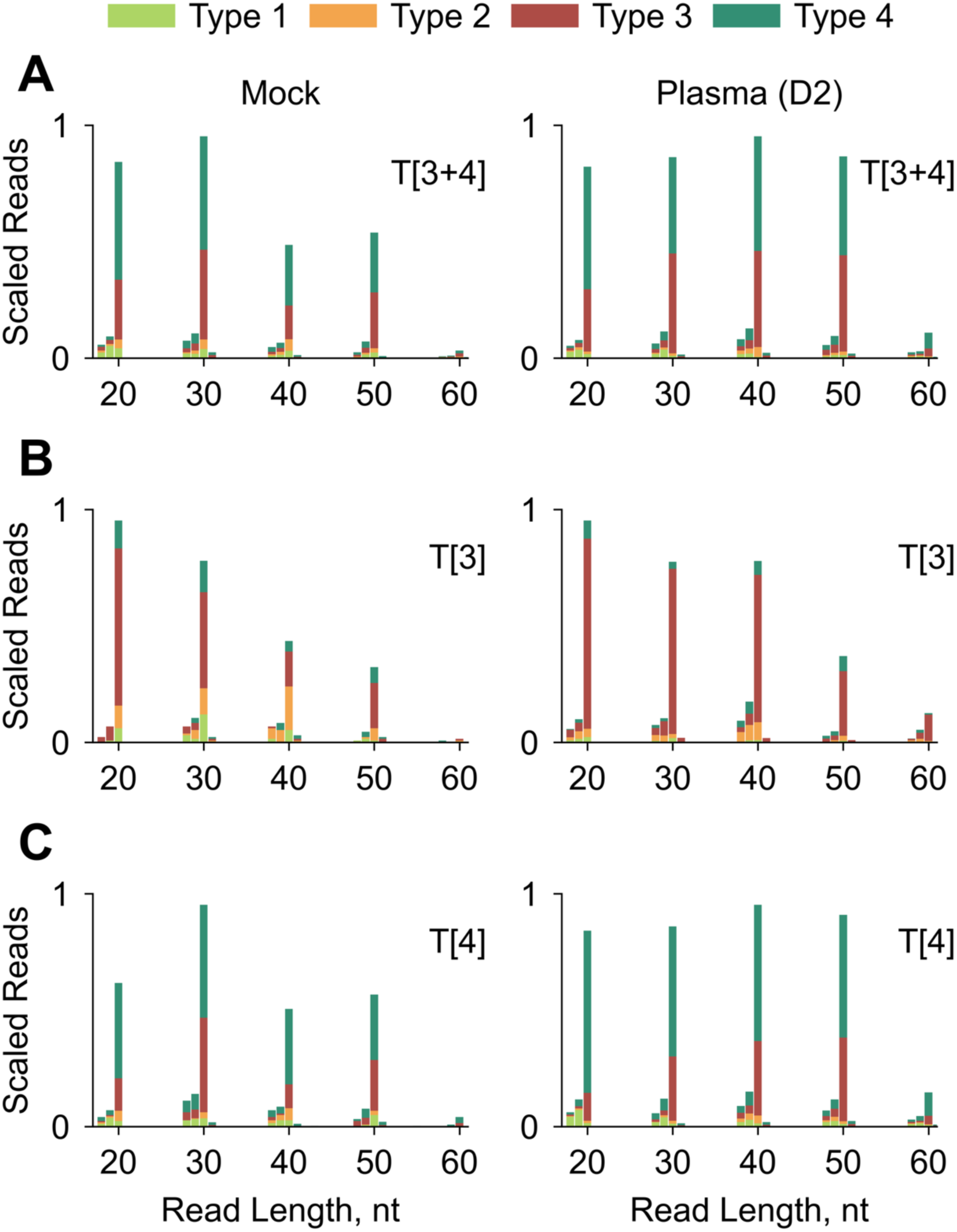
Representative sequencing length and RNA Type profiles for different stsRNA contexts, including: a mock control lacking a naturally occurring sRNA background (left column) and stsRNAs spiked into D2 human plasma RNA sample (right column), for libraries prepared by protocols (**A**) T[3+4], (**B**) T[3], or (**C**) T[4].

Second, we evaluated protocol T[3], schematically described in Table 1. In contrast to protocol T[2] enriching for another individual stsRNA Type (Fig. 2A–D, right panels), the stsRNA sequencing length profiles for protocol T[3] significantly varied between the mock control (Fig. 7B, left panel and supplementary Fig. S2E) and stsRNAs spiked in a representative human D (D2) plasma RNA sample (Fig. 7B, right panel) as well as between the human H and D plasma RNA samples (supplementary Fig. S21B). Both for mock control and stsRNAs spiked in the human plasma RNA samples, we found a steady decline of stsRNA capture in the 20–50 nt range. Also, we analyzed the RNA Type distributions for the stsRNA of different lengths spiked into the different human plasma RNA samples. In comparison to the mock control, the background of the naturally occurring RNA significantly improved the stsRNA Type 3 specificity of this protocol for the stsRNAs spiked in the human plasma RNA samples, where average percentages of the stsRNA Type 3 had similar values (76.96±1.68% and 75.09±1.42%) for the human H and D plasma RNA samples, respectively (supplementary Tables S13A–B).

Third, we evaluated protocol T[4], schematically described in Table 1. In the first pretreatment step of this protocol, we circularized sRNA Type 3 by RtcB ligase to prevent their incorporation into sequencing libraries (like we used T4 RNA ligases to circularize sRNA Type 1). For this purpose, we adopted previously described RtcB ligase reaction conditions (Chakravarty et al. 2012; Petkovic and Müller 2015) without additional optimization. Similarly, as for protocol T[3], length profiles for protocol T[4] significantly varied between the mock control (Fig. 7C, left panel and supplementary Fig. S2F) and stsRNAs spiked in a representative human plasma RNA sample (Fig. 7C, right panel), while there were few differences in profiles of stsRNAs spiked in the human H and D plasma RNA samples (supplementary Fig. S21C). Protocol T[4] provided almost even coverage of stsRNA Type 4 in the 20–50 nt range. However, we did not achieve very strong enrichment for stsRNA Type 4 over stsRNA Type 3 (Fig. 7C) presumably because RtcB is not as effective as T4 RNA ligases in circularization of their sRNA substrates under tested conditions. Protocol T[4] yielded a stsRNA Type 4 to Type 3 enrichment ratio of approximately 2.1 to 1 for stsRNAs in human H plasma RNA samples (Type 4: 58.98±1.01% and Type 3: 27.89±2.06%) and 2.3 to 1 for stsRNAs in human D plasma RNA samples (Type 4: 61.95±4.40% and Type 3: 26.88±4.60%) (supplementary Tables S14A–B). Nevertheless, a comparison of sequencing profiles for protocols T[3+4], T[3], and T[4] provided clear discrimination between Type 3 and Type 4 of both stsRNAs (Fig.7) and naturally occurring sRNAs of different classes (Fig. 8 and supplementary Fig. 23). This discrimination is reminiscent of “a strong G/weak A” and vice versa displays in Maxam-Gilbert A- and G-specific reactions, in which the redundant sequencing information serves as a check on the identifications of these nucleotides (Maxam and Gilbert 1977).

**FIGURE 8.**
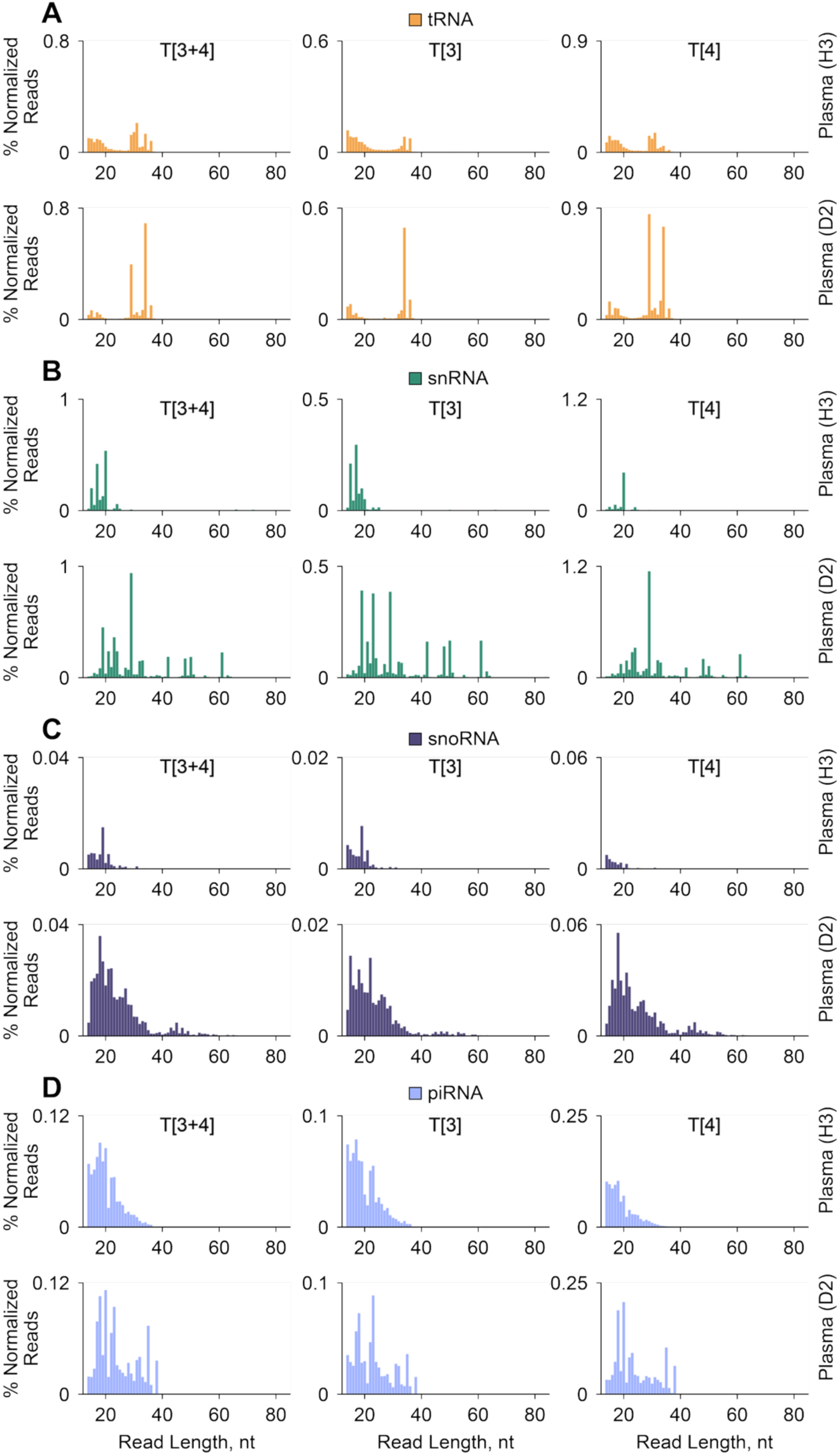
Comparison of sequencing length and RNA Type profiles for sRNAs derived from (**A**) tRNA, (**B**) snRNA, (**C**) snoRNA, and (**D**) piRNA from representative human plasma samples (indicated on the right) prepared by protocols T[3+4] (left column), T[3] (middle column), or T[4] (right column).

### Detection of naturally occurring RNA Types 3 and 4

After evaluation of protocols T[3+4], T[3], and T[4] using the stsRNAs, we explored naturally occurring sRNAs present in various human plasma RNA samples. Overall, sequencing length profiles showing all main sRNA classes simultaneously for protocol T[3+4] (supplementary Fig. S22A) were similar to the profiles of corresponding H and D human plasma RNA samples for protocol T[1+2+3+4] (supplementary Fig. S13) because of the global dominance of sequencing reads corresponding to RNA Type 3 and/or RNA Type 4 fragments of rRNA (supplementary Fig. S14C). A comparison of the percentage of normalized sequencing reads for the main individual sRNA classes in libraries of different human H and D plasma RNA samples detected by all RiboMarker^®^ protocols showed that sRNA Type 3 and/or Type 4 were also derived from tRNA, mRNA, lncRNA, and piRNA transcripts (supplementary Fig. S14C). The percentage of detected sRNA Types 3 and 4 was about the same for rRNA, snRNA, mRNA, and lncRNA, whereas sRNA Type 4 dominated over sRNA Type 3 for tRNA and snoRNA classes (supplementary Fig. S14C).

Although the differences in length profiles for all RNA classes simultaneously (supplementary Figs. S22A–C) were not easily notable for protocols T[3+4], T[3] and T[4], individual RNA classes had distinctive protocol-dependent profiles as well as sample differences between human H and D plasma RNA samples, especially for tRNAs, snRNAs, snoRNAs, and piRNAs (Fig. 8 and supplementary Figs. S23A–H). As shown in Fig. 8 for the representative human H and D plasma RNA samples, the human D plasma RNA sample for these four sRNA classes contained longer sequences, which were more dominant than shorter sRNA sequences in comparison to the human H plasma RNA sample. The distinctive features of sequencing length profiles for protocols T[3+4], T[3], and T[4], such as the length of sRNA sequences and/or ratio of sequencing reads between the sRNA sequences of the same lengths, could allow discrimination between sRNA Type 3 and Type 4. For example in the human D plasma RNA sample, a subset of 35-nt tRNA fragments could be identified as sRNA Type 3 whereas other 30-nt fragments were observed as sRNA Type 4 (Fig. 8A). Comparison of the sequencing profiles for the human H and D plasma RNA samples processed by all RiboMarker^®^ protocols, including protocols T[1+2] and T[1+2+3+4] (supplementary Figs. S16 and S17), T[2] (Fig. 4), T[3+4], T[3] and T[4] (Fig. 8 and supplementary Figs. S23A–H) allows a detailed analysis of the differences between the lengths of individual sRNAs and their RNA Types as well as highlights differences among different human plasma samples.

### Analysis of sRNA transcriptomics using all RiboMarker^®^ protocols

In contrast to sRNAs generated by random (or semi-random) fragmentation of RNAs, sRNAs derived from specific segments (or sites) of larger RNA transcripts may not only have different RNA Types but also be differentially abundant in biological samples. Comparative analysis of sRNA sequencing profiles for various RiboMarker^®^ protocols could serve for the identification of sRNA sequences of (biological and/or diagnostic) interest, including their RNA class(es) and RNA Type(s), and thereby a method to select a specific protocol that would provide detection of these sRNA(s) with highest sensitivity (comparatively to the other protocols). Moving beyond the sRNA sequencing length profiles, we have evaluated other analytical tools to compare the capacities of different protocols to detect sRNAs derived from main RNA classes for human H and D plasma samples.

First, to better understand the origin of sRNAs, we generated consensus mapping profiles of reads aligned for human H and D plasma RNA samples to both tRNA (Fig. 9A) and snRNA (Fig. 9B) transcripts. These data revealed that different Type-specific protocols captured sRNAs derived from distinct regions of their parent transcript. For tRNAs, protocol T[2] primarily captured tRNA fragments (tRFs) derived from the 3’-end, protocol T[4] from the 5’-end, and protocol T[3] from either end (Fig. 9A, left panels). In the case of snRNAs, sRNA Type 2 were found to be derived from their internal (middle) regions, sRNA Type 3 from both 3’- and 5’-ends, while sRNA Type 4 mostly overlapped with sRNA Type 2 molecules (Fig. 9B, right panels). We observed notable differences in these consensus profiles between human H and D plasma RNA samples, particularly for Type 4 tRNAs (Fig. 9A) and for Type 2 and Type 3 derived snRNA fragments (Fig. 9B). Although protocols detecting combinations of RNA classes (such as protocols T[1+2+3+4], T[1+2], and T[3+4]) also showed some differences between the sRNA consensus profiles, our data demonstrated that the Type-specific protocols are much better at isolating these differences.

**FIGURE 9.**
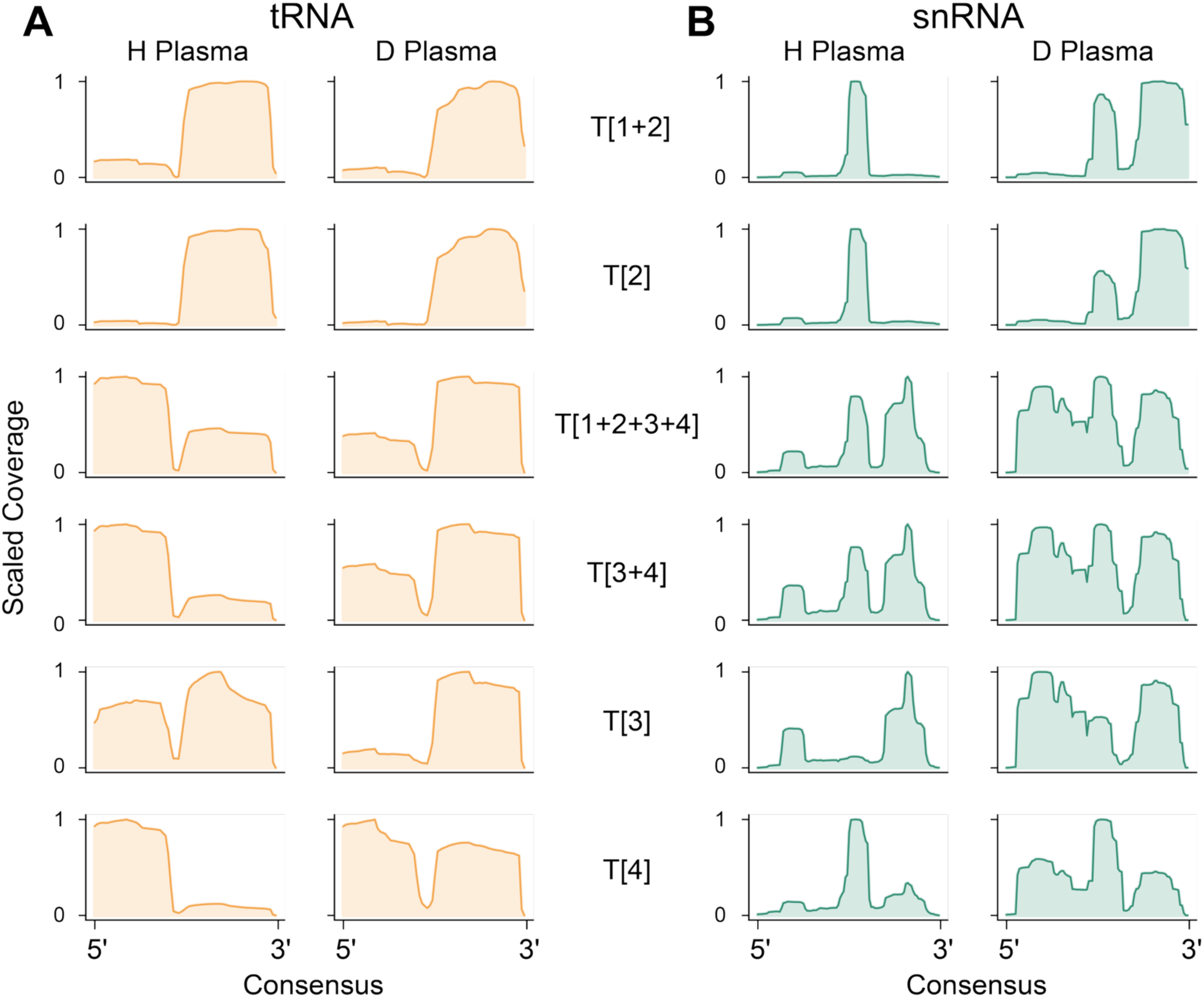
Consensus sequencing read coverage profiles for all (**A**) tRNA and (**B**) snRNA molecules, using the average distribution from either human H (H1, H2, H3) or D (D1, D2, D3) plasma RNA samples for libraries prepared using the indicated protocols (rows).

Second, we examined heatmaps visualizing patterns of RNA transcripts and/or fragmented sequences derived from different transcripts of all main RNA classes that were detected by the different RiboMarker^®^ protocols for individual H (Fig. 10A) and D (Fig. 10B) human plasma RNA samples. Rather than identifying specific protocols detecting entire sRNA classes with higher sensitivity than other protocols, these analyses facilitate the identification of the best protocol(s) for detecting specific groups of transcripts within each RNA class. In these figures, we marked groups of transcripts specifically enriched by different protocols using variable-colored lines. Also, we demonstrated that the different protocols could be matched with specific groups of transcripts within each RNA class because sRNAs derived from the same RNA class or even from the same RNA transcript could have different RNA Types. For example, comparative sizes of the marked heatmap segments with specifically enriched transcripts by different protocols were found to be protocol T[1+2] ∼ T[2] > T[4] >T[3+4] > T[3] for tRNA fragments and protocol T[1+2] >> T2 > T[4] for miRNAs in both human H and D plasma RNA samples (Figs. 10A–B). However, the number of tRNA fragments detected in human H plasma RNA was ∼ 2 times greater in comparison to human D plasma RNA samples. Although the dominance of protocol T[1+2] for detecting Type 1 miRNAs was obvious and the presence of small fractions of miRNAs, which were detected by both protocols T[1+2] and T[2], could be explained by incomplete circularization of miRNAs, the uncovering of significant fractions of Type 4 miRNAs by protocol T[4] was intriguing. We also clearly identified shorter (e.g., 17-nt in length) fragments of miRNA in sequencing length profiles for protocol T[4] (supplementary Fig. S23C). Previously, small numbers of 3’-end phosphorylated miRNAs have also been detected by Phospho-RNA-Seq in human plasma (Giraldez et al. 2019) and in cells (Lai et al. 2023).

**FIGURE 10.**
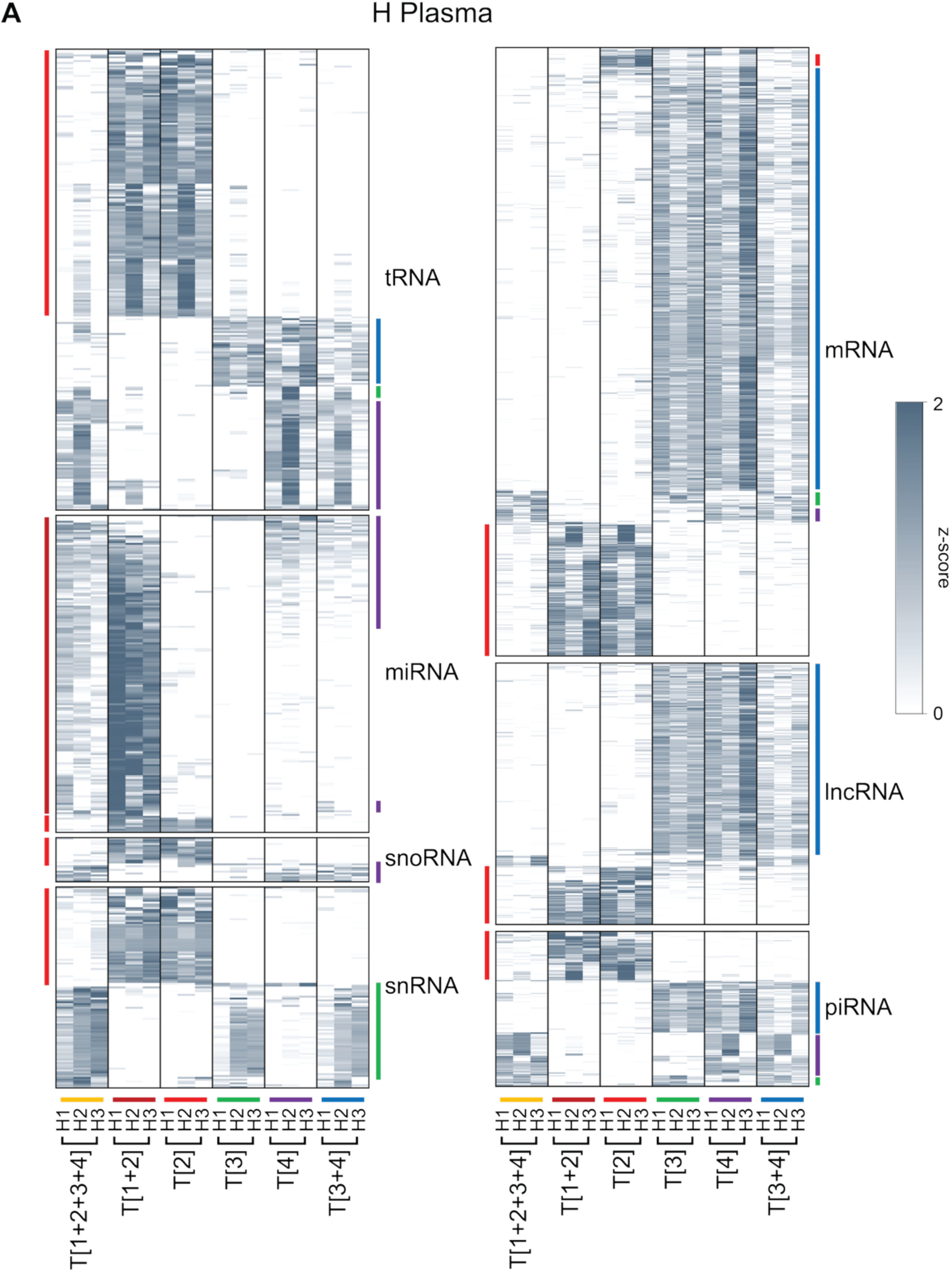
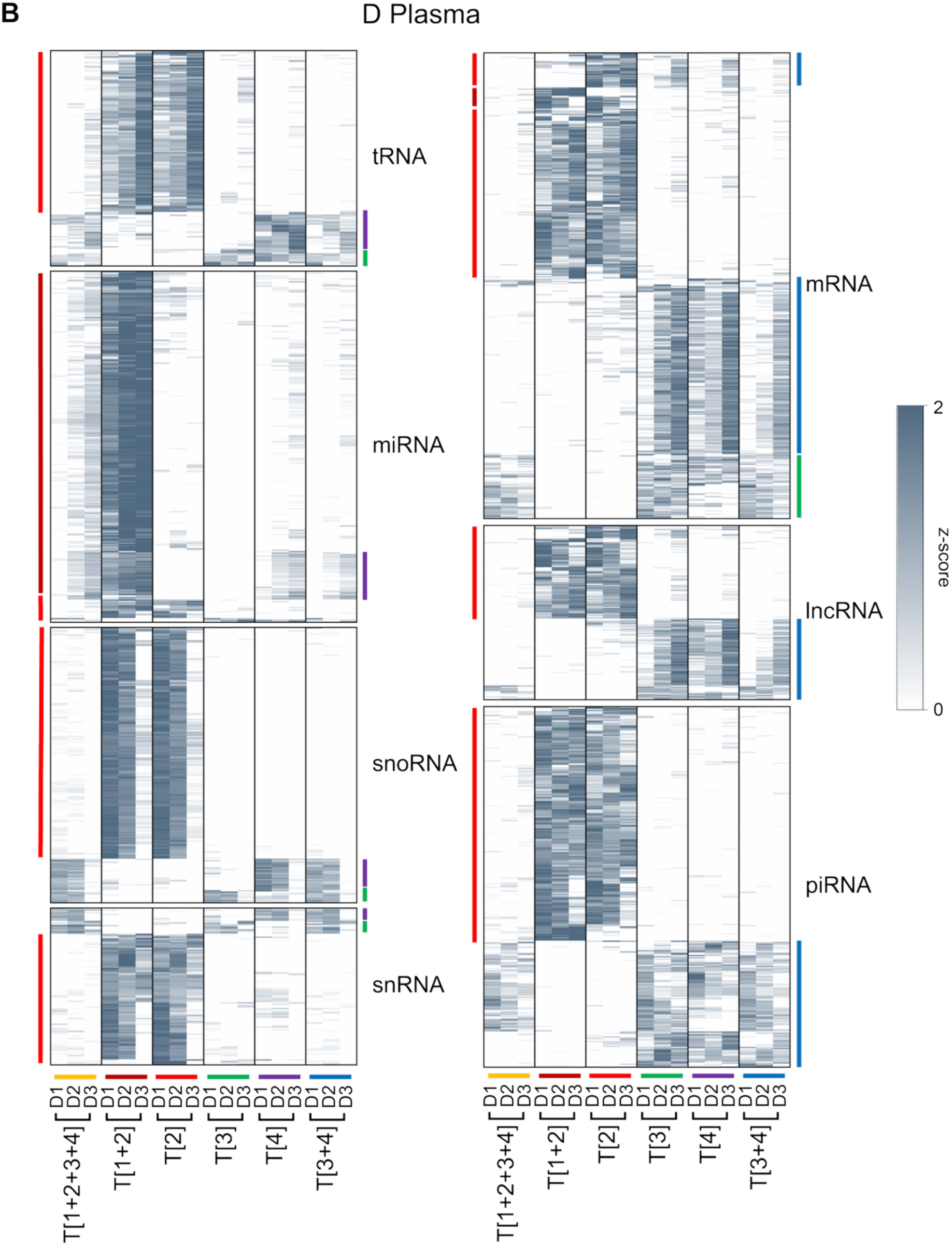
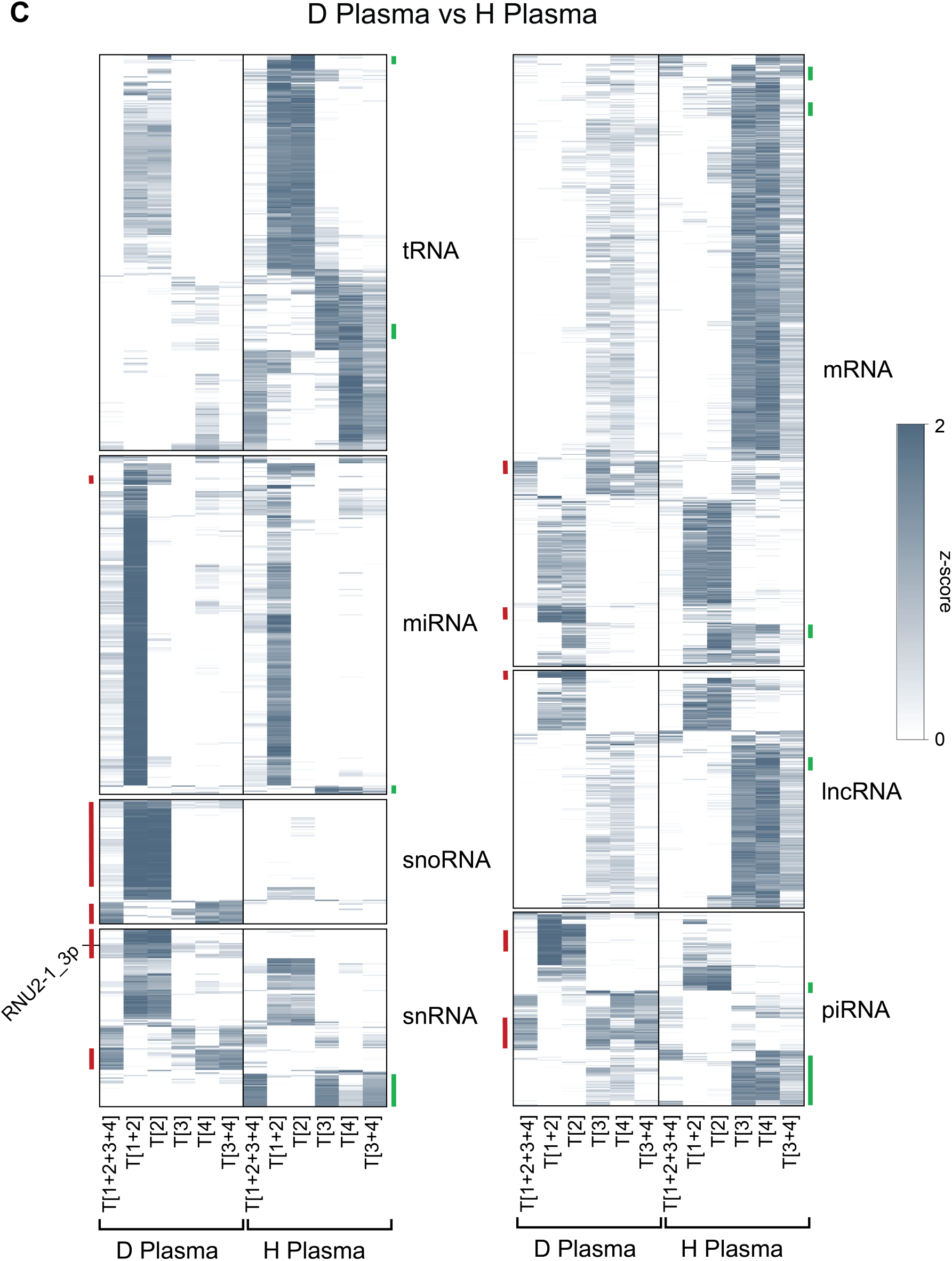
Hierarchically clustered heatmaps showing the relative abundance (row-wise z-score) of significantly variable RNA transcripts and/or fragments among protocols (DESeq2 LRT; padj < 0.05) from either human (**A**) H (H1, H2, H3) plasma or (**B**) D (D1, D2, D3) plasma RNA samples. Darker lines indicate a higher relative abundance of the corresponding RNA transcripts, and colored lines outside of the plot mark the protocol providing the highest sensitivity of their detection (relative enrichment). (**C**) Clustering of significantly enriched (DESeq2 LRT; padj < 0.05) sRNAs providing the best discrimination between human H or D plasma RNA samples using the various protocols. High-confidence D-enriched biomarkers are highlighted in red, H-enriched biomarkers in green.

In cases of other RNA classes, we also noticed significant differences in their Type enrichment. For instance, the ratio between the number of fragments derived from either snoRNAs or snRNAs was about 1 to 4 in human H plasma (Fig. 10A) and 2 to 1 in D plasma (Fig. 10B) samples. Comparative sizes of the heatmap segments with specifically enriched transcripts by different protocols for snoRNAs were found to be protocol T[1+2] ∼ T[2] > T[4] in the human H plasma and protocol T[1+2] ∼ T[2] >>T[4] > T[3] in the D plasma samples, whereas for snRNA, protocol T[1+2] ∼ T[2] ∼ T[3] in the human H plasma and protocol T[2] > T[1+2] >> T[4] ∼ T[3] in the D plasma samples (Figs. 10A–B). Other important RNA classes featured in this study were mRNA and lncRNA. Although the primary goal of the Phospho-RNA-Seq approach was to increase the overall recovery of unique mRNA and lncRNA fragments (Akat et al. 2019; Giraldez et al. 2019; Yao et al. 2020), we found that Phospho-RNA-Seq and protocol T[1+2+3+4] (supplementary Fig. 24) underperformed in this task in comparison to Type-specific protocols T[2], T[3] and T[4] (Figs. 10A–B). For instance, the best performing protocols for detection of sRNAs derived from mRNAs were protocol T[3+4] > T[1+2] ∼ T[2] > T[3] in the human H plasma and protocol T[3+4] ∼ T[2] > T[1+2] > T[3] > T[1+2] in the D plasma samples, whereas the best performing protocols for detection of lncRNA were protocol T[3+4] > T[2] > T[1+2] in the human H plasma and protocol T[2] > T[1+2] > T[3+4] in the D plasma samples (Figs. 10A–B). Regarding piRNAs, the ratio between the number of corresponding transcripts detected in either H plasma or D plasma samples was ∼1 to 2.5, and the best protocols for their detection were protocol T[1+2] ∼ T[2] ∼ T[3+4] > T[3] in the H plasma and protocol T[1+2] > T[2] > T[3+4] in the D plasma samples (Figs. 10A–B).

Additionally, we noticed differences among the detected transcript patterns for all featured RNA classes within the human H plasma (Fig. 10A) and D plasma (Fig. 10B) sample groups that indicated biological variations between the donors. Overall, we did not find any individual RNA class for which RiboMarker^®^ protocol T[1+2+3+4] outperformed the Type-specific protocols in variety and sensitivity of detecting transcripts for all RNA classes featured in this study (Figs. 10A–B). Interestingly, a comparison of libraries prepared using either protocol T[1+2+3+4] or Phospho-RNA-Seq on human plasma RNA samples (supplementary Fig. S24) revealed striking differences, indicating that these two protocols preferentially detected different groups of sRNA transcripts (with some overlaps) within every RNA class analyzed in this study, despite them capturing the same RNA Types.

### Discrimination between human H and D plasma samples as a prototype for sRNA biomarker discovery

Previously, we identified differences among sequencing profiles for the human H and D plasma RNA samples to highlight both technical aspects as well as biological variation observed using different RiboMarker^®^ protocols. In theory, these differential analyses laid the groundwork for the discovery of novel sRNA biomarkers to distinguish between the two plasma sample groups. The human H (H1, H2, H3) plasma samples were collected from three healthy donors, while the human D (D1, D2, D3) plasma samples were collected from three patients diagnosed with breast cancer (see more details in supplementary Table S15). Although our study was limited to a small sample size, these data serve as a prototype for utilizing the RiboMarker^®^ platform for biomarker discovery. Upon meeting the strict requirements for studies (including adequate numbers of plasma samples from large representative donor cohorts, controlled sample collection, handling, and processing), the workflow could be applied to reveal a suite of novel sRNA biomarkers for cancer and other various disease pathologies.

The significant differences among the sequencing profiles of H (healthy) and D (disease/cancer) human plasma RNA samples for certain sRNA classes have already been demonstrated by us here for selected RiboMarker^®^ protocols (see Figs. 4, 8, and 9). However, these profiles and the differences among them were determined for average data covering entire sRNA classes, and rather individual, selected sRNA sequences within these RNA classes could prove to be more sensitive and precise, as previously shown using miRNAs (Wang et al. 2018). To demonstrate the feasibility of this approach, we first identified the most significant differences among the human H and D plasma RNA samples for the main RNA classes (Fig. 10C). These data highlight the significantly enriched sRNAs providing the best discrimination between human H or D plasma RNA samples using the various RiboMarker^®^ protocols as marked with either red lines on the left for transcripts enriched in human D plasma samples or by green lines on the right for transcripts enriched in the H plasma samples. Although the enrichment of certain sRNAs in either human D or H samples could be used for biomarker selection, perhaps sRNAs that are unique for the D samples over the H samples could be better biomarker of the disease.

Among the analyzed RNA classes, sRNAs derived from snoRNAs and snRNAs provided the largest number of biomarker candidates (Fig. 10C). Without any specific preferences, we selected a group of fragments of RNU2-1 snRNA, hereby referred to as RNU2-1_3p, which we have marked in Fig. 10C, and whose sequences are shown in supplementary Fig. S25, as an example for further analysis. Protocols T[1+2] and T[2] were found to provide the best enrichment of RNU2-1_3p among the human D plasma samples to discriminate these versus the H plasma samples (Fig. 10C). Differential expression analyses (Fig. 11A) confirmed that RNU2-1_3p was indeed among the most significantly enriched sRNAs among the human D (D1, D2, D3) plasma versus the H (H1, H2, H3) plasma RNA samples using both protocols. Protocols T[1+2] and T[2] also excel in the sensitivity of detection of 24-nt, 52-nt, and 65-nt long RNU2-1 fragments that were found to be unique for both human D1 and D2 plasma samples (Fig. 11B). The profile of one of the human D (D3) plasma samples had only minor differences as compared to the human H plasma samples, which may be related to a different breast cancer subtype in the D3 sample in comparison to the human D1 and D2 plasma samples. Finally, we compared read coverage profiles (Fig. 11C) mapping to the RNU2-1 snRNA transcript found in these human H and D plasma samples for libraries prepared by protocols T[1+2] and T[2]. These profiles identified two areas, A and B, comprising sequences of the detected RNU2-1 snRNA fragments, where area A was specific to the human D plasma samples. In contrast, area B was common for both human H and D plasma samples (Fig. 11C). Also, the finding that protocol T[2] outperformed protocol T[1+2] in the detection of the RNU2-1 fragments in area A for the human D1 and D2 plasma samples (two bottom panels in Fig. 11C) indicated that that most of these RNU2-1 fragments had RNA Type 2 ends. The top 20 most abundant sRNA sequences aligned to the RNU2-1 snRNA that we detected in the selected human D (D1, D2) plasma RNA samples for libraries prepared by protocol T[2] are shown in supplementary Fig. S25.

**FIGURE 11.**
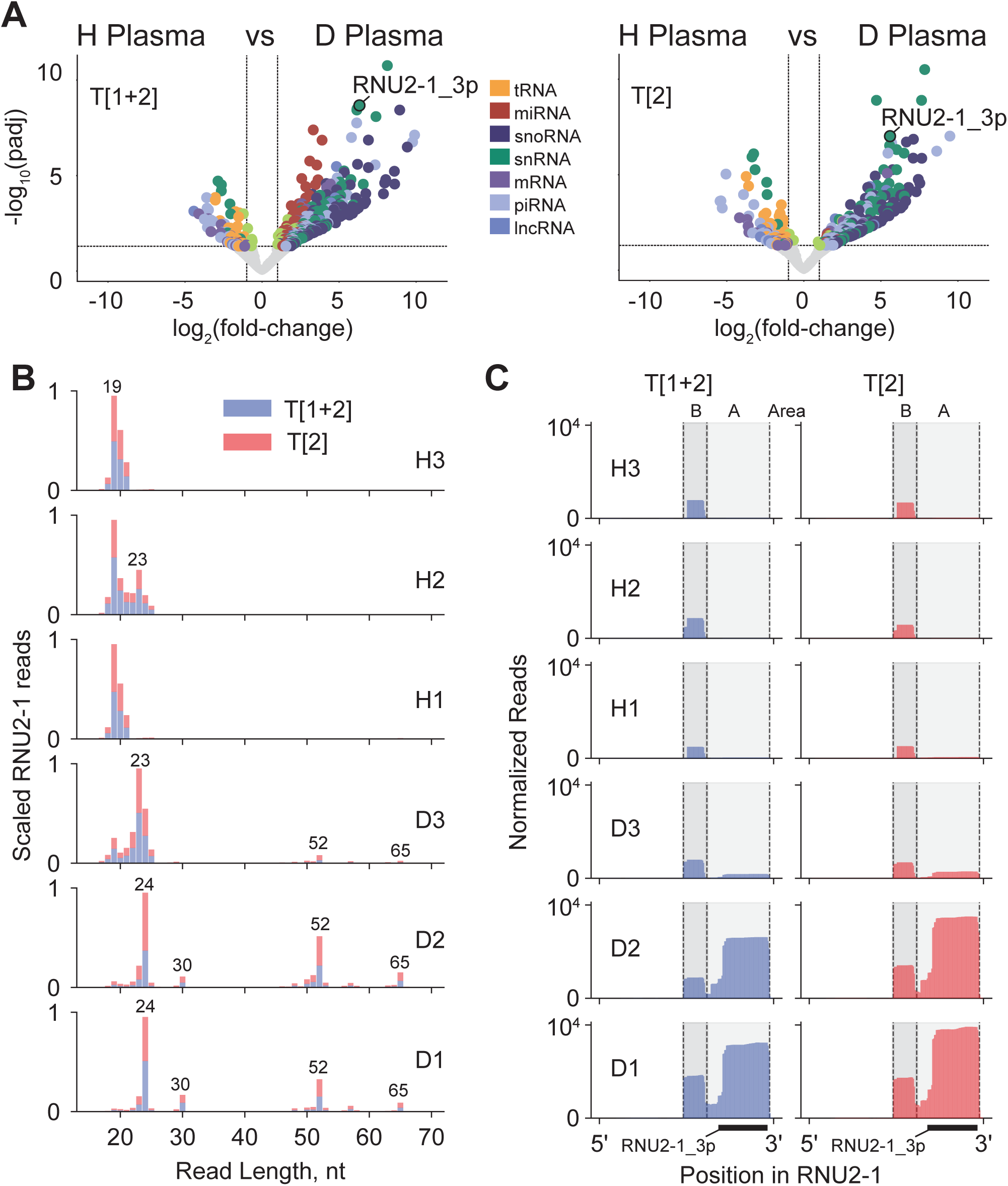
Differentially enriched sRNAs found among the human H and D plasma RNA samples. (**A**) Volcano plots showing the significant changes in (|log2fc|>1, padj<0.05) the abundance of sRNAs indicated by dots, whose colors correspond to the RNA classes they were derived from. Data for protocols T[1+2] and T[2] are shown in the left and right panels, respectively. (**B**) Overlapped sequencing length profiles for sRNAs derived from the RNU2-1 snRNA for libraries prepared using either protocols T[1+2] (blue) or T[2] (red) for human H (H1, H2, H3) and D (D1, D2, D3) plasma RNA samples. (**C**) Read coverage profiles for those reads mapping to the RNU2-1 snRNA using either protocols T[1+2] (blue; left) or T[2] (red; right) for human H (H1, H2, H3) and D (D1, D2, D3) plasma RNA samples. The region corresponding to the counts associated with the RNU2-1_3p feature in panel (**A**) is underlined, as well as the designated areas “A” and “B” of the RNU2-1 snRNA whose sequences are indicated in supplementary Fig. S25.

Although short 17–25-nt fragments derived from human RNU2-1 snRNA area B were previously identified as RNA Type 1 miR-U2-1 species in human blood using standard sRNA-Seq methods (Mazieres et al. 2013; Akat et al. 2019), the existence of sRNA Type 2 of longer 24–30-nt fragment sequences corresponding to area B and those 52–65-nt fragment sequences mapping to area A had not been observed. Additionally, the miR-U2-1 species were found in the blood of both cancer patients and healthy donors and have been implicated as potential multi-cancer diagnostic and prognostic biomarkers through their overexpression in non-small-cell lung, pancreatic, colorectal, and breast cancer (Mazieres et al. 2013; Hahn et al. 2015; Köhleret al. 2016; Pantazi et al. 2022). However, the RNA Type 2 RNU2-1 fragments could conceivably be alternative, more sensitive biomarkers than the miR-U2-1 fragments, as we observed higher disease specificity, and therefore hold promise for early detection of different cancer types and post-treatment minimal residual disease monitoring when they are more treatable. This example further emphasizes the general power of the RiboMarker^®^ platform for the identification and utilization of alternative Type-specific sRNAs to serve as novel, more potent biomarkers for cancer and other diseases.

The Ribomarker® platform enables a powerful capability to accurately quantify the complete RNA fragmentome. This work demonstrates that distinctive enzymatic pretreatments can be applied to RNA samples prior to universal sequencing library preparation. These pretreatments, which include enzymatic conversions of RNA termini and circularization, make it possible to either detect all or specifically enrich for individual sRNAs. The groundbreaking protocols can be used individually or in combination to enrich for sequencing reads of the RNA Type of greatest interest. The RiboMarker® platform protocols developed here offer a pivotal foundation to advance customizable RNA fragmentomics, empowering the search for novel biomarkers.

## MATERIALS AND METHODS

### Synthetic spike-in sRNA oligonucleotides

We designed 20 custom synthetic spike-in RNA oligonucleotides called stsRNAs. The core sequences of these stsRNAs, which do not have homology to the human genome, were adapted from Locati et al. (2015). The stsRNAs comprise 5’- and 3’-end combinations corresponding to four main RNA Types (1 through 4) shown in Fig. 1. For each RNA Type, there were five stsRNAs of different lengths: 20, 30, 40, 50, and 60 nt. The five stsRNAs for each RNA Type contain a 6-nt barcode that is RNA Type-specific in their middle section. Also, all stsRNAs feature 4 randomized nucleotides at both ends. The complete sequences of these stsRNAs are shown in Table 2. The stsRNAs were diluted to working concentrations with IDTE pH7.5 buffer (IDT) and equimolar pooled while working on ice to minimize RNA degradation. The resulting equimolar pool was aliquoted and stored immediately at -80°C. Sequences of combo adapter (CAD), blocking oligonucleotides, adapter dimer capture probes, RT blocking oligonucleotide, RT and PCR primers (schematically shown in supplementary Fig. S1) were the same as previously described in Barberán-Soler et al. (2018) except the CAD having an extra A ribonucleotide at its 3’-end. All synthetic oligonucleotides were synthesized and quantified spectrophotometrically by IDT.

### Biological samples

Reference human brain total RNA samples (Thermo Fisher Scientific) were diluted to 100 ng/µL with IDTE pH 7.5 buffer (IDT), aliquoted, and stored at -80°C. Commercially available human plasma samples collected in EDTA plasma tubes from various individuals are described in supplementary Table S15. Total RNA was extracted from plasma using Zymo Quick cfRNA Serum and Plasma kit (Zymo Research) according to the manufacturer’s protocol. Briefly, 1 mL of plasma was extracted per column. Columns were eluted with 12 µL of nuclease-free water so that 1 µL of extracted RNA is equivalent to the amount of RNA found in 83 µL of plasma. Plasma and extracted RNA was stored at -80°C.

### Experimental details of RiboMarker^®^ protocols

The RiboMarker^®^ protocols include RNA Type-specific enzymatic pretreatments of RNA samples and a universal library preparation protocol for the pretreated RNA samples (Table 1). The RNA sample inputs for each of these protocols were 50 ng of human brain total RNA or the total human plasma RNA amount equivalent to 83 µL of plasma. Each input of naturally occurring RNA was supplemented with 16 picomoles of synthetic RNA pool spike-ins (stsRNAs). Mock control RNA sample input comprises only 16 picomoles of the stsRNA pool without the naturally occurring RNA.

Protocol T[1+2] is used here as the universal method for sRNA sequencing library preparation method, and its experimental details are indicated in the next section. This is the only protocol that does not have a pretreatment step. All other RiboMarker^®^ protocols have pretreatment step(s) before the library preparation that are described below.

To select an optimal reaction condition for protocol T[2], we used the stsRNA pool spiked into human brain RNA samples incubated in standard T4 RNA ligase reaction buffer supplemented with 10% PEG 8000, 20 U Murine RNase Inhibitor (all NEB) and 100 µM ATP at 37°C with the following variables: (i) 10 U Rnl1 for 1 hour; (ii) 20 U Rnl1 for 1 hour; (iii) 20 U Rnl1 for 2 hours; or (iv) 10 U Rnl1 + 10 U Rnl2 for 1 hour. The selected protocol T[2] consisted of a single pretreatment reaction with a mix of T4 RNA ligase 1 and T4 RNA ligase 2 under condition (iv). The same reaction condition was also used for one of several pretreatment steps in protocols T[3+4], T[3] and T[4] as described in Table 1.

To select an optimal reaction condition for protocol T[1+2+3+4], we used the stsRNA pool spiked either into human brain RNA samples or plasma RNA samples (indicated in supplementary Table S9) incubated with 10 U T4 PNK (NEB), 20 U Murine RNase Inhibitor (NEB), 5 mM DTT (Thermo Fisher Scientific), 10 mM MgCl_2_ at 37°C for 30 minutes in various buffers, including: (a) 100 mM Tris-HCl at pH 7.6 in the presence of 1 mM ATP; (b) 100 mM Imidazole-HCl at pH 6.5 in the absence of ATP; or (c) 100 mM MES-NaOH at pH 6.0 in the absence of ATP. The selected protocol T[1+2+3+4] had a single pretreatment reaction under condition (c).

Protocol T[3+4] involved two sequential pretreatment reactions. The first reaction is 10 U T4 PNK 3’ minus, 10 U T4 RNA ligase 1, 10 U T4 RNA ligase 2, 20 U Murine RNase Inhibitor, 1 mM ATP, 10% PEG8000, and 1X T4 PNK reaction buffer pH7.6 (all NEB), incubated at 37°C for 1 hour. The second reaction is 10 U T4 PNK (NEB), 20 U Murine RNase Inhibitor (NEB), 5 mM DTT (ThermoFisher), 10 mM MgCl_2_, and 100 mM MES pH 6.0 (ThermoFisher), incubated at 37°C for 30 minutes.

Protocol T[3] involved three sequential pretreatment reactions. The first reaction is 1 U Terminator 5’-Phosphate-dependent exonuclease (LCG Biosearch Technologies), 1X Terminator Reaction Buffer A (LCG Biosearch Technologies), and 10 U RNase Inhibitor (NEB), incubated at 37°C for 1 hour. This reaction was terminated by adding 1 µL of 100 mM EDTA pH 8.0 with a final reaction concentration of 5 mM EDTA. The second reaction is 10 U T4 PNK 3’ minus, 10 U T4 RNA ligase 1, 10 U T4 RNA ligase 2, 20 U Murine RNase Inhibitor, 1 mM ATP, 10% PEG8000, and 1X T4 PNK reaction buffer pH7.6 (all NEB), incubated at 37°C for 1 hour. The third reaction is 10 U T4 PNK (NEB), 20 U Murine RNase Inhibitor (NEB), 5 mM DTT (ThermoFisher), 10 mM MgCl_2_, and 100 mM MES pH 6.0 (ThermoFisher), incubated at 37°C for 30 minutes.

Protocol T[4] consisted of three sequential pretreatment reactions. The first reaction is 30 pmol RtcB, 20 U Murine RNase Inhibitor, 100 µM MnCl_2_, 10% PEG8000, 1x RtcB Reaction Buffer (all NEB), incubated at 37°C for 1 hour as previously described (Chakravarty et al. 2012; Petkovic and Müller 2015). The second reaction is 10 U T4 PNK 3’ minus, 10 U T4 RNA ligase 1, 10 U T4 RNA ligase 2, 20 U Murine RNase Inhibitor, 1 mM ATP, 10% PEG8000, and 1X T4 PNK reaction buffer pH7.6 (all NEB), incubated at 37°C for 1 hour. The third reaction is 10 U T4 PNK (NEB), 20 U Murine RNase Inhibitor (NEB), 5 mM DTT (ThermoFisher), 10 mM MgCl_2_, and 100 mM MES pH 6.0 (ThermoFisher), incubated at 37°C for 30 minutes.

After each step of enzymatic pretreatments of the input RNA, the reactions were stopped by immediately processing the pretreated RNA through RNA Clean and Concentrator Kit columns (Zymo Research). The column washing protocol was modified to retain sRNAs of ≥ 15 nt in length. Briefly, 20 µL pretreatment reactions were mixed with 40 µL Binding Buffer. The resultant 60 µL mix was then mixed with 150 µL 95% ethanol. The final mix was applied to the column and washed according to the manufacturer’s instructions. All pretreated RNA was eluted into 6 µL nuclease-free water and used as input for the next pretreatment reaction or directly into the sequencing library preparation as described below.

### Library preparation and sequencing

All enzymatically pretreated RNA samples were prepared for sequencing using universal library preparation protocol T[1+2], which was also applied to input RNA samples that had not been enzymatically pretreated. Protocol T[1+2] was a modified version of commercially available RealSeq^®^-Biofluids kit protocol, which is based on the original RealSeq^®^-AC method for preparation of small RNA sequencing libraries (Barberán-Soler et al. 2018). The protocol T[1+2] workflow is schematically shown in Supplemental Figure S1 and includes the following modifications to the RealSeq^®^-Biofluids protocol: (i) adding 100 µM ATP to the adapter blocking reaction of step 2; (ii) decreasing the concentration of ATP from 1 mM to 100 µM during circularization step 3; and (iii) decreasing the concentration of additionally added dNTPs in PCR step 6 from 3 mM to 1.2 mM. The RNA inputs for protocol T[1+2] were either: (i) 16 picomoles of stsRNAs (for mock controls); (ii) 50 ng of human brain total RNA with 16 picomoles of stsRNAs; (iii) RNA volume equivalent to 83 µL of human plasma with 16 picomoles of the stsRNAs; or (iv) 6 µL of pretreated RNA samples with an initial pretreatment input as in (i), (ii), and (iii).

Phospho-RNA-Seq libraries were prepared with the sRNA input of the extracted total RNA volume equivalent to 83 µL of human plasma with 16 picomoles of the stsRNA pool. The RNA was pretreated in a reaction of 10 U T4 PNK, 20 U Murine RNase Inhibitor, 1 mM ATP, and 1X T4 PNK Reaction Buffer pH7.6 (all NEB), incubated at 37°C for 30 minutes. Following the pretreatment incubation, the reactions were stopped by immediately processing the pretreated RNA using RNA Clean and Concentrator Kit (Zymo Research). Similar to RiboMarker^®^ protocol T[1+2+3+4], the column-washing protocol was modified to retain sRNAs of ≥ 15 nt in length. The pretreated RNA was eluted into 6 µL nuclease-free water and used as input directly into the NEBNext Small RNA Library Prep Set for Illumina (NEB). The libraries were prepared according to the manufacturer’s protocol through PCR amplification. To more accurately compare Phospho-RNA-Seq and RiboMarker^®^ prepared libraries, 50 µL of the Phospho-RNA-Seq PCR reaction was cleaned up following the same SPRI bead size selection protocol used in RiboMarker^®^ protocol T[1+2].

During the preparation of sequencing libraries, PCR was performed with indexed primers to enable identification of reads unique to each sample during multiplex sequencing. The libraries were analyzed with an Agilent D1000 ScreenTape on a 4150 TapeStation instrument (Agilent) and then quantified with the Tape Station computer program and the Qubit dsDNA BR Assay kit (Thermo Fisher Scientific). Libraries were pooled and then sequenced in multiplex using the Singular Genomics G4 as 130 bp single end runs.

### Computational methods

#### stsRNA quantification

To robustly assess the efficiency of capture of differentially fragmented RNAs based on end chemistries and fragment length, 20 unique synthetic RNAs were generated and henceforth referred to as “stsRNAs”. For each of the four different RNA Types 1 through 4 (Fig. 1), a unique type-specific barcode was encoded in the internal part of the sequence (Table 2) to distinguish between the different types during mapping.

Mapping to these sequences was done after adapter trimming using bowtie2 (Martin 2011) [--no-1mm-upfront --mp 6 --rdg 6,3 --rfg 6,3 --norc --n-ceil L,5,0.15 -k 1 -D 20 -R 2 -N 0 -L 10 -i S,1,0.40 --score-min L,0,-0.6 --np 1] to account for the 4-nt randomized ends, internal type-specific barcodes, and differing stsRNA lengths. A custom Python script was used to parse the alignments for each sample and quantify individual stsRNAs based on these mappings.

#### Read processing and feature counting

Sequencing reads were trimmed of the adapter sequence using cutadapt (Martin 2011) and then further filtered to remove any reads less than 15 nt in length. Reads were then sequentially aligned using either bowtie or bowtie2 (Langmead and Salzberg 2012) to vector contaminants (UniVec), rRNAs, tRNAs (with the addition of the -CCA tail) identified using tRNAScan-SE (Chan et al. 2021), miRNAs from miRbase (Kozomara et al. 2019), snRNAs/snoRNAs from Ensembl (Dyer et al., 2025), piRNAs from piRbase (Wang et al. 2022), processed mRNA transcripts including 5’UTR, CDS and 3’UTR (mRNA/protein_coding) from Ensembl (Dyer et al. 2025), and lastly to the genome. Reads associated with these features were then quantified using a custom Python script, choosing the feature annotation with the largest degree of overlap. Multimapping ncRNA reads with more than one highest-scoring alignment were randomly selected, while multimapping mRNA reads were not included in the downstream analysis. For tRNA, snRNA, and snoRNA mapping reads, a read was assigned as either a ‘5p’, ‘3p’, or an ‘internal’ feature based on where it mapped in the parent transcript. For ‘5p’ features, it must map within 10 nt of the 5’ end, ‘3p’ features must map within 10 nt of the 3’ end, and an ‘internal’ feature does not map within 10 nt of either the 5’ or 3’ end of the parent transcript. The resulting read data underwent normalization and differential expression analysis (volcano plots and heatmaps showing significant features) using DESeq2 (Love 2014). Visualizations were generated using custom Python scripts. Raw fastq files are available on the NCBI Sequence Read Archive database under bioproject accession: PRJNA1348206.

#### stsRNA and natural RNA read length profiles

The proportion (P) of a given stsRNA Type, or RNA class (*T*) (for natural RNAs, excluding unmapped reads), at a specified read length (*L*) was calculated by dividing its raw read count (*C_T,L_*) by the total number of reads across all classes and lengths for its respective category (i.e., all stsRNA or all naturally occurring RNAs). This is shown by the following equation:

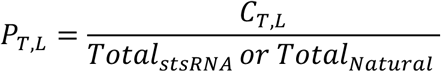

#### Consensus read mapping profiles

The distribution of read coverage across transcript bodies was calculated from the 5’-end transcript start to the 3’ transcript end by first generating a vector of read coverage at each position in a given RNA transcript. To normalize for different transcript lengths, each coverage vector was partitioned into 100 equal-sized bins, and the total counts within each bin were summed. These 100-point vectors were then aggregated by summing them for all transcripts within a given RNA class (e.g., tRNA, snRNA) for each sample. To create a final profile for each experimental protocol, these aggregated vectors were averaged across all biological replicates. For visualization, the resulting average profiles were min-max scaled to a range of [0,1] to emphasize the shape of the read distribution.

#### Differential read length profiles

The differential read length profiles were generated by calculating the proportion (P_T,L_) of a given stsRNA Type, or RNA class (*T*) (for natural RNAs, excluding unmapped reads), at a specified read length (*L*) for libraries prepared using different methods. These were then subtracted from one another to yield the percent difference as shown below.

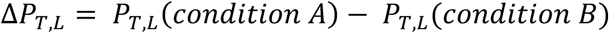

## ACKNOWLEDGMENTS

We thank Dr. Denise McGrath for helpful discussions, comments and technical editing of the manuscript. This work was supported in part by NIH grants 1R43HG013284-01 and 2R44HG013284-02A1 to S.A.K.

