## Supplementary Materials for "A method of comprehensive sequencing analysis of the small RNA fragmentome (RiboMarker^®^)"

**Supplementary TABLE S1.** Suggested RNA Type-specific alternative protocols for the preparation of sequencing libraries using standard RNA Type 1-specific methods of sRNA-Seq library preparation.

| Suggested Ribomarker® Protocols | ST[1] | ST[2] | ST[1+2+3+4] | ST[2+3] | ST[3+4] | ST[4] | ST[1+2] | ST[3] |
|---|---|---|---|---|---|---|---|---|
| RNA Types in the sample | 1, 2, 3, 4 | 1, 2, 3, 4 | 1, 2, 3, 4 | 1, 2, 3, 4 | 1, 2, 3, 4 | 1, 2, 3, 4 | 1, 2, 3, 4 | 1, 2, 3, 4 |
| Pretreatment # 1 | – | Rnl1 + Rnl2 ligases (+ATP) | PNK (-ATP) pH 6.0 | PNK (-ATP) pH 6.0 | PNK, 3'-minus (+ATP) pH 7.6 | RtcB ligase | PNK, 3'-minus (+ATP) pH 7.6 | Terminator 5'-p-dependent exonuclease |
| RF Types conversion | – | 1 → 0 | 3 → 2<br>4 → 1 | 3 → 2<br>4 → 1 | 2 → 1<br>3 → 4 | 3 → 0 | 2 → 1 | Degrade 1 , 4 |
| Pretreatment # 2 | – | PNK, 3'-minus (+ATP) pH 7.6 | PNK (+ATP) pH 7.6 | Rnl1 + Rnl2 ligases (+ATP) | Rnl1 + Rnl2 ligases (+ATP) | PNK, 3'-minus (+ATP) pH 7.6 | – | PNK, 3'-minus (+ATP) pH 7.6 |
| RNA Types conversion | – | 2 → 1 | 2 → 1 | 1 → 0 | 1 → 0 | 2 → 1 | – | 2 → 1<br>3 → 4 |
| Pretreatment # 3 | – | – | – | PNK (+ATP) pH 7.6 | PNK (-ATP) pH 6.0 | Rnl1 + Rnl2 ligases (+ATP) | – | Rnl1 + Rnl2 ligases (+ATP) |
| RF Types conversion | – | – | – | 2 → 1 | 4 → 1 | 1 → 0 | – | 1 → 0 |
| Pretreatment # 4 | – | – | – | – | – | PNK (-ATP) pH 6.0 | – | PNK (-ATP) pH 6.0 |
| RNA Types conversion | – | – | – | – | – | 4 → 1 | – | 4 → 1 |
| Sequencing library preparation Protocol | ST[1] | ST[1] | ST[1] | ST[1] | ST 1] | ST[1] | ST[1] | ST[1] |
| RNA Type(s) depleted from library | 2, 3 , 4 | 1 , 3 , 4 | None | 1 , 4 | 1 , 2 | 1 , 2 , 3 | 3 , 4 | 1 , 2, 4 |
| RNA Type(s) enriched in library | 1 | 2 | 1 , 2 , 3 , 4 (All Types) | 2 , 3 (5'-OH Types) | 3 , 4 (3'-P Types) | 4 | 1 , 2 (3'-OH Types) | 3 |

**Supplementary TABLE S2.** Average percentage of RNA Types for stsRNA sequences of all lengths spiked in human brain and plasma RNA samples, in triplicate, for libraries prepared by protocol T[1+2]. Mock controls (without input of naturally occurring RNA) were performed in duplicate.

| RNA sample | Mock Control | Brain | Plasma (H) | Plasma (D) |
|---|---|---|---|---|
| stsRNA Type | | | | |
| 1 | 47.01 ± 0.48 | 53.64 ± 0.36 | 42.40 ± 0.30 | 43.24 ±0 .59 |
| 2 | 52.32 ± 0.56 | 45.97 ± 0.33 | 57.72 ± 0.42 | 56.60 ± 0.38 |
| 3 | 0.28 ± 0.03 | 0.14 ± 0.02 | 0.06 ± 0.01 | 0.07 ± 0.01 |
| 4 | 0.44 ± 0.04 | 0.25 ± 0.05 | 0.06 ± 0.01 | 0.09 ± 0.01 |

**Supplementary TABLE S3.** Average percentage of RNA Types for stsRNA sequences of all lengths spiked in human brain and plasma RNA samples, in triplicate, for libraries prepared by protocol T[2] under the indicated ligation reaction conditions. Mock controls (without input of naturally occurring RNA) were performed in duplicate.

| Reaction conditions | Rnl1_1h | 2xRnl1_1h | 2xRnl1_2h | Rnl1+Rnl2_ 1h | Rnl1+Rnl2_ 1h | Rnl1+Rnl2_ 1h | Rnl1+Rnl2_ 1h |
|---|---|---|---|---|---|---|---|
| RNA sample | Brain | Brain | Brain | Brain | Plasma (H) | Plasma (D) | Mock Control |
| stsRNA Type | | | | | | | |
| 1 | 7.99 ± 0.11 | 8.89 ± 0.02 | 6.52 ± 0.21 | 8.31 ± 0.16 | 6.30 ± 0.51 | 5.71 ± 0.21 | 6.67 ± 0.23 |
| 2 | 90.59 ± 0.28 | 89.71 ± 0.02 | 83.80 ± 0.32 | 90.91 ± 0.19 | 93.50 ± 0.51 | 93.90 ± 0.24 | 91.53 ± 0.23 |
| 3 | 0.89 ± 0.09 | 1.07 ± 0.00 | 8.25 ± 0.03 | 0.56 ± 0.02 | 0.14 ± 0.01 | 0.29 ± 0.19 | 0.97 ± 0.03 |
| 4 | 0.33 ± 0.02 | 0.33 ± 0.00 | 1.43 ± 0.08 | 0.21 ± 0.00 | 0.07 ± 0.01 | 0.12 ± 0.04 | 0.82 ± 0.03 |

**Supplementary TABLE S4.** Average percentage of stsRNA Types of all lengths in sequencing libraries prepared in duplicate by various RiboMarker® protocols in the absence of naturally occurring RNAs (mock controls).

| Protocol | T[1+2] | T[2] | T[1+2+3+4] | T[3+4] | T[3] | T[4] |
|---|---|---|---|---|---|---|
| stsRNA Type | | | | | | |
| 1 | 47.01 ± 0.48 | 6.67 ± 0.23 | 27.63 ± 0.68 | 8.56 ± 0.32 | 10.31 ± 0.24 | 9.00 ± 0.50 |
| 2 | 52.32 ± 0.56 | 91.53 ± 0.23 | 27.14 ± 0.45 | 6.90 ± 0.26 | 20.01 ± 0.85 | 7.08 ± 0.13 |
| 3 | 0.28 ± 0.03 | 0.97 ± 0.03 | 17.76 ± 0.25 | 32.63 ± 1.41 | 53.07 ± 1.61 | 27.94 ± 0.73 |
| 4 | 0.4 ± 0.04 | 0.82 ± 0.03 | 27.47 ± 0.89 | 51.91 ± 2.00 | 16.61 ± 2.22 | 55.98 ± 1.10 |

**Supplementary TABLE S5.** Average percentage of RNA Types for stsRNA sequences of different lengths in the human H (H1, H2, H3) plasma RNA samples for libraries prepared by protocol T[1+2].

| stsRNA length | 20 | 30 | 40 | 50 | 60 | All lengths |
|---|---|---|---|---|---|---|
| stsRNA Type | | | | | | |
| 1 | 56.39 ± 0.48 | 41.74 ± 0.30 | 31.31 ± 0.98 | 40.0 2± 0.22 | 50.96 ± 1.13 | 42.16 ± 0.48 |
| 2 | 43.54 ± 0.48 | 58.12 ± 0.29 | 68.58 ± 0.98 | 59.80 ± 0.21 | 48.83 ± 1.18 | 57.72 ± 0.48 |
| 3 | 0.02 ± 0.01 | 0.08 ± 0.01 | 0.04 ± 0.01 | 0.10 ± 0.01 | 0.09 ± 0.03 | 0.06 ± 0.00 |
| 4 | 0.05 ± 0.01 | 0.06 ± 0.01 | 0.06 ± 0.01 | 0.08 ± 0.01 | 0.12 ± 0.02 | 0.07 ± 0.00 |

**Supplementary TABLE S6.** Average percentage of RNA Types for stsRNAs of different lengths spiked in the human H (H1, H2, H3) plasma RNA samples for libraries prepared by protocol T[2].

| stsRNA length | 20 | 30 | 40 | 50 | 60 | All lengths |
|---|---|---|---|---|---|---|
| stsRNA Type | | | | | | |
| 1 | 11.36 ± 1.38 | 6.97 ± 0.42 | 3.43 ± 0.23 | 4.49 ± 0.44 | 13.24 ± 0.76 | 6.30 ± 0.51 |
| 2 | 88.48 ± 1.38 | 92.78 ± 0.43 | 96.42 ± 0.22 | 95.21 ± 0.44 | 86.39 ± 1.00 | 93.50 ± 0.51 |
| 3 | 0.10 ± 0.01 | 019 ± 0.01 | 0.09 ± 0.01 | 0.18 ± 0.01 | 0.22 ± 0.06 | 0.14 ± 0.01 |
| 4 | 0.06 ± 0.02 | 0.06 ± 0.01 | 0.6 ± 0.01 | 0.12 ± 0.02 | 0.15 ± 0.01 | 0.07 ± 0.01 |

**Supplementary TABLE S7.** Reduction of the average percentage of Type 1 stsRNAs of different lengths spiked in the human plasma H (H1, H2, H3) RNA samples for libraries prepared by protocol T[2] in comparison to protocol T[1+2].

| stsRNA length | 20 | 30 | 40 | 50 | 60 |
|---|---|---|---|---|---|
| RNA Type 1 in | | | | | |
| Protocol T[1+2] (from Table S5) | 56.39 | 41.74 | 31.31 | 40.02 | 50.96 |
| Protocol T[2] (from Table S6) | 11.36 | 6.97 | 3.43 | 4.49 | 13.24 |
| Reduction of RNA Type 1 | (56.39 - 11.36) / 56.39 x 100% = 79.85 | (41.74 - 6.97) / 41.74 x 100% = 83.30 | (31.31 - 3.43) / 31.31 x 100% = 89.05 | (40.02 - 4.49) / 40.02 x 100% = 87.90 | (50.96 - 13.24) / 50.96 x 100% = 74.02 |

**Supplementary TABLE S8A.** Reduction of the percentage of miRNA reads in a human brain RNA sample for libraries prepared by protocol T[2] in comparison to protocol T[1+2].

| Replicate | 1 | 2 | 3 | Average |
|---|---|---|---|---|
| Protocol T[1+2] | 20.07 | 21.70 | 20.18 | 20.65 ± 0.70 |
| Protocol T[2] | 2.84 | 1.61 | 1.53 | 1.99 ± 0.56 |
| Reduced by fold | 7.07 | 13.48 | 13.18 | 11.24 ± 2.80 |
| Reduction by % | (20.07 - 2.84) / 20.07 x 100% = 85.85 | (21.70 – 1.61) / 21.70 x 100% = 92.59 | (20.18 – 1.53) / 20.18 x 100% = 92.41 | 90.28 ± 2.96 |

**Supplementary TABLE S8B.** Reduction of the percentage of piRNA reads in a human brain RNA sample for libraries prepared by protocol T[2] in comparison to protocol T[1+2].

| Replicate | 1 | 2 | 3 | Average |
|---|---|---|---|---|
| Protocol T[1+2] | 5.17 | 5.66 | 5.38 | 5.40 ± 0.17 |
| Protocol T[2] | 4.16 | 4.09 | 4.17 | 4.14 ± 0.03 |
| Reduced by fold | 1.24 | 1.38 | 1.29 | 1.30 ± 0.05 |
| Reduction by % | (5.17 – 4.16) / 5.17 x 100% = 19.54 | (5.66 – 4.09) / 5.66 x 100% = 27.73 | (5.38 – 4.17) / 5.38 x 100% = 22.50 | 23.26 ± 2.98 |

**Supplementary TABLE S8C.** Reduction of the percentage of miRNA reads in human H (H1, H2, H3) RNA plasma RNA samples for libraries prepared by protocol T[2] in comparison to protocol T[1+2].

| Sample | H1 | H2 | H3 | Average |
|---|---|---|---|---|
| Protocol T[1+2] | 8.53 | 7.34 | 4.54 | 6.80 ± 1.51 |
| Protocol T[2] | 0.41 | 0.19 | 0.20 | 0.27 ± 0.06 |
| Reduced by fold | 20.80 | 38.63 | 22.70 | 27.38 ± 7.50 |
| Reduction by % | (8.53 – 0.41) / 8.53 x 100% = 95.19 | (7.34 – 0.19) / 7.34 x 100% = 97.41 | (4.54 – 0.20) / 4.54 x 100% = 95.59 | 96.10 ± 0.88 |

**Supplementary TABLE S8D.** Reduction of the percentage of piRNA reads in the human H (H1, H2, H3) plasma RNA samples for libraries prepared by protocol T[2] in comparison to protocol T[1+2].

| Sample | H1 | H2 | H3 | Average |
|---|---|---|---|---|
| Protocol T[1+2] | 4.52 | 4.14 | 9.05 | 5.90 ± 2.10 |
| Protocol T[2] | 4.33 | 3.47 | 7.15 | 4.98 ± 1.44 |
| Reduced by fold | 1.04 | 1.19 | 1.27 | 1.17 ± 0.05 |
| Reduction by % | (4.52 – 4.33) / 4.52 x 100% = 4.20 | (4.14 – 3.47) / 4.14 x 100% = 16.18 | (9.05 – 7.15) / 9.05 x 100% = 21.00 | 13.79 ± 6.39 |

**Supplementary TABLE S9.** Average percentage of RNA Types for stsRNAs spiked into either a human brain RNA sample or indicated plasma RNA samples for libraries prepared in triplicates by different versions of protocol T[1+2+3+4] using the indicated PNK pretreatment reaction conditions.

| Buffer | Tris-HCl pH 7.6 +ATP | Imidazole-HCl, pH 6.5 | MES-NaOH pH 6.0 | MES-NaOH pH 6.0 | MES-NaOH pH 6.0 | MES-NaOH pH 6.0 | MES-NaOH pH 6.0 |
|---|---|---|---|---|---|---|---|
| PNK Units | 10 | 10 | 10 | 10 | 10 | 10 | 20 |
| RNA sample | Brain | Brain | Brain | Plasma (H1, H2, H3) | Plasma (D1, D2, D3) | Plasma (NA2) | Plasma (NA2) |
| stsRNA Types | | | | | | | |
| 1 | 29.88 ± 0.28 | 30.68 ± 0.23 | 30.69 ± 0.50 | 28.37 ± 0.24 | 28.40 ± 0.49 | 27.10 ± 0.28 | 27.21 ± 0.04 |
| 2 | 32.27 ± 0.17 | 32.10 ± 0.23 | 31.93 ± 0.73 | 31.81 ± 0.34 | 31.54 ± 0.67 | 31.61 ± 0.17 | 31.37 ± 0.31 |
| 3 | 15.98 ± 0.17 | 14.72 ± 0.23 | 14.76 ± 0.35 | 18.72 ± 0.19 | 18.74 ±0.58 | 19.29 ± 0.03 | 18.61 ± 0.14 |
| 4 | 21.88 ±0.09 | 22.50 ± 0.06 | 22.62 ±0.09 | 21.09 ± 0.29 | 21.31 ± 0.39 | 22.00 ± 0.18 | 22.81 ± 0.21 |

**Supplementary TABLE S10.** Average percentage of RNA Types for stsRNAs of different lengths spiked in the human CL (CL1, CL2, CL4) plasma RNA samples for libraries prepared by RiboMarker® protocol T[1+2+3+4].

| stsRNA length | 20 | 30 | 40 | 50 | 60 | All lengths |
|---|---|---|---|---|---|---|
| stsRNA Type | | | | | | |
| 1 | 36.56 ± 3.66 | 25.57 ± 1.94 | 23.76 ± 1.18 | 19.79 ± 0.82 | 31.54 ± 3.07 | 26.56 ± 1.86 |
| 2 | 27.21 ± 1.78 | 30.01 ± 2.07 | 55.90 ± 4.90 | 29.21 ± 0.60 | 34.89 ± 1.45 | 35.15 ± 1.51 |
| 3 | 9.55 ± 2.03 | 20.24 ± 2.15 | 4.84 ± 2.09 | 27.06 ± 1.43 | 16.37 ± 2.17 | 15.80 ± 1.75 |
| 4 | 26.69 ± 3.23 | 24.18 ± 1.86 | 15.50 ± 2.85 | 23.94 ± 0.53 | 17.20 ± 0.79 | 22.49 ± 1.52 |

**Supplementary TABLE S11.** Average percentage of RNA Types for stsRNAs of different lengths spiked in the human CL (CL1, CL2, CL4) plasma RNA samples for libraries prepared by Phospho-RNA-Seq.

| stsRNA length | 20 | 30 | 40 | 50 | 60 | All lengths |
|---|---|---|---|---|---|---|
| stsRNA Type | | | | | | |
| 1 | 58.18 ± 1.98 | 19.69 ± 0.87 | 32.59 ± 0.70 | 17.32 ± 0.62 | 45.43 ± 1.02 | 29.84 ± 0.44 |
| 2 | 11.27 ± 0.34 | 28.69 ± 1.49 | 17.35 ± 0.53 | 37.48 ± 1.76 | 28.35 ± 0.61 | 23.35 ± 0.56 |
| 3 | 10.34 ± 1.23 | 24.26 ± 0.39 | 17.29 ± 0.25 | 22.80 ± 1.20 | 14.76 ± 0.67 | 19.02 ± 0.32 |
| 4 | 20.21 ± 0.95 | 27.36 ± 0.86 | 32.77 ± 0.72 | 22.39 ± 0.34 | 11.46 ± 0.35 | 27.79 ± 0.49 |

**Supplementary TABLE S12A.** Average percentage of RNA Types for stsRNAs of different lengths spiked in the human H (H1, H2, H3) plasma RNA samples in libraries prepared by protocol T[3+4].

| stsRNA length | 20 | 30 | 40 | 50 | 60 | All lengths |
|---|---|---|---|---|---|---|
| stsRNA Type | | | | | | |
| 1 | 9.12 ± 0.05 | 5.40 ± 0.29 | 3.88 ± 0.26 | 4.42 ± 0.24 | 28.48 ± 11.03 | 7.00 ± 0.79 |
| 2 | 2.37 ± 0.09 | 2.63 ± 0.22 | 38.53 ± 0.68 | 2.67 ± 0.35 | 2.67 ± 0.35 | 3.81 ± 0.16 |
| 3 | 32.72 ± 1.00 | 45.68 ± 0.32 | 38.53 ± 0.68 | 45.28 ± 0.90 | 25.58 ± 1.90 | 39.80 ± 0.17 |
| 4 | 55.79 ± 1.06 | 46.29 ± 0.29 | 50.61 ± 0.96 | 47.63 ± 1.39 | 40.87 ± 7.82 | 49.39 ± 0.94 |

**Supplementary TABLE S12B.** Average percentage of RNA Types for stsRNAs of different lengths spiked in the human D (D1, D2, D3) plasma RNA samples for libraries prepared by the protocol T[3+4].

| stsRNA length | 20 | 30 | 40 | 50 | 60 | All lengths |
|---|---|---|---|---|---|---|
| stsRNA Type | | | | | | |
| 1 | 9.55 ± 1.07 | 6.24 ± 1.03 | 4.23 ± 0.72 | 3.62 ± 0.48 | 12.52 ± 2.07 | 6.15 ± 0.70 |
| 2 | 2.37 ± 0.09 | 2.63 ± 0.22 | 6.98 ± 0.14 | 2.67 ± 0.35 | 2.67 ± 0.35 | 3.81 ± 0.16 |
| 3 | 31.17 ± 0.90 | 44.55 ± 1.69 | 39.77 ± 1.06 | 45.45 ± 1.46 | 30.11 ± 2.07 | 39.92 ± 0.81 |
| 4 | 56.26 ± 1.17 | 46.52 ± 0.63 | 48.75 ± 2.80 | 48.33 ± 2.27 | 52.45 ± 2.09 | 49.90 ± 0.51 |

**Supplementary TABLE S13A.** Average percentage of RNA Types for stsRNAs of different lengths spiked in the human H (H1, H2, H3) plasma RNA samples for libraries prepared by protocol T[3].

| stsRNA length | 20 | 30 | 40 | 50 | 60 | All lengths |
|---|---|---|---|---|---|---|
| stsRNA Type | | | | | | |
| 1 | 3.49 ± 0.43 | 2.94 ± 0.47 | 1.78 ± 0.65 | 1.79 ± 0.49 | 3.80 ± 1.75 | 2.67 ± 0.24 |
| 2 | 5.81 ± 1.11 | 8.46 ± 1.56 | 19.01 ± 0.82 | 5.89 ± 0.89 | 8.31 ± 1.59 | 10.11 ± 1.06 |
| 3 | 81.00 ± 1.07 | 83.37 ± 2.32 | 67.92 ± 1.61 | 75.09 ± 2.58 | 74.21 ± 3.04 | 76.96 ± 1.68 |
| 4 | 9.70 ± 0.66 | 5.23 ± 0.35 | 11.28 ± 0.50 | 17.24 ± 1.53 | 13.68 ± 1.32 | 10.26 ± 0.61 |

**Supplementary TABLE S13B.** Average percentage of RNA Types for stsRNAs of different lengths spiked in the human D (D1, D2, D3) plasma RNA samples for libraries prepared by protocol T[3].

| stsRNA length | 20 | 30 | 40 | 50 | 60 | All lengths |
|---|---|---|---|---|---|---|
| stsRNA Type | | | | | | |
| 1 | 4.43 ± 0.25 | 3.00 ± 0.51 | 2.79 ± 1.36 | 1.52 ± 0.42 | 5.11 ± 3.3 | 3.22 ± 0.73 |
| 2 | 7.10 ± 1.45 | 6.75 ± 1.01 | 15.46 ± 1.56 | 6.00 ± 1.37 | 9.87 ± 2.35 | 9.24 ± 0.63 |
| 3 | 74.86 ± 4.38 | 84.49 ± 0.54 | 67.54 ± 0.68 | 74.57 ± 1.75 | 69.35 ± 2.8 | 75.09 ± 1.42 |
| 4 | 13.61 ± 2.92 | 5.77 ± 0.08 | 14.20 ± 0.96 | 17.90 ± 0.04 | 15.67 ± 2.51 | 12.46 ± 1.26 |

**Supplementary TABLE S14A.** Average percentage of RNA Types for stsRNAs of different lengths spiked in the human H (H1, H2, H3) plasma RNA samples for libraries prepared by protocol T[4].

| stsRNA length | 20 | 30 | 40 | 50 | 60 | All lengths |
|---|---|---|---|---|---|---|
| stsRNA Type | | | | | | |
| 1 | 12.34 ± 0.46 | 7.05 ± 0.22 | 4.74 ± 0.15 | 4.62 ± 0.12 | 42.08 ± 8.61 | 10.13 ± 1.12 |
| 2 | 1.97 ± 0.22 | 2.26 ± 0.08 | 5.21 ± 0.33 | 2.25 ± 0.33 | 3.31 ± 0.91 | 3.00 ± 0.08 |
| 3 | 12.48 ± 1.42 | 30.70 ± 2.76 | 31.68 ± 2.19 | 39.01 ± 0.73 | 18.65 ± 3.23 | 27.89 ± 2.06 |
| 4 | 73.21 ± 1.36 | 59.99 ± 2.54 | 58.36 ± 2.40 | 54.12 ± 0.90 | 35.96 ± 4.48 | 58.98 ± 1.01 |

**Supplementary TABLE S14B.** Average percentage of RNA Types for stsRNAs of different lengths in the human D (D1, D2, D3) plasma RNA samples for libraries prepared by protocol T[4].

| stsRNA length | 20 | 30 | 40 | 50 | 60 | All lengths |
|---|---|---|---|---|---|---|
| stsRNA Type | | | | | | |
| 1 | 12.52 ± 0.40 | 6.71 ± 0.25 | 5.39 ± 0.58 | 4.89 ± 0.66 | 12.63 ± 1.91 | 7.49 ± 0.63 |
| 2 | 2.28 ± 0.73 | 2.79 ± 0.44 | 6.02 ± 0.69 | 3.10 ± 0.16 | 4.48 ± 0.34 | 3.69 ± 0.42 |
| 3 | 12.57 ± 2.02 | 29.54 ± 4.51 | 29.27 ± 5.22 | 35.66 ± 5.78 | 22.20 ± 3.01 | 26.88 ± 4.60 |
| 4 | 72.64 ± 2.32 | 60.96 ± 4.62 | 59.32 ± 5.32 | 56.34 ± 5.19 | 60.69 ± 1.69 | 61.95 ± 4.40 |

**Supplementary TABLE S15.** Specifications of human plasma samples used in this study.

| Internal ID | Vendor's Sample ID | Plasma Matrix Type | Donor's ethnicity and gender, age | Pathology Metadata |
|---|---|---|---|---|
| H1 | Innovative Research HMN756159 | Single healthy donor panel | Hispanic male, 41 | None |
| H2 | Innovative Research HMN756160 | Single healthy donor panel | Hispanic male, 42 | None |
| H3 | Innovative Research HMN756161 | Single healthy donor panel | Hispanic male, 29 | None |
| D1 | Innovative Research 200944-PL-2 | Single donor with breast cancer | Caucasian female, 58 | Invasive carcinoma, Stage: IIA |
| D2 | Innovative Research 200969-PL-5 | Single donor with breast cancer | Caucasian female, 66 | Invasive carcinoma, Stage: II |
| D3 | Innovative Research 200979-PL-5 | Single donor with breast cancer | Caucasian female, 53 | Invasive carcinoma, Stage: IIA |
| NA2 | Innovative Research IPLASNAE10ML-2396153A | Single healthy donor | Hispanic female, 40 | None |
| CL1 | Cureline CU170517-116-248BTB-3EVIIh BrCa P | Single donor with breast cancer | Caucasian female, 27 | Infiltrating ductal carcinoma, Stage: IIIC |
| CL2 | Cureline CU170517-116-652BTB-7CXh BrCa P | Single donor with breast cancer | Caucasian female, 35 | Infiltrating ductal carcinoma, Stage: III |
| CL4 | Cureline CU170517-116-312BTB-3LIh BrCa P | Single donor with breast cancer | Caucasian female, 38 | Infiltrating ductal carcinoma, Stage: IIIA |

#### SUPPLEMENTARY FIGURE LEGENDS

**Supplementary FIGURE S1.** Schematics of RiboMarker® protocol T[1+2] for sRNA sequencing library preparation. (**A**) General workflow including the ligation of a single combo adapter (CAD) to sRNA, circularization of the ligation products (RCAD), reverse transcription of the circularized RCAD, generating cDNA products, and PCR amplification of these cDNA. (**B**) Blocking of the unligated CAD. See text for details.

**Supplementary FIGURE S2.** Sequencing length and RNA Type profiles for stsRNAs in the absence of naturally occurring sRNAs (mock controls) for the libraries prepared by the indicated RiboMarker® protocols.

**Supplementary FIGURE S3.** Sequencing length and RNA Type profiles for stsRNAs spiked into human brain RNA samples for libraries prepared by protocol T[1+2] (left panels) or protocol T[2] (right panels) made in technical triplicates.

**Supplementary FIGURE S4.** Sequencing length and RNA Type profiles for stsRNAs spiked into either human H (H1, H2, H3; left panels) plasma or D (D1, D2, D3; right panels) plasma RNA samples for libraries prepared by protocol T[1+2].

**Supplementary FIGURE S5.** Sequencing length and RNA Type profiles for stsRNAs spiked into total RNA isolated from either human H (H1, H2, H3; left panels) plasma or D (D1, D2, D3; right panels) plasma RNA samples for libraries prepared by protocol T[2].

**Supplementary FIGURE S6.** Sequencing length and RNA Type profiles for stsRNAs spiked into human brain RNA samples for libraries prepared by protocol T[1+2] and different versions of protocol T[2] performed under the indicated RNA ligase reaction conditions. The profiles for stsRNA Type 3 and Type 4 are indicative of the yields of Rnl1 “reverse reaction” products.

**Supplementary FIGURE S7.** Sequencing length profiles of naturally occurring sRNA Type 1 found in a human brain total RNA sample that were determined by subtracting percentages of normalized sequencing reads for libraries prepared by protocol T[2] from those prepared by protocol T[1+2].

**Supplementary FIGURE S8.** Sequencing length profiles for indicated RNA classes of naturally occurring sRNAs found in either the indicated human H plasma (**A**) or D plasma (**B**) total RNA samples for libraries prepared by protocol T[1+2].

**Supplementary FIGURE S9.** Sequencing length and RNA Type profiles for stsRNAs spiked into human NA2 plasma RNA sample for libraries prepared by protocol T[1+2] either (**A**) directly (controls) or (**B**) after purification using the Zymo RNA clean and concentrator kit (column purified). The sequencing libraries were made and sequenced in technical triplicates.

**Supplementary FIGURE S10.** Sequencing length and RNA Type profiles for stsRNAs spiked into human NA2 plasma RNA sample for libraries prepared by protocol T[1+2+3+4] and with either (**A**) heat-inactivation of PNK (heated) or (**B**) removal of PNK using the Zymo RNA clean and concentrator kit (column purified). The sequencing libraries were made and sequenced in technical triplicates.

**Supplementary FIGURE S11.** Sequencing length and RNA Type profiles for stsRNAs spiked into human CL (CL1, CL2, CL4) plasma RNA samples for libraries prepared by either (**A**) RiboMarker® protocol T[1+2+3+4] or (**B**) Phospho-RNA-Seq .

**Supplementary FIGURE S12.** Sequencing length profiles for indicated RNA classes of naturally occurring sRNAs found in a human brain total RNA sample prepared with protocol T[1+2+3+4]. The sequencing libraries were made and sequenced in technical triplicates (Rep. 1, 2, 3).

**Supplementary FIGURE S13.** Sequencing length profiles for indicated RNA classes of naturally occurring sRNAs found in either human H plasma (H1, H2, H3; left panels) or D plasma (D1, D2, D3; right panels) total RNA samples for libraries prepared by protocol T[1+2+3+4].

**Supplementary FIGURE S14.** Percentage of normalized sequencing reads of main RNA classes for libraries prepared for: (**A**) human brain total RNA samples (1, 2, 3); (**B**) human H (H1, H2, H3) plasma or D (D1, D2, D3) plasma RNA samples using either protocol T[1+2] or protocol T[1+2+3+4] with numbers indicating the average fold change for inclusion of the indicated sRNA classes into sequencing libraries between these two protocols; and (**C**) the same (as above) human H and D plasma total RNA samples prepared by protocols T[2], T[3], T[4] or T[3+4].

**Supplementary FIGURE S15.** Comparison of sequencing length profiles for naturally occurring sRNAs found in a human brain total RNA sample among libraries prepared by protocols T[1+2] and T[1+2+3+4] for the following RNA classes: (**A**) rRNA, (**B**) miRNA, (**C**) tRNA, (**D**) snRNA, (**E**) snoRNA, (**F**) mRNA, (**G**) lncRNA and (**H**) piRNA.

**Supplementary FIGURE S16.** Comparison of sequencing length profiles for naturally occurring sRNAs found in the representative human H (H3) plasma RNA sample between libraries prepared by protocols T[1+2] and T[1+2+3+4] for the indicated RNA classes: (**A**) rRNA, (**B**) miRNA, (**C**) tRNA, (**D**) snRNA, (**E**) snoRNA, (**F**) mRNA, (**G**) lncRNA and (**H**) piRNA.

**Supplementary FIGURE S17.** Comparison of sequencing length profiles for naturally occurring sRNAs found in the representative D (D2) plasma RNA sample between libraries prepared by protocols T[1+2] and T[1+2+3+4] for the indicated RNA classes: (**A**) rRNA, (**B**) miRNA, (**C**) tRNA, (**D**) snRNA, (**E**) snoRNA, (**F**) mRNA, (**G**) lncRNA and (**H**) piRNA.

**Supplementary FIGURE S18.** Sequencing length and RNA Type profiles for naturally occurring sRNAs of the indicated RNA classes found in human CL (CL1, CL2, CL4) plasma RNA samples for libraries prepared by (**A**) RiboMarker® protocol T[1+2+3+4] or (**B**) Phospho-RNA-Seq.

**Supplementary FIGURE S19.** Comparison of sequencing length profiles of naturally occurring sRNAs of the indicated RNA classes found in the representative human CL (CL4) plasma RNA sample between libraries prepared by (**A**) RiboMarker® protocol T[1+2+3+4] or (**B**) Phospho-RNA-Seq.

**Supplementary FIGURE S20.** Percentages of normalized sequencing reads for the main sRNA classes found in the human CL (CL1, CL2, CL4) plasma RNA samples among libraries prepared by the indicated RiboMarker® protocols and Phospho-RNA-Seq.

**Supplementary FIGURE S21.** Sequencing length and RNA Type profiles for stsRNAs spiked in total RNA isolated from either human H (H1, H2, H3; left panels) plasma or D (D1, D2, D3; right panels) plasma RNA samples for the libraries prepared by protocols (**A**) T[3+4], (**B**) T[3], or (**C**) T[4] .

**Supplementary FIGURE S22.** Sequencing length and RNA Type profiles for naturally occurring sRNAs in total RNA isolated from either human H (H1, H2, H3; left panels) plasma or D (D1, D2, D3; right panels) plasma RNA samples for libraries prepared by protocols (**A**) T[3+4], (**B**) T[3], or (**C**) T[4]; and (**D**) a comparison of profiles for the representative human H (H3) and D (D3) plasma samples prepared by protocols T[3+4] (top panels), T[3] (middle panels) or T[4] (bottom panels).

**Supplementary FIGURE S23.** Comparison of sequencing length profiles for naturally occurring sRNAs for the representative human H (H3) and D (D2) plasma RNA samples between libraries prepared by protocols T[3+4], T[3], and T[4] for the following RNA classes: (**A**) rRNA, (**B**) tRNA, (**C**) miRNA, (**D**) snRNA, (**E**) snoRNA, (**F**) mRNA, (**G**) lncRNA and (**H**) piRNA.

**Supplementary FIGURE S24.** Hierarchically clustered heatmap showing the relative abundance (row-wise z-score) of significantly variable RNA transcripts and/or fragments among RiboMarker® protocol T[1+2+3+4] and Phospho-RNA-Seq (DESeq2 LRT; padj < 0.05) from human CL (CL1, CL2, CL4) plasma RNA samples. Darker patterns indicate a higher relative abundance of corresponding RNA transcripts. The side color lines denote a higher relative enrichment of certain RNA sequences within the indicated RNA classes by RiboMarker® protocol T[1+2+3+4] (orange lines) or by Phospho-RNA-Seq (purple lines).

**Supplementary FIGURE S25.** Top 20 most abundant sRNA sequencing reads (5'-3') aligned to the RNU2-1 snRNA that were detected in the selected human D (D1, D2) plasma RNA samples for libraries prepared by protocol [T2]. The underlined A and B areas of the RNU2-1 transcript sequences correspond to the highlighted areas in Fig. 11C.

**Supplementary FIGURE S1A**

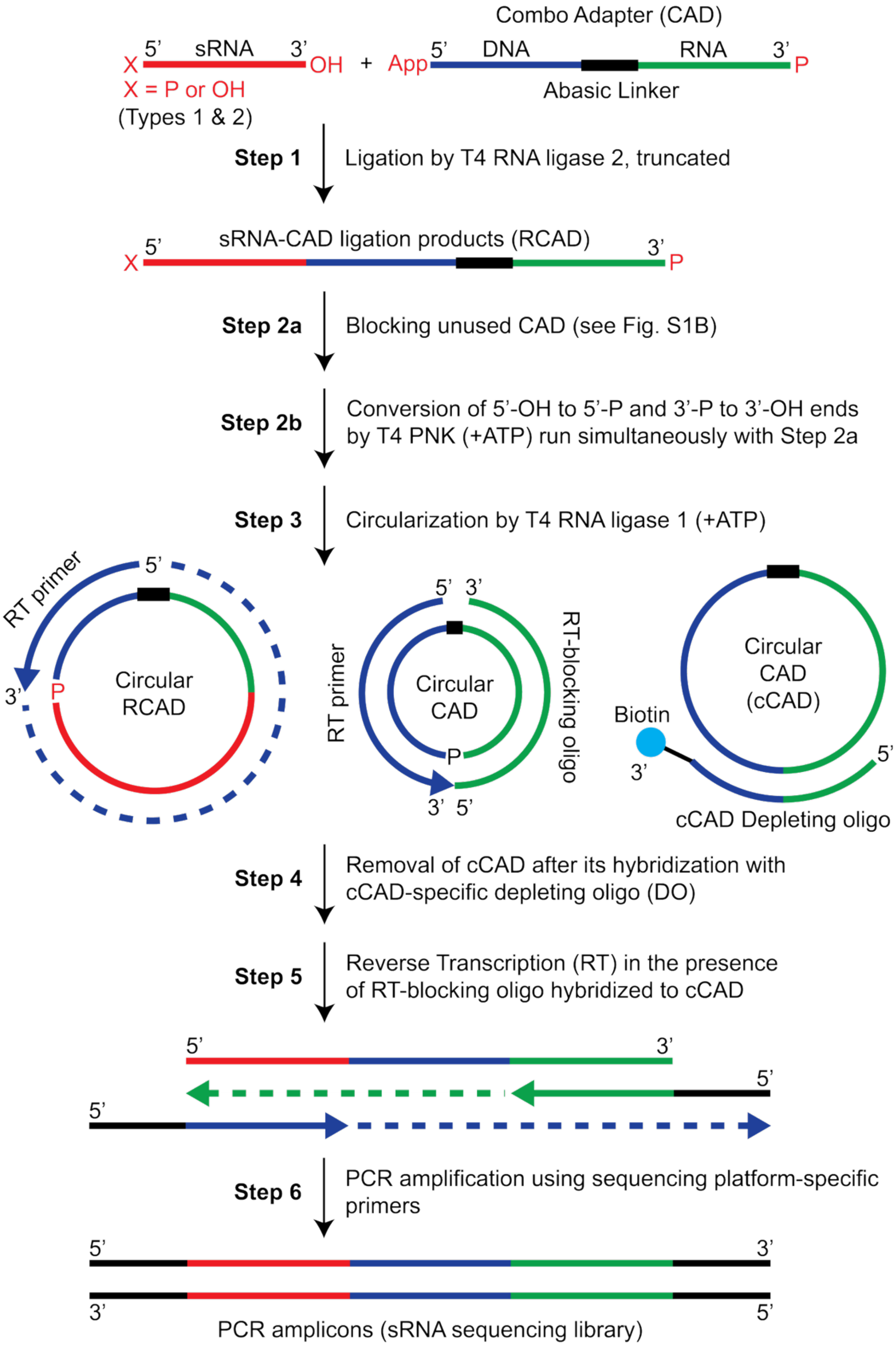

**Supplementary FIGURE S1B**

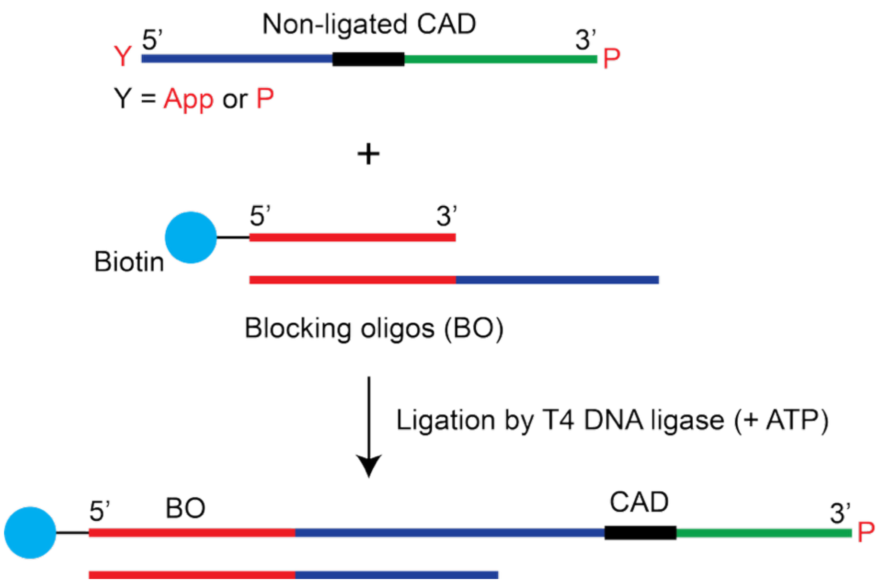

**Supplementary FIGURE S2**

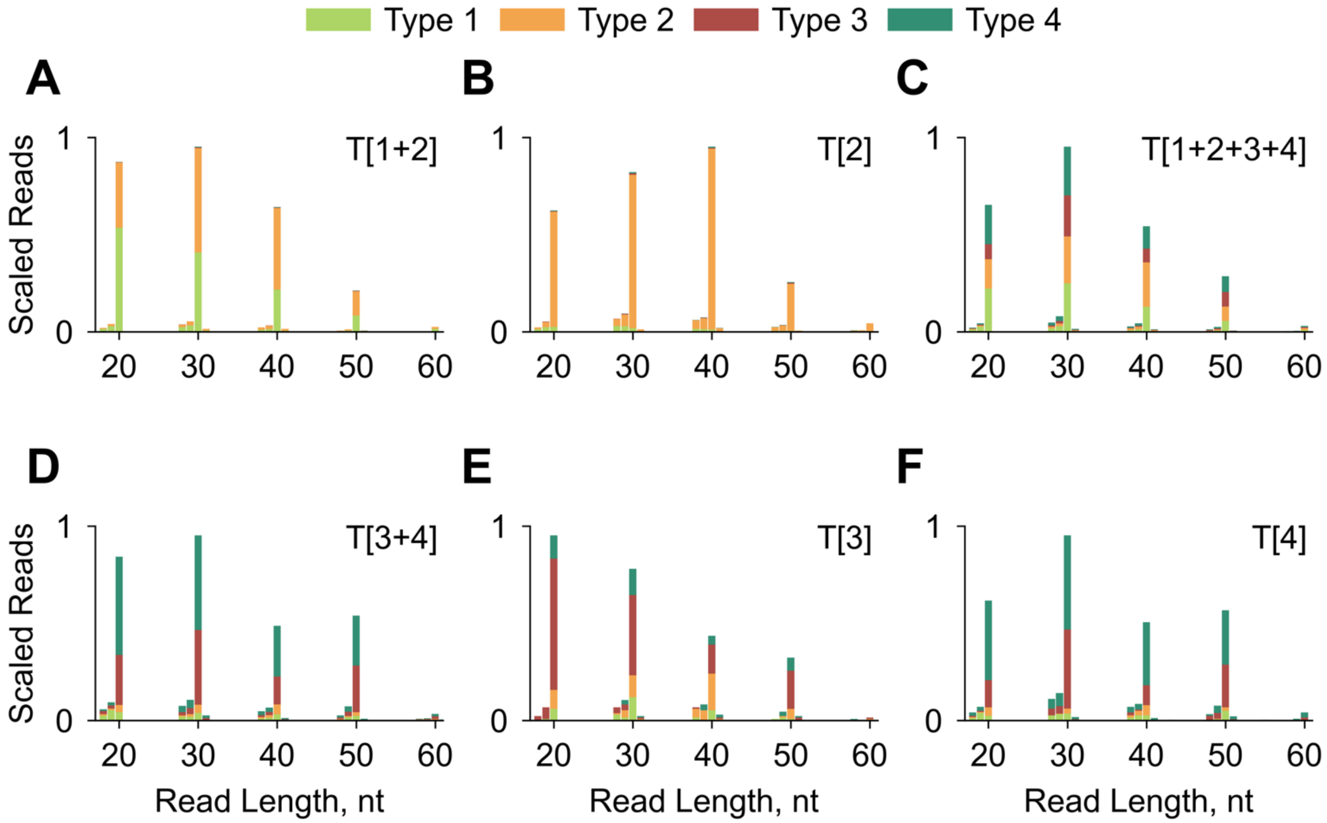

Supplementary FIGURE S3

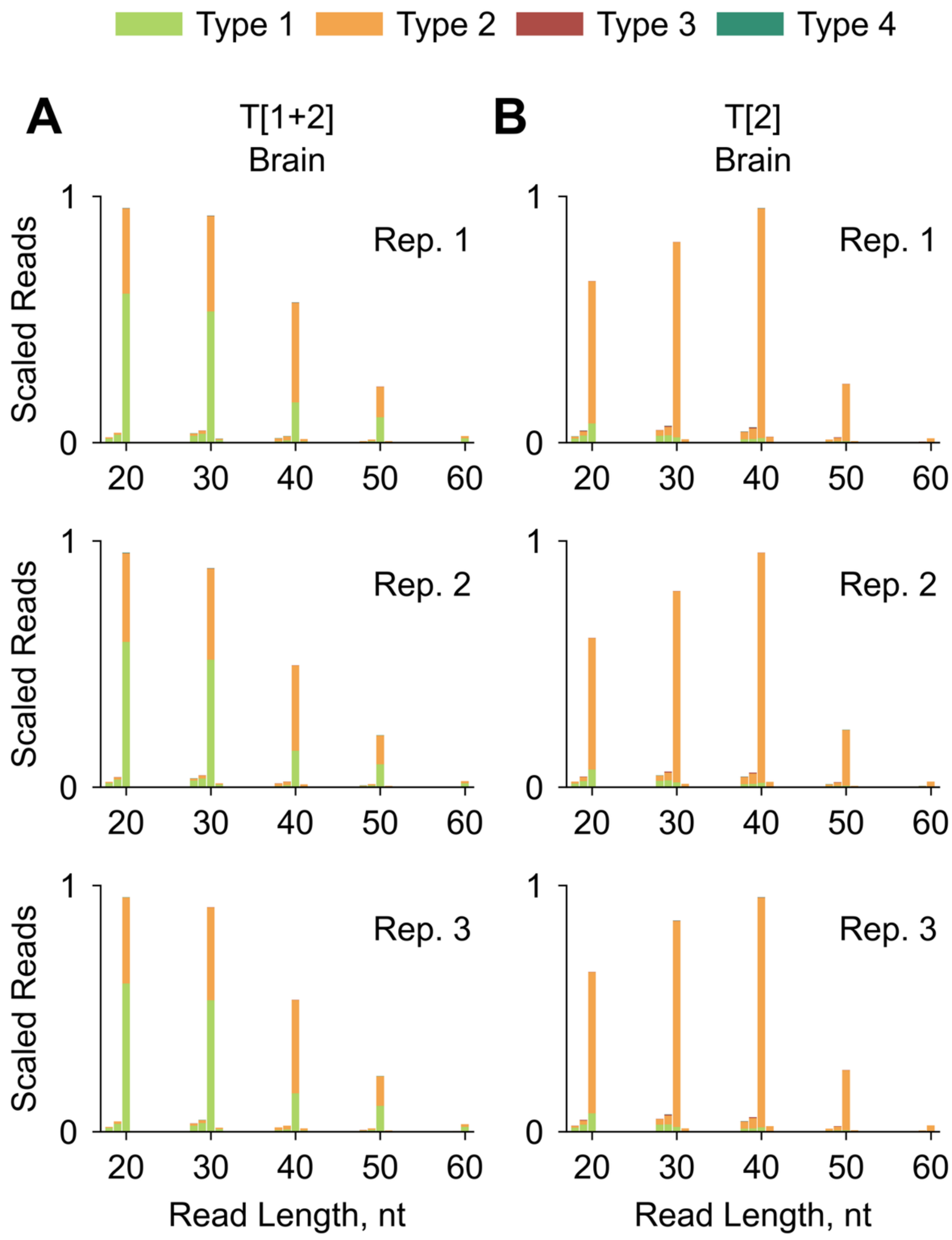

**Supplementary FIGURE S4**

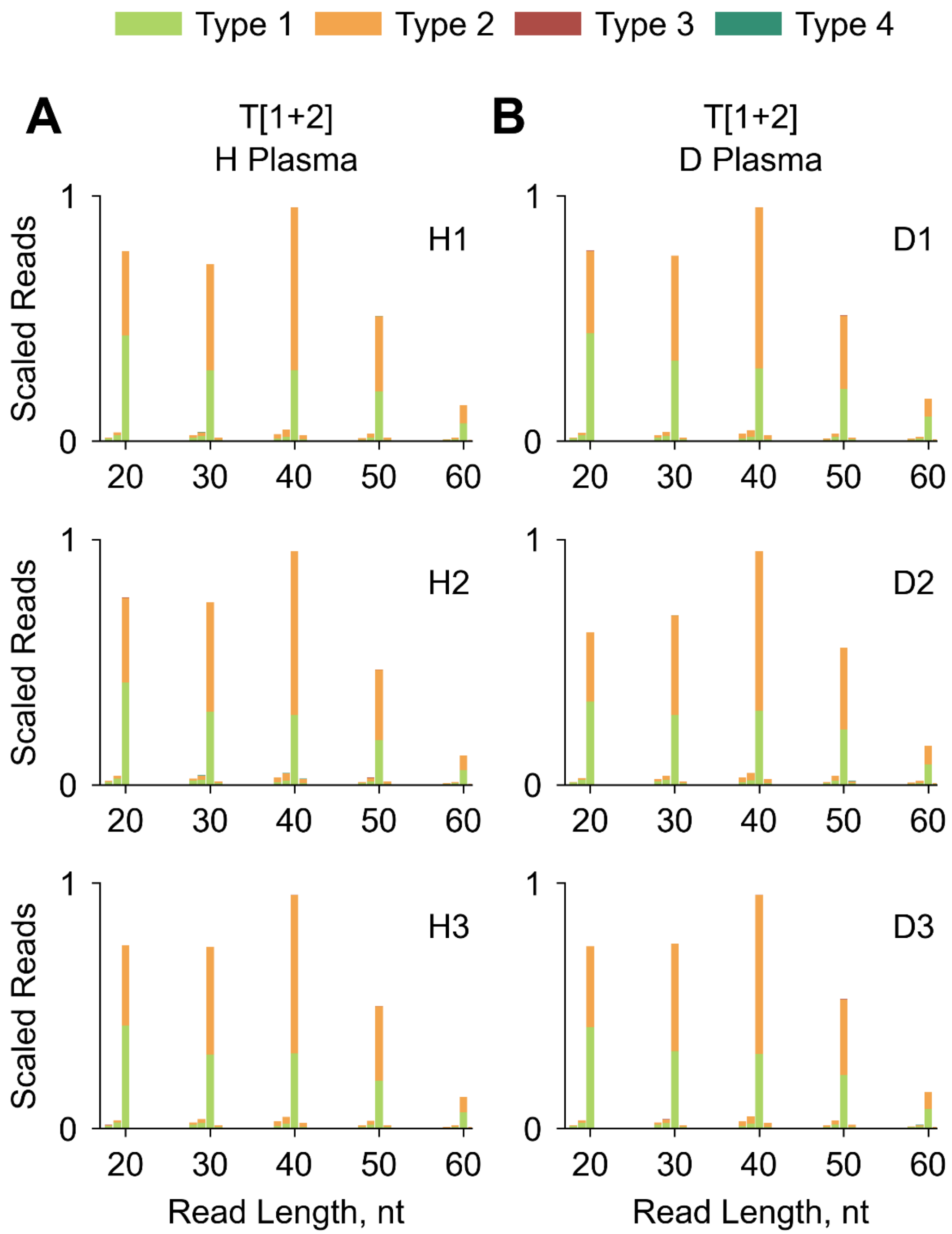

**Supplementary FIGURE S5**

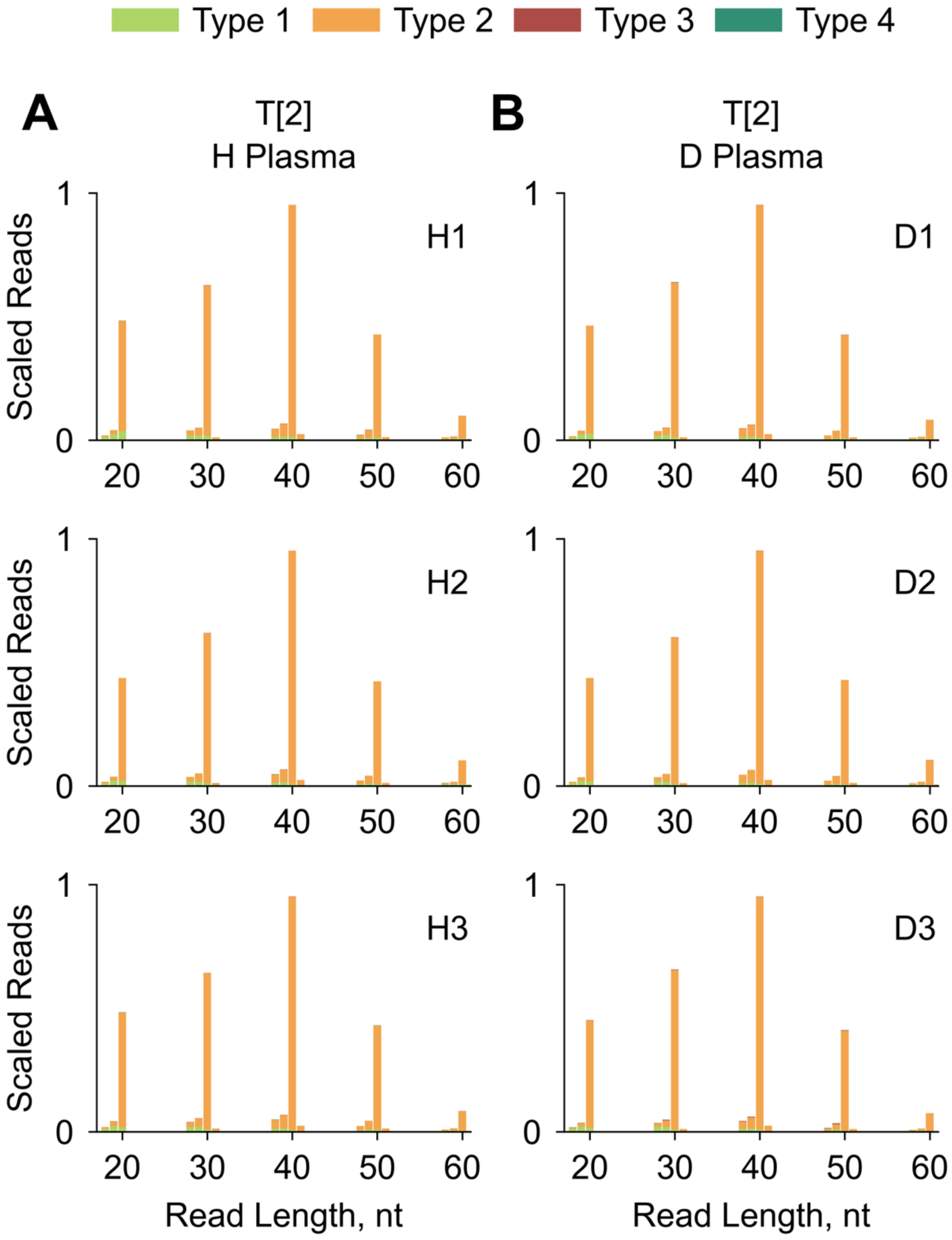

Supplementary FIGURE S6

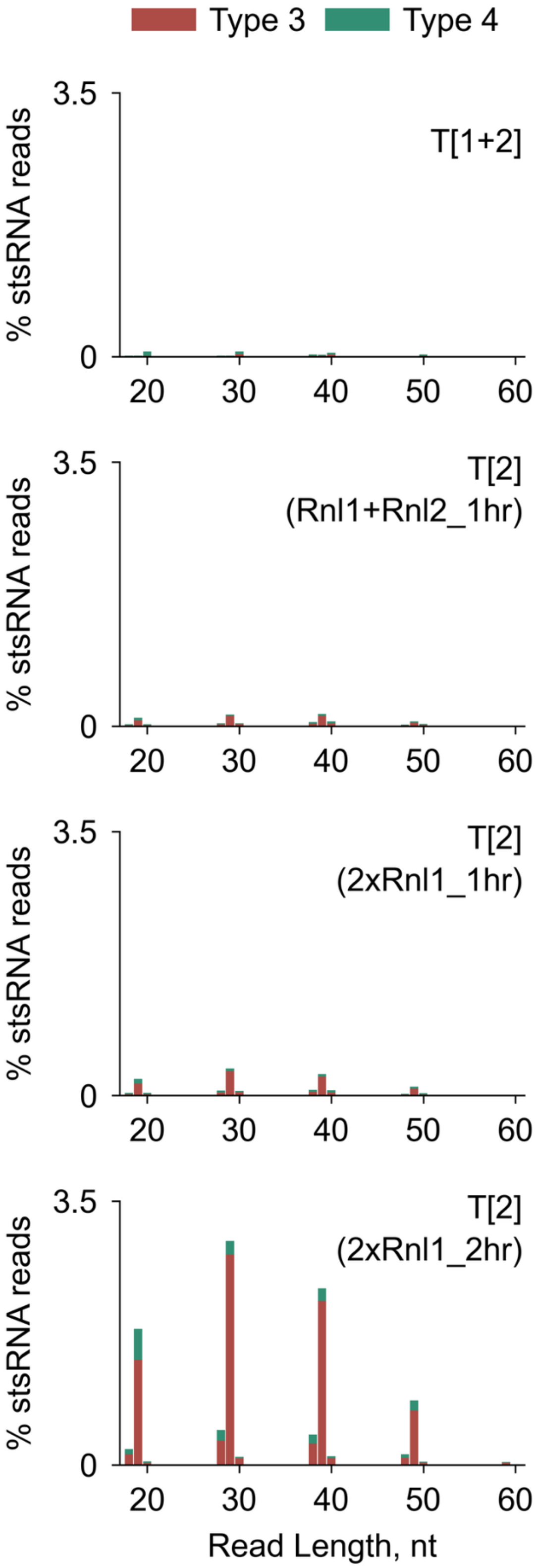

**Supplementary FIGURE S7**

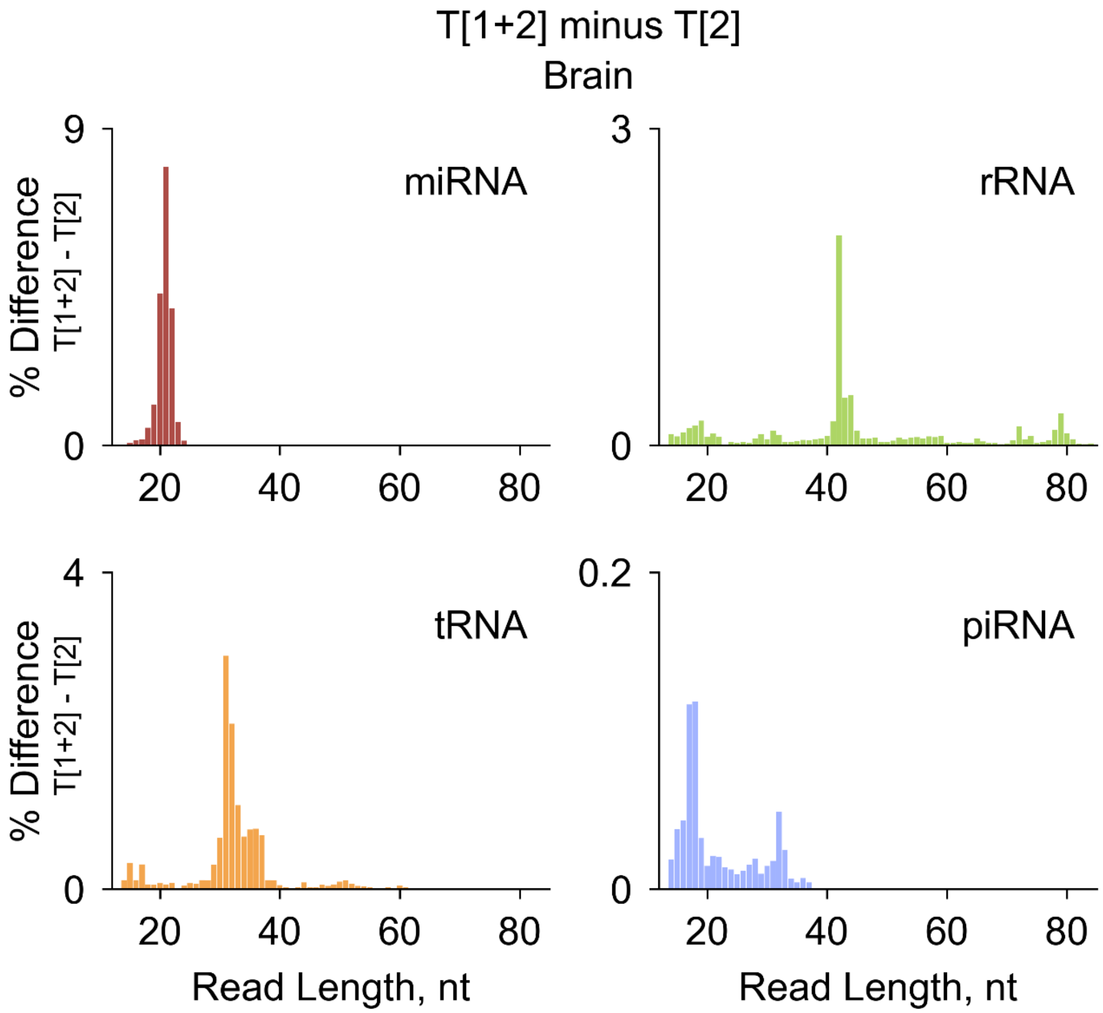

**Supplementary FIGURE S8**

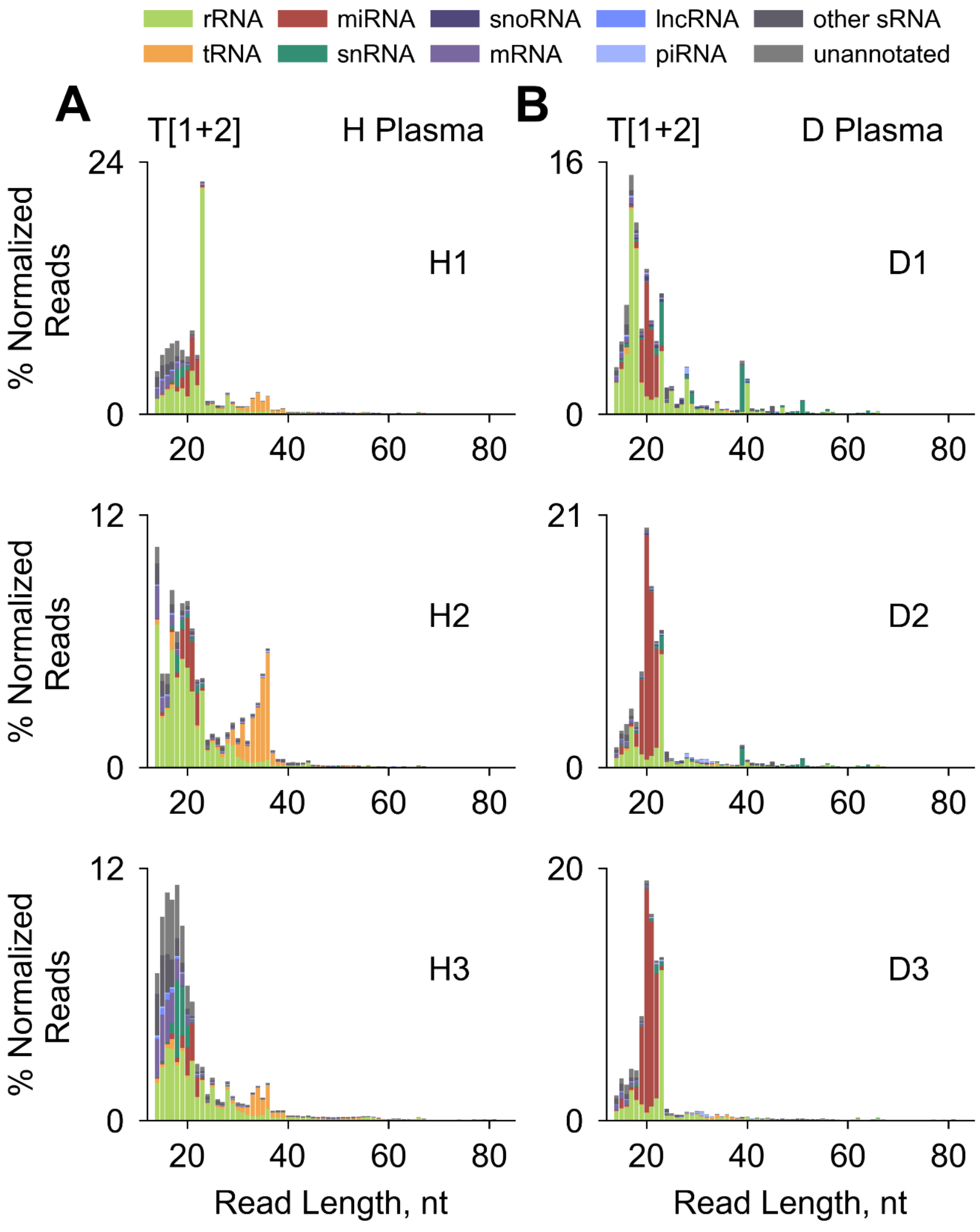

**Supplementary FIGURE S9**

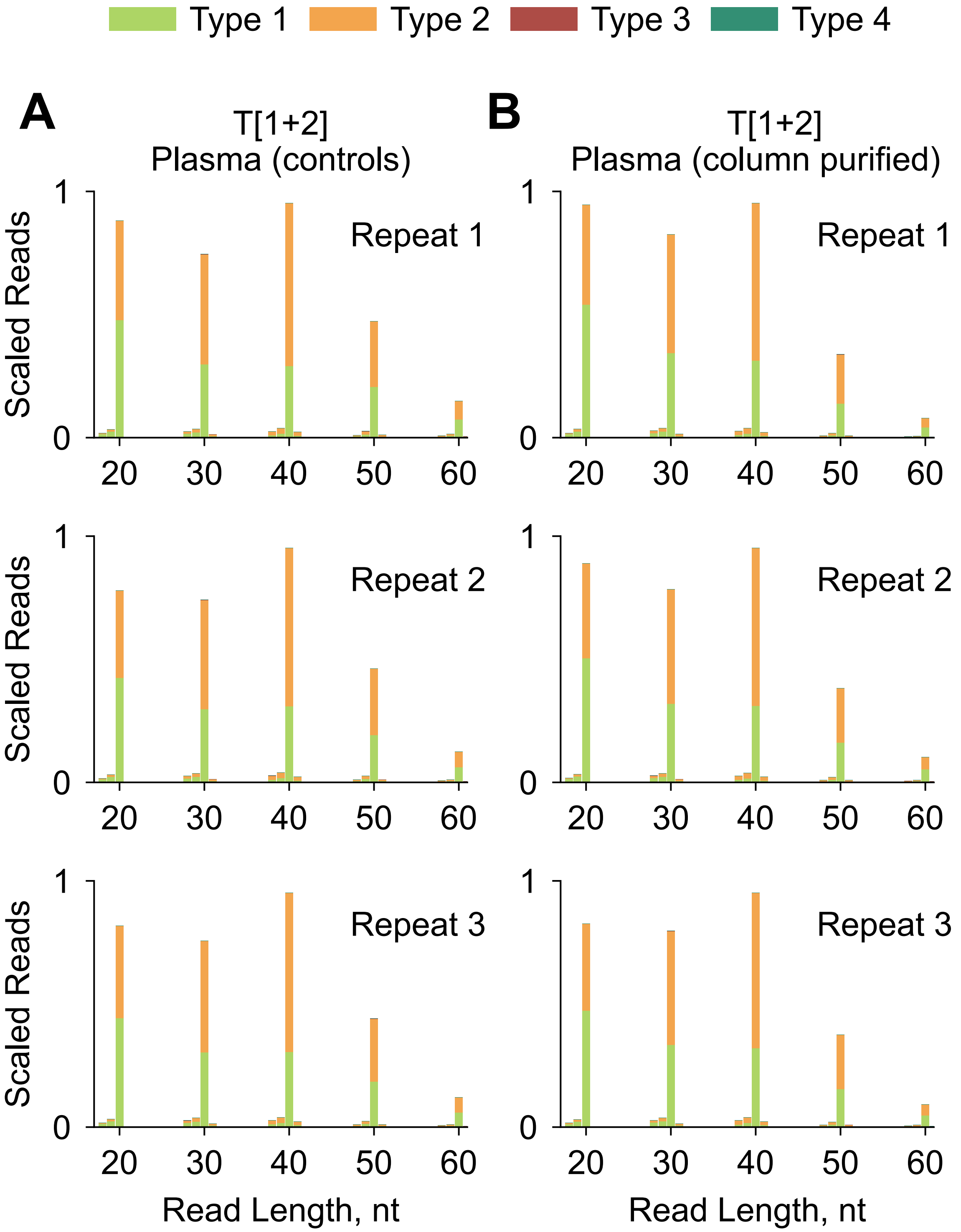

**Supplementary FIGURE S10**

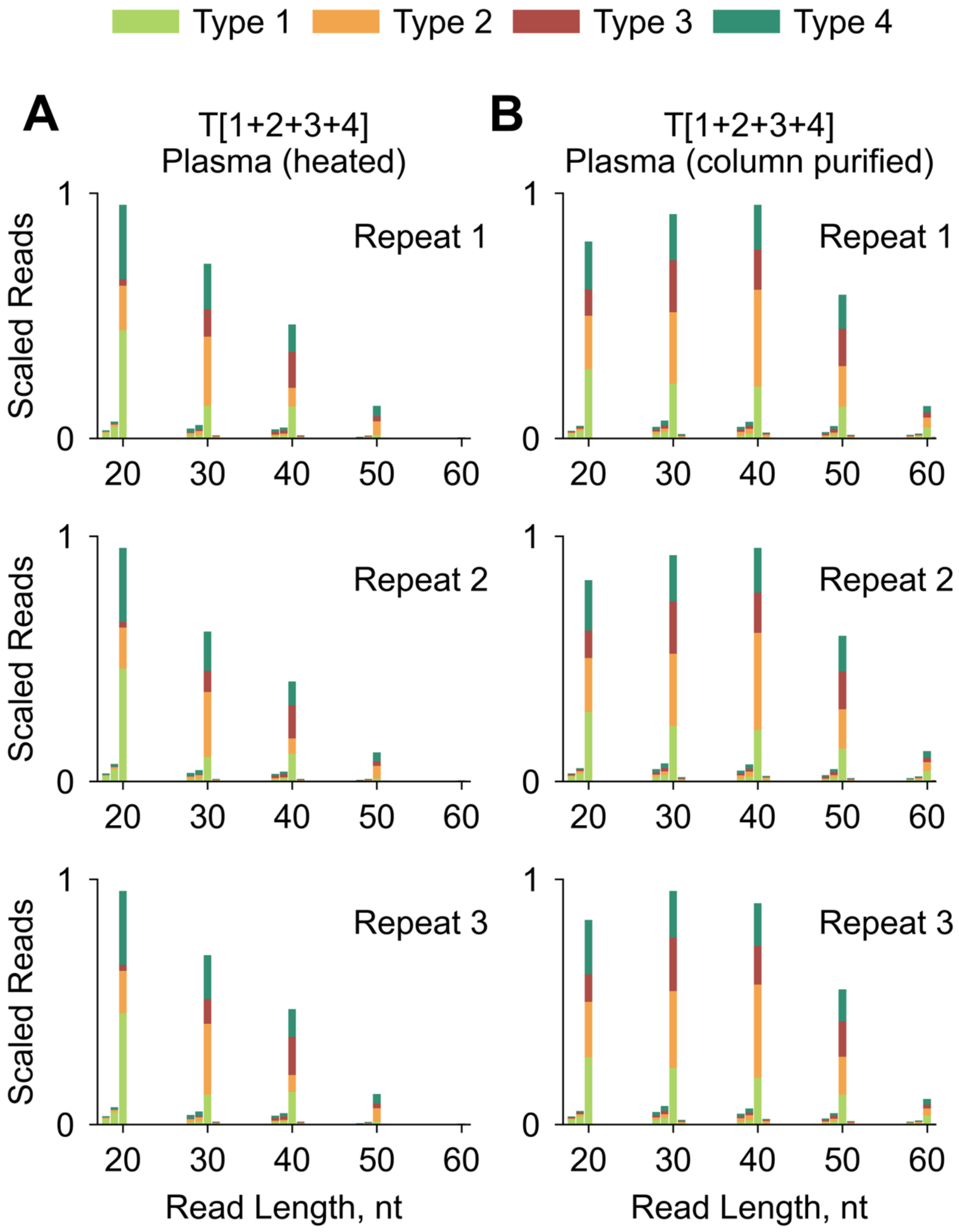

Supplementary FIGURE S11

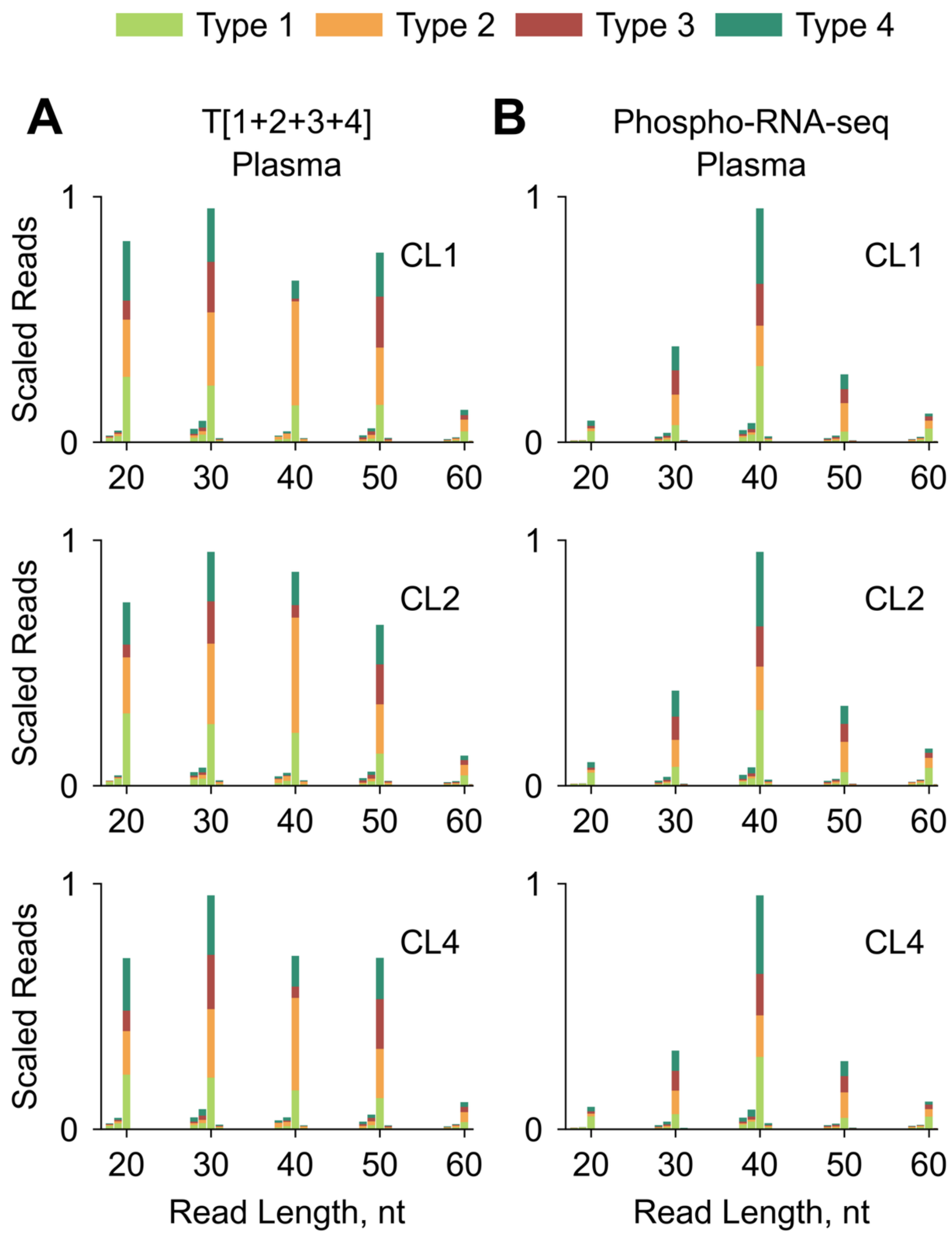

**Supplementary FIGURE S12**

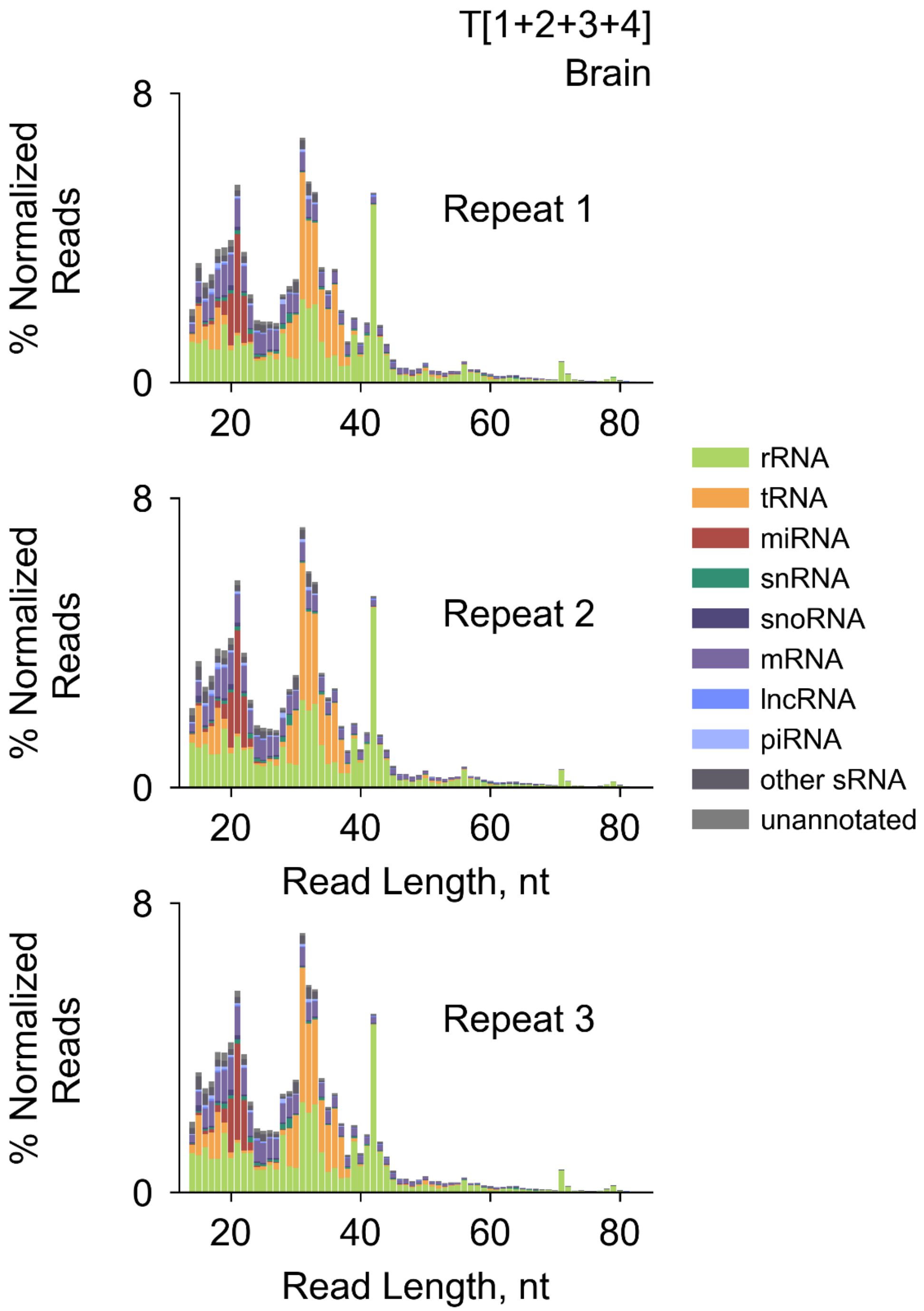

**Supplementary FIGURE S13**

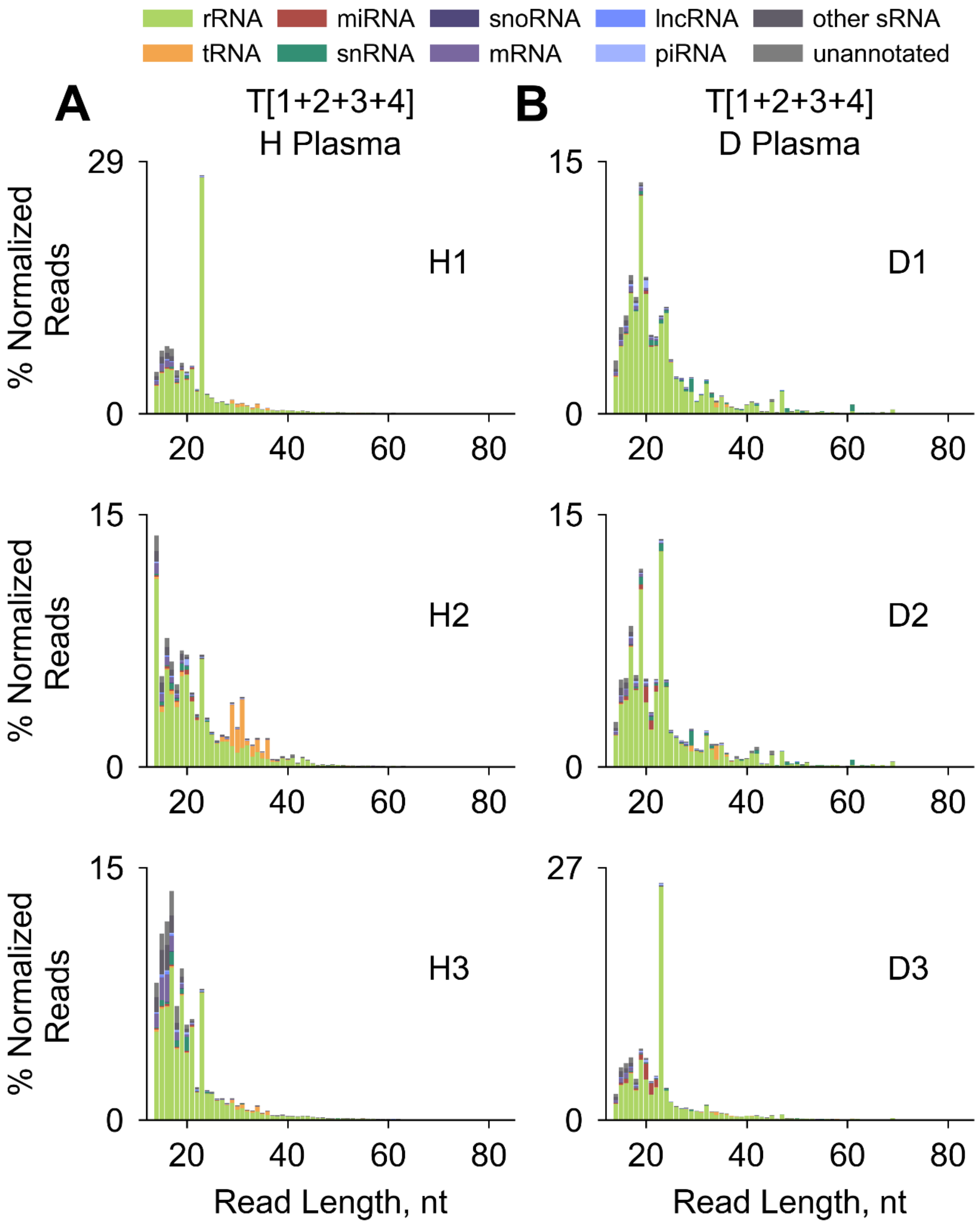

### Supplementary FIGURE S14

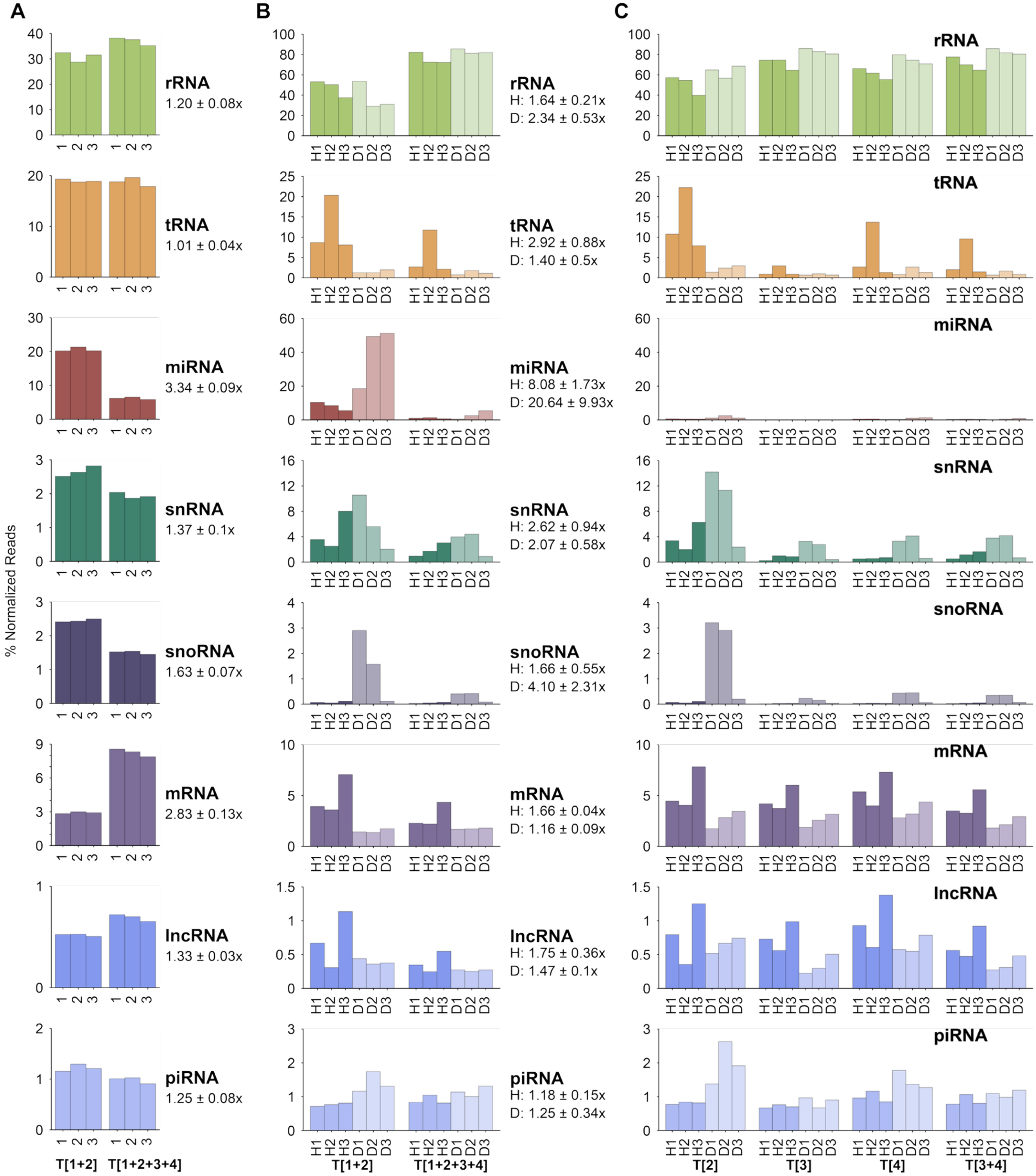

**Supplementary FIGURE S15**

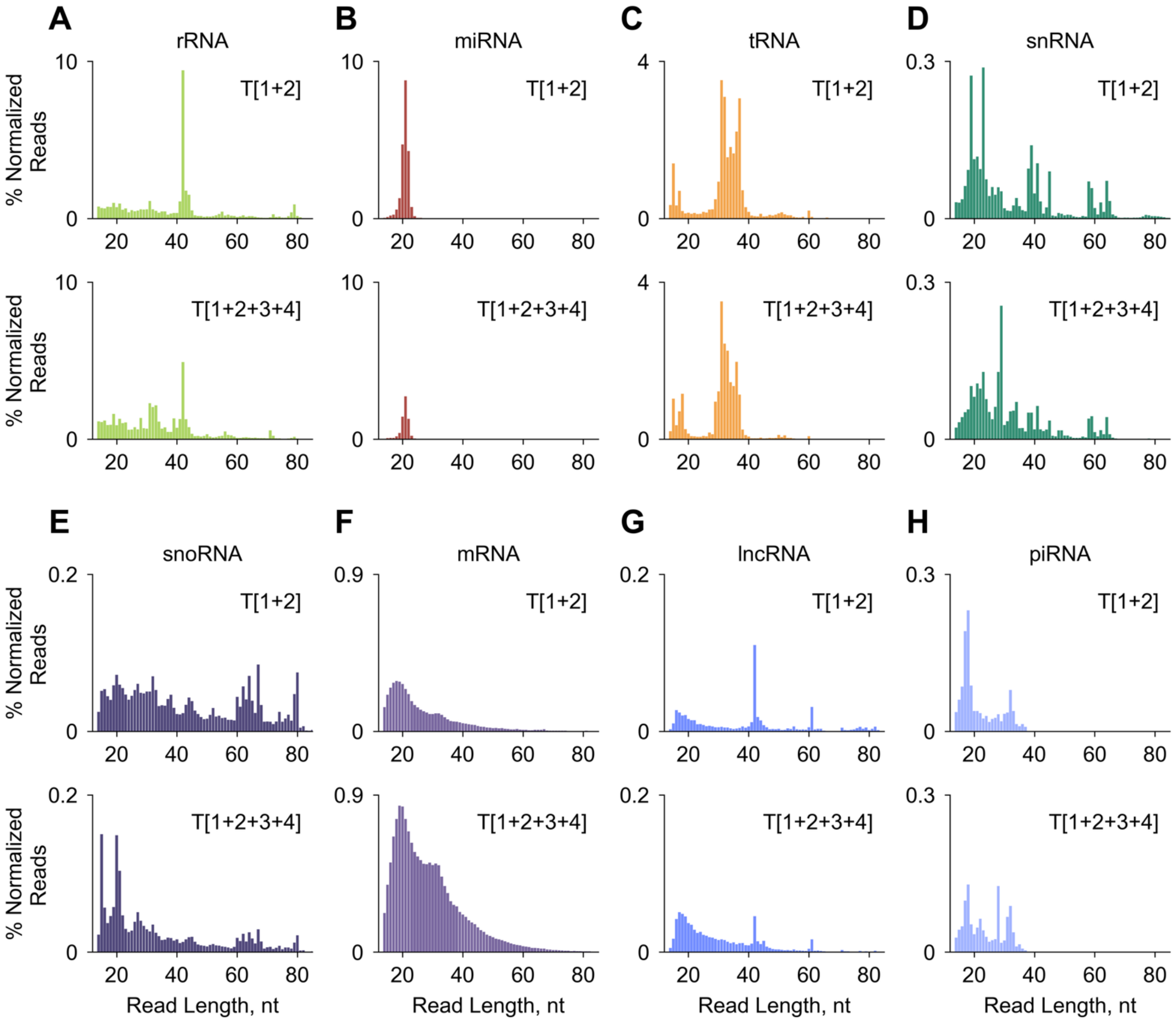

**Supplementary FIGURE S16**

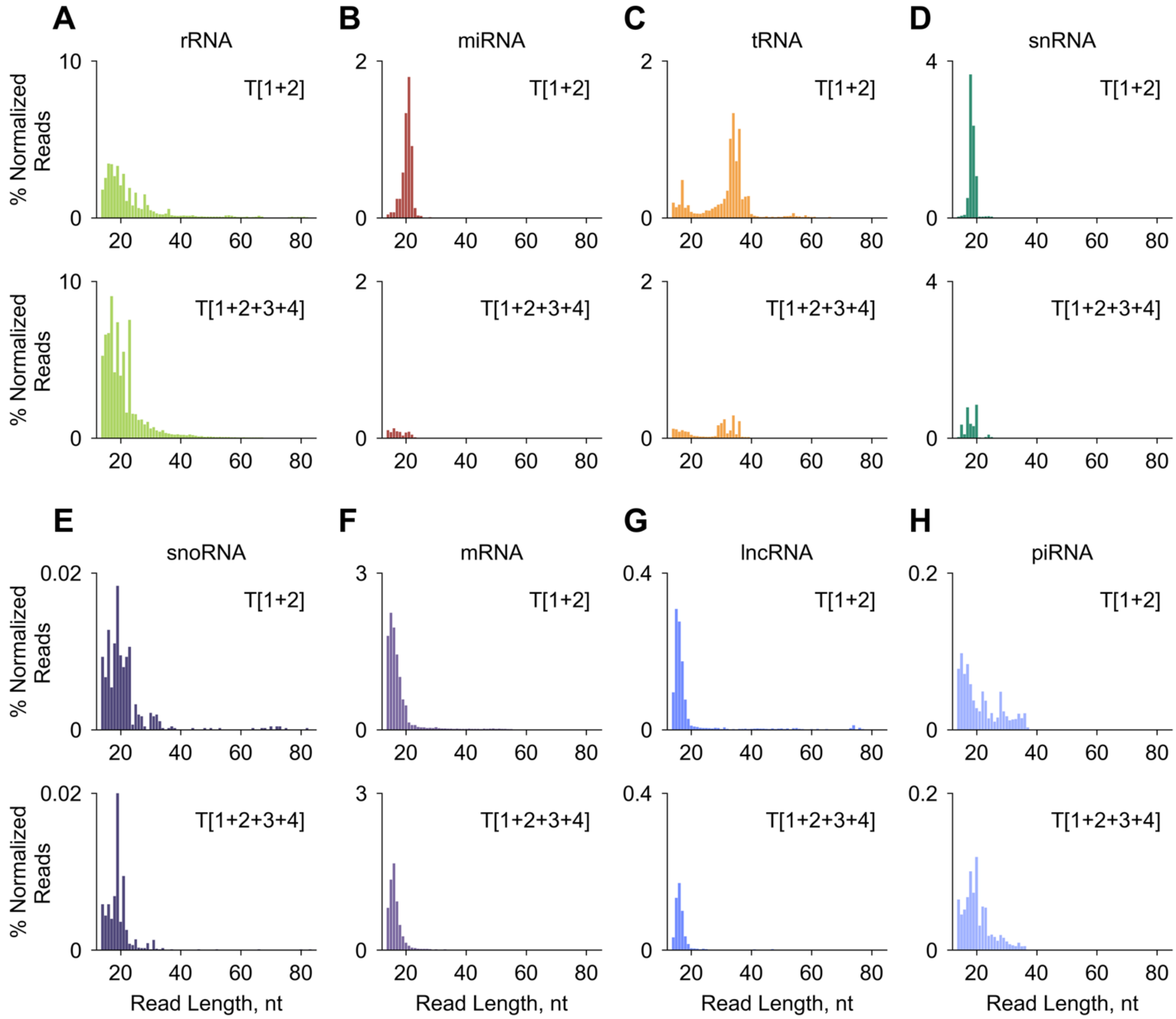

**Supplementary FIGURE S17**

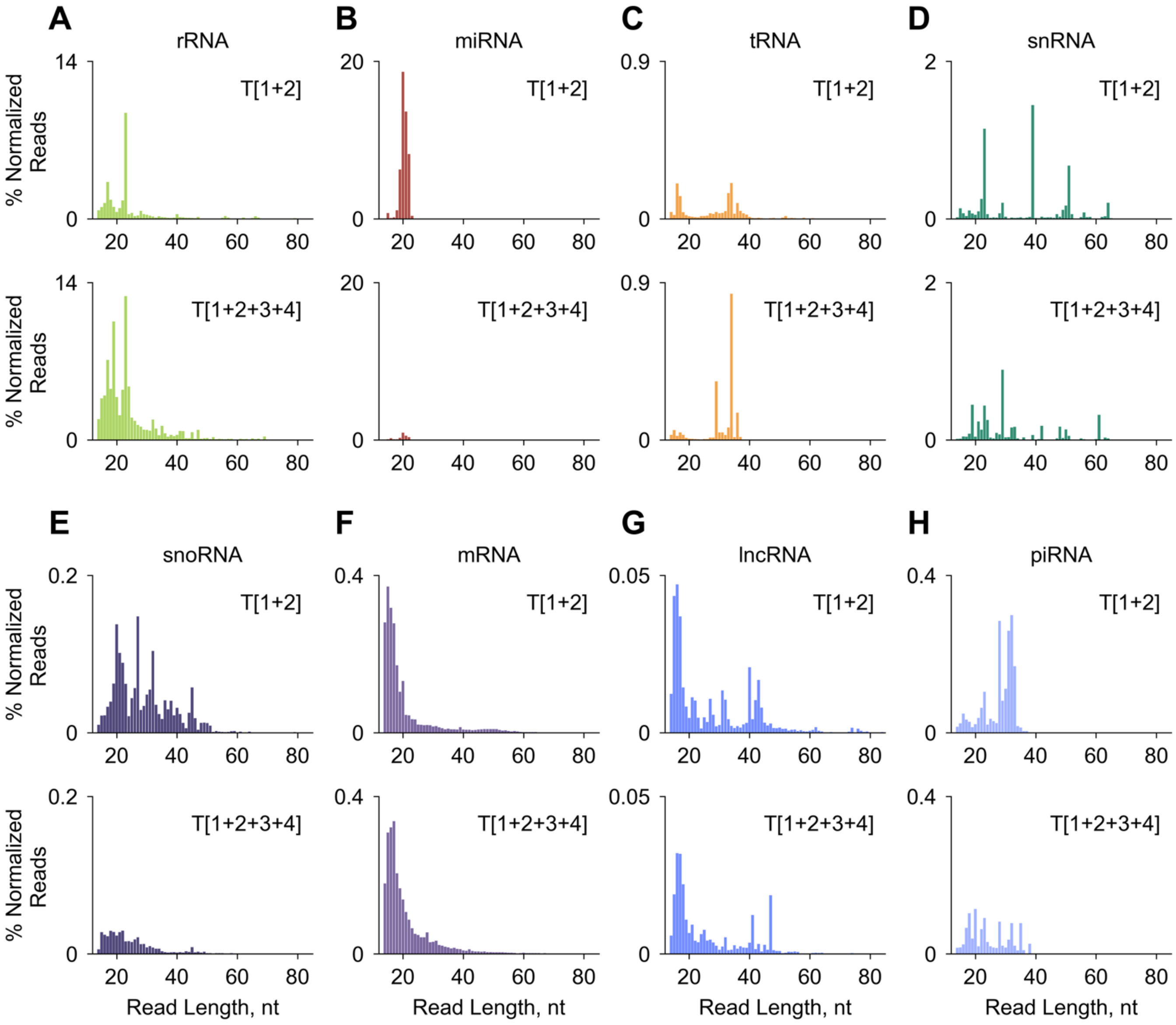

**Supplementary FIGURE S18**

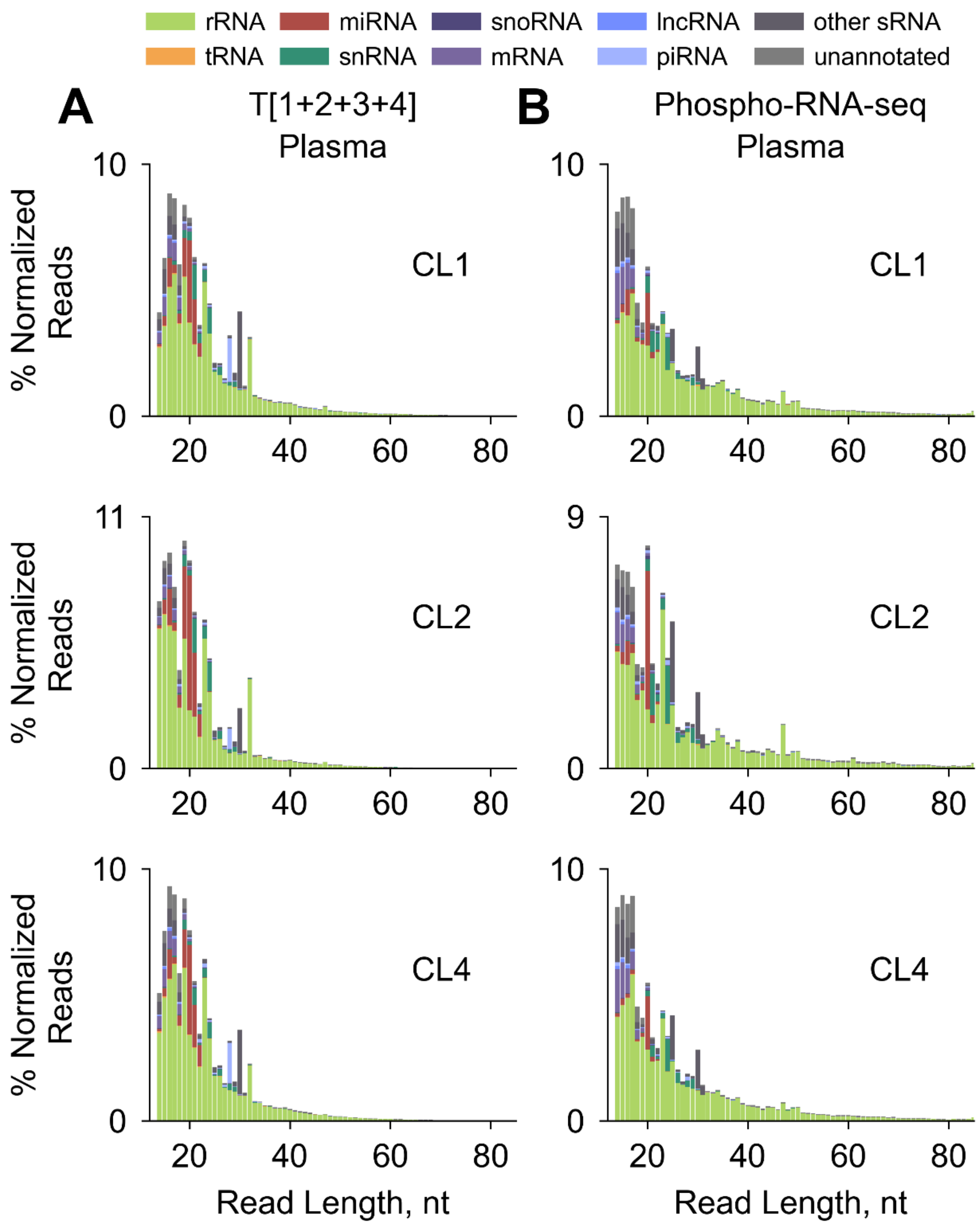

**Supplementary FIGURE S19**

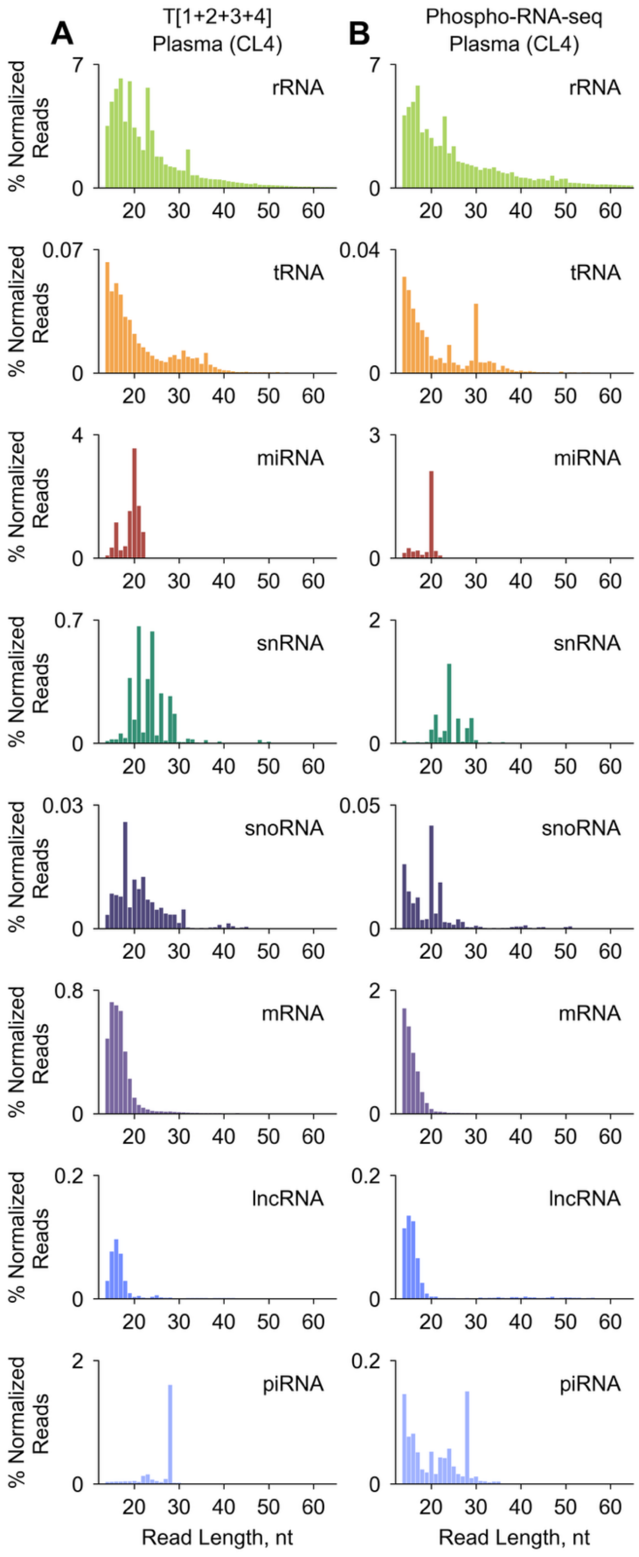

**Supplementary FIGURE S20**

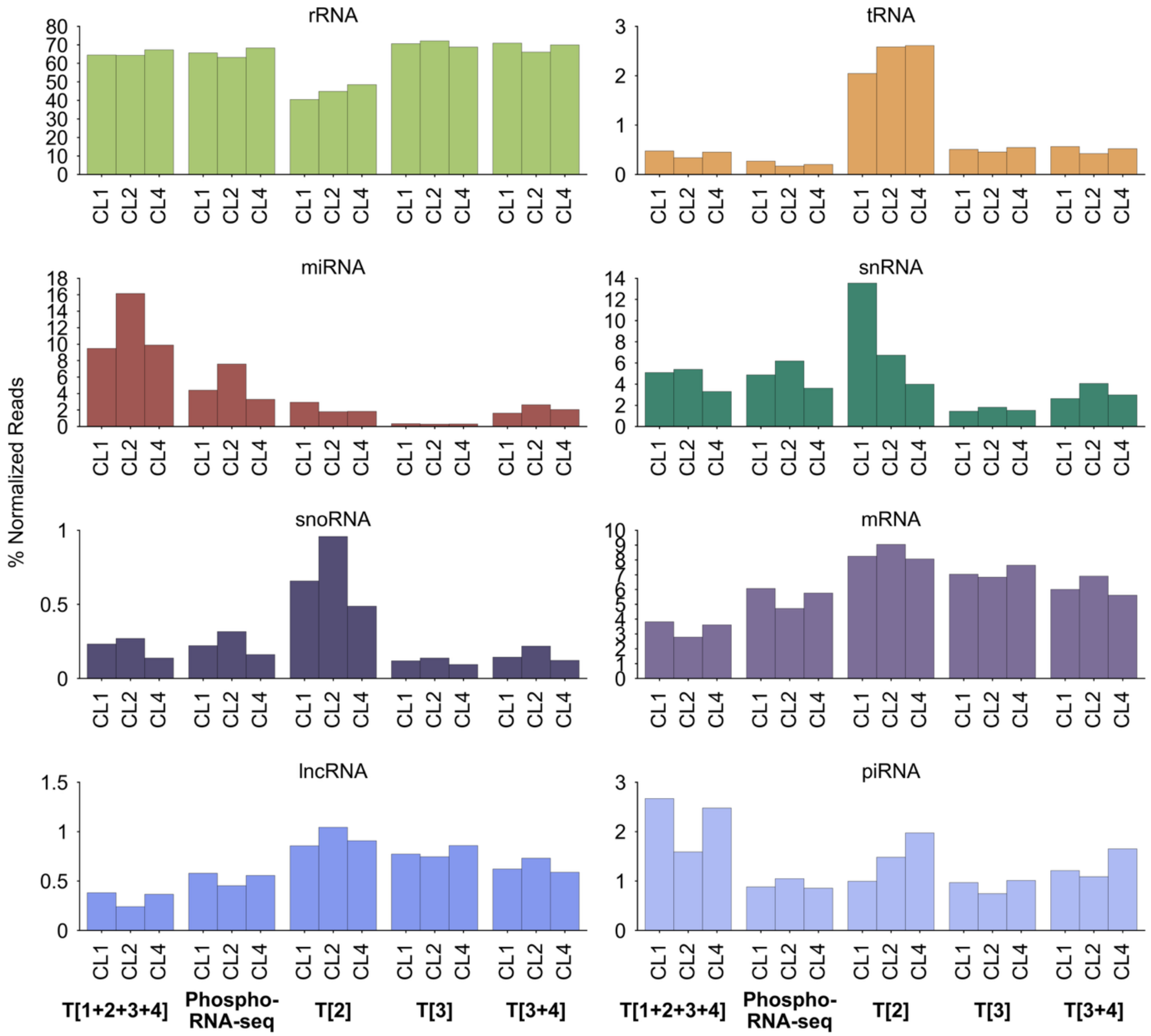

**Supplementary FIGURE S21A**

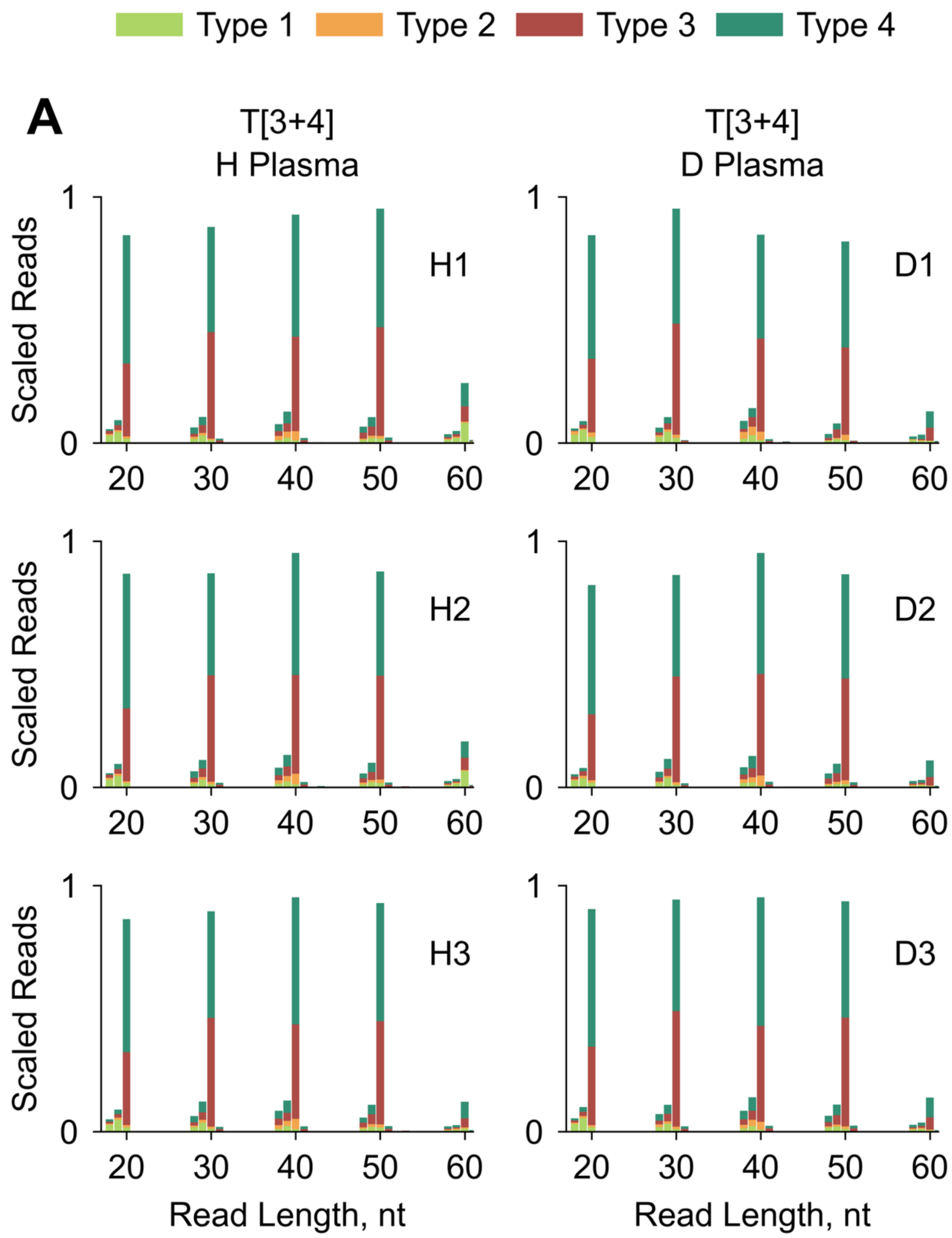

**Supplementary FIGURE S21B**

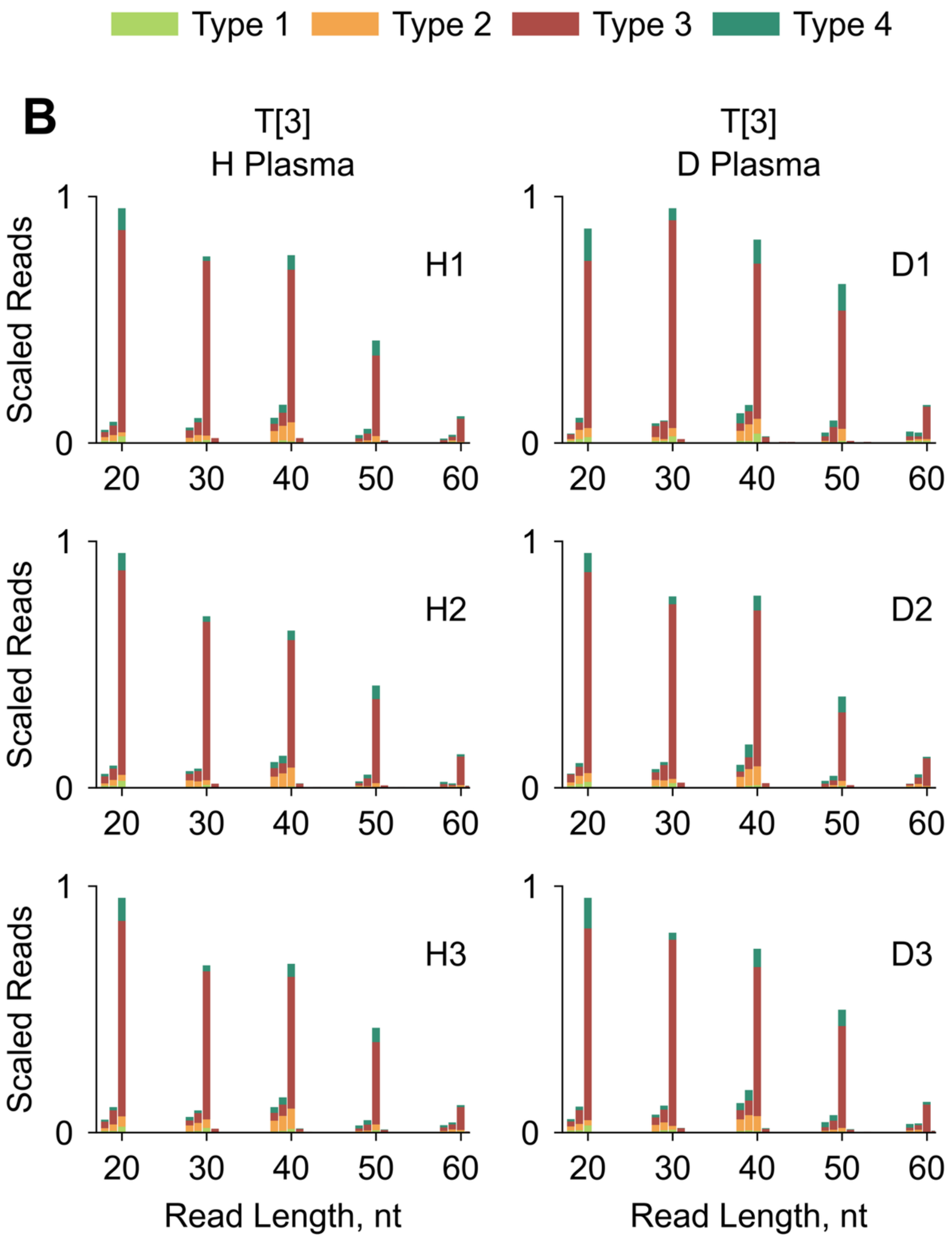

**Supplementary FIGURE S21C**

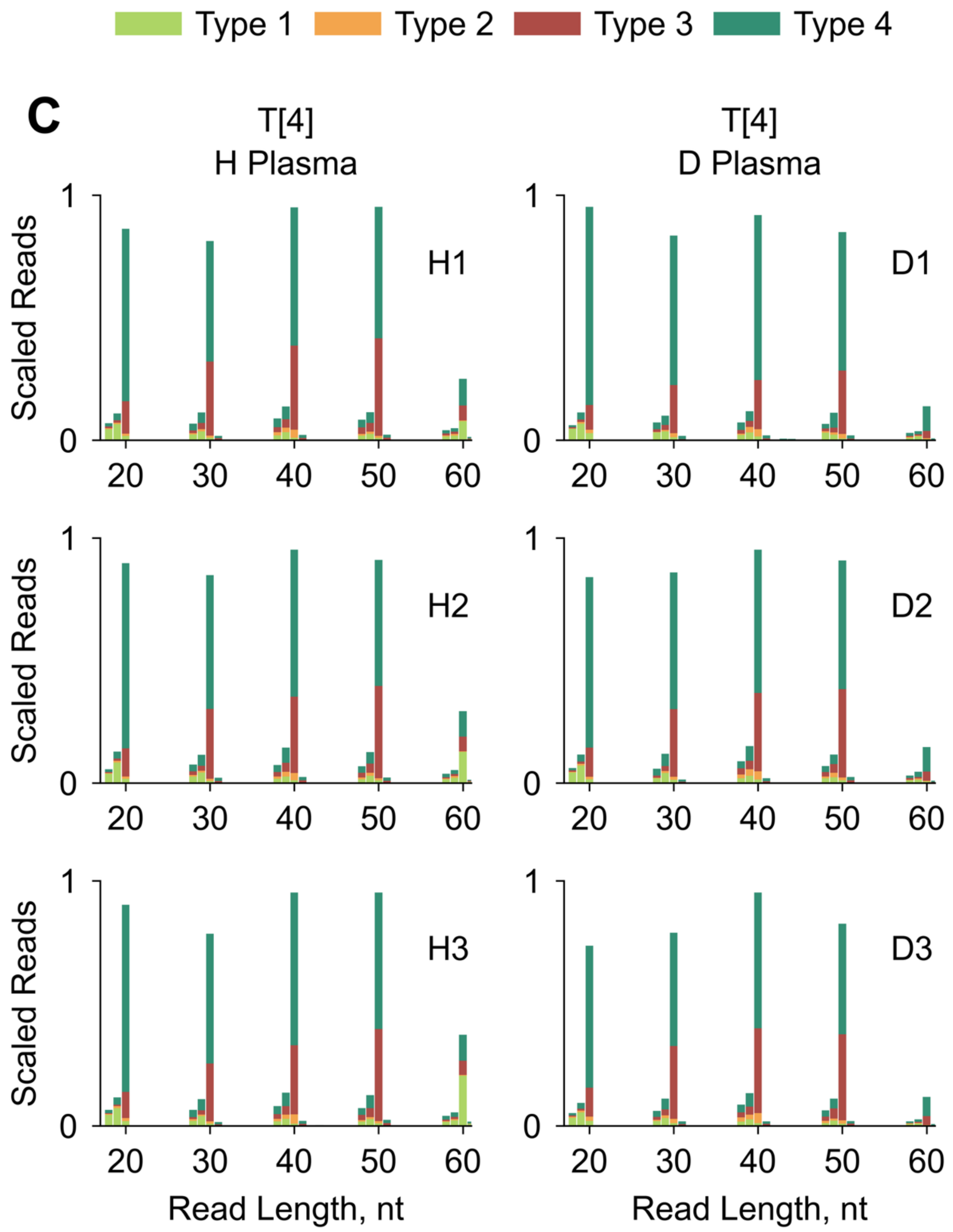

**Supplementary FIGURE S22A**

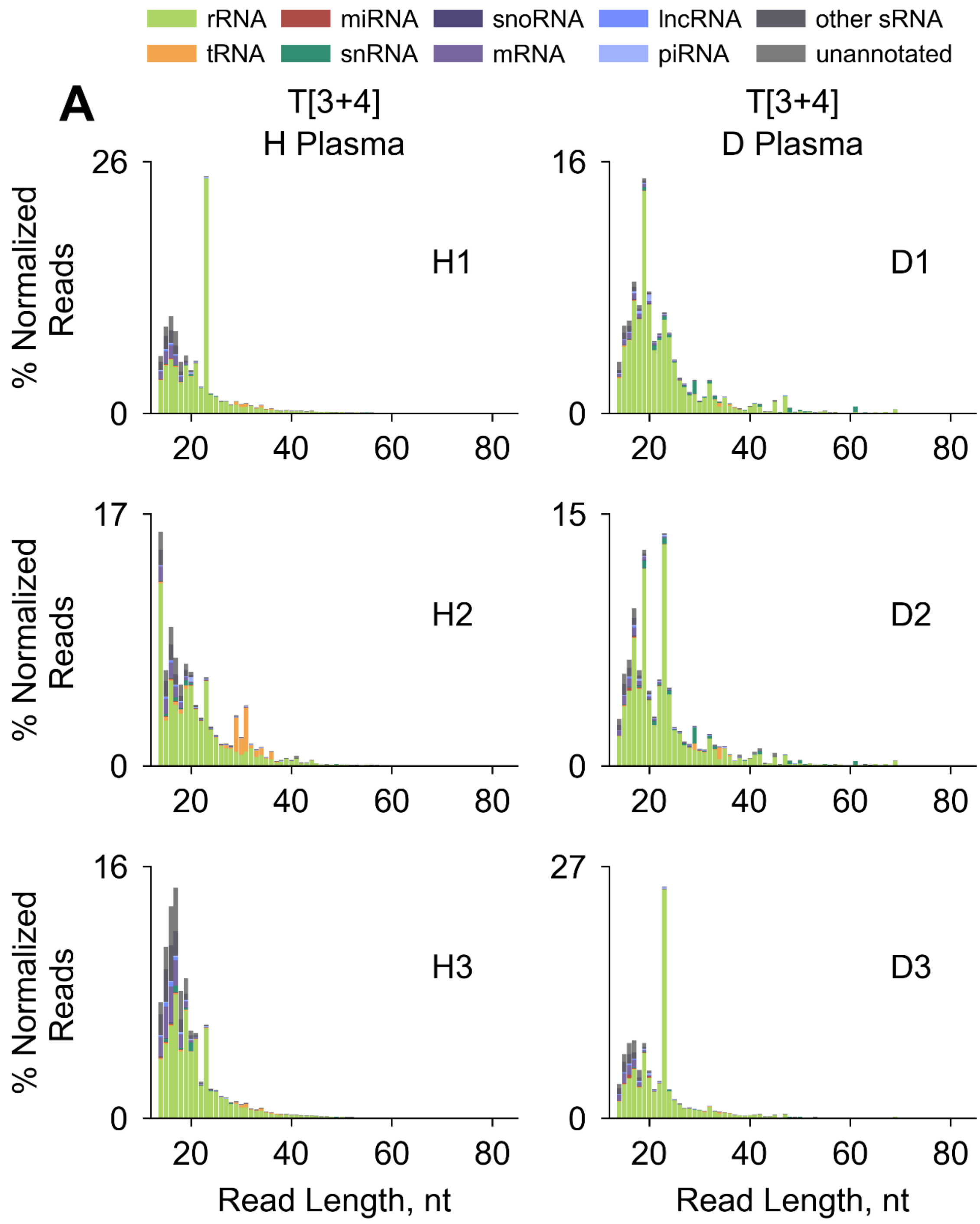

**Supplementary FIGURE S22B**

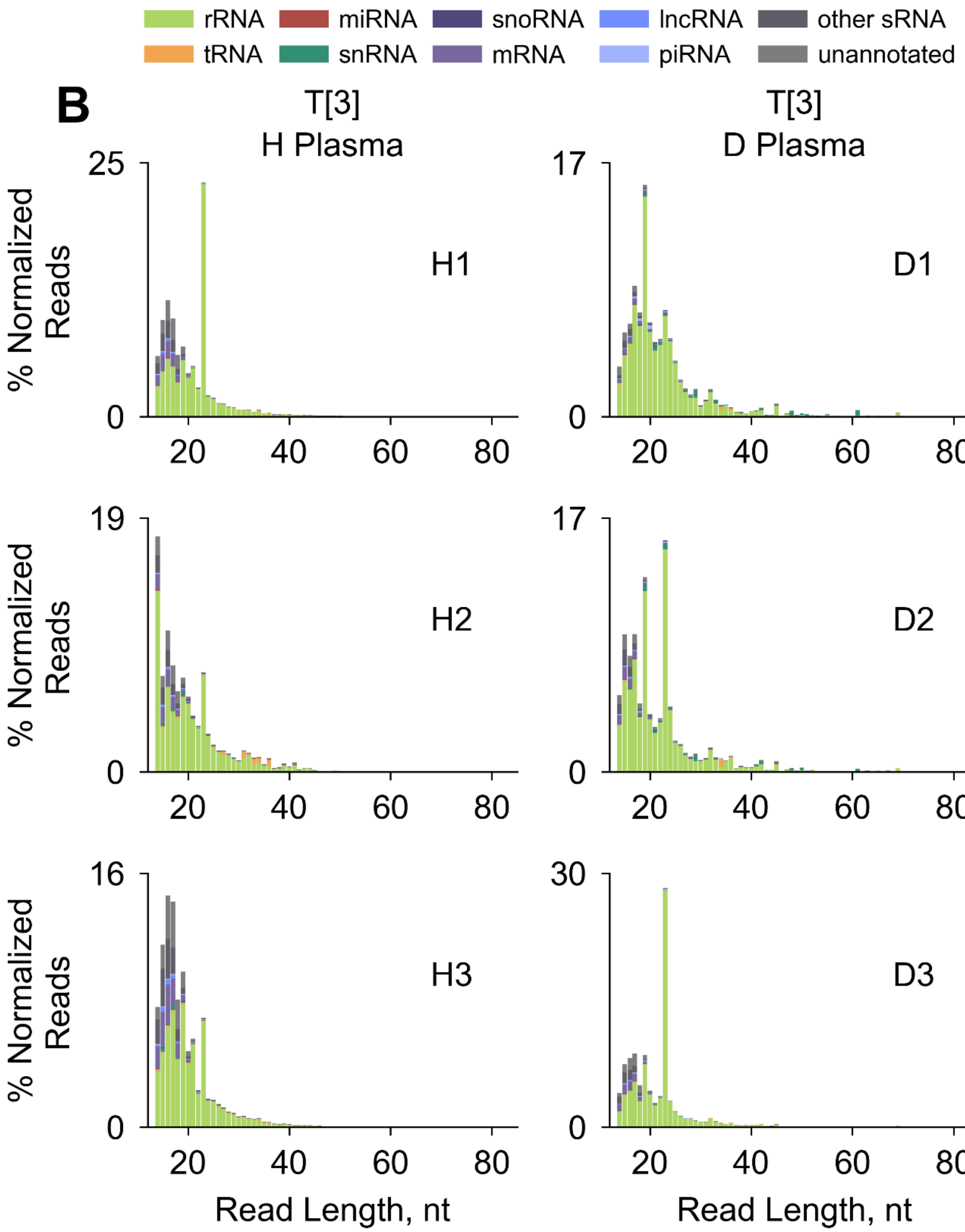

**Supplementary FIGURE S22C**

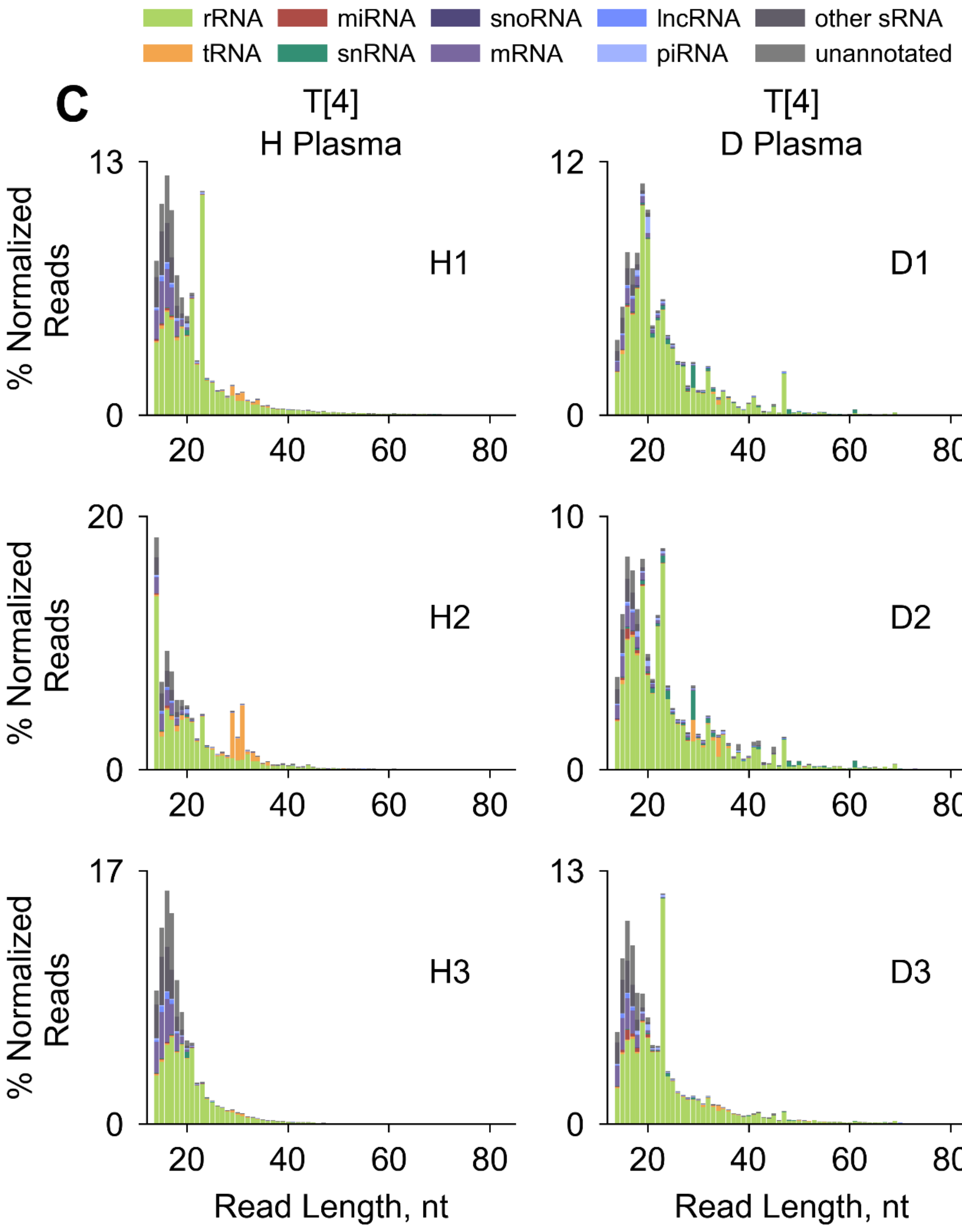

**Supplementary FIGURE S22D**

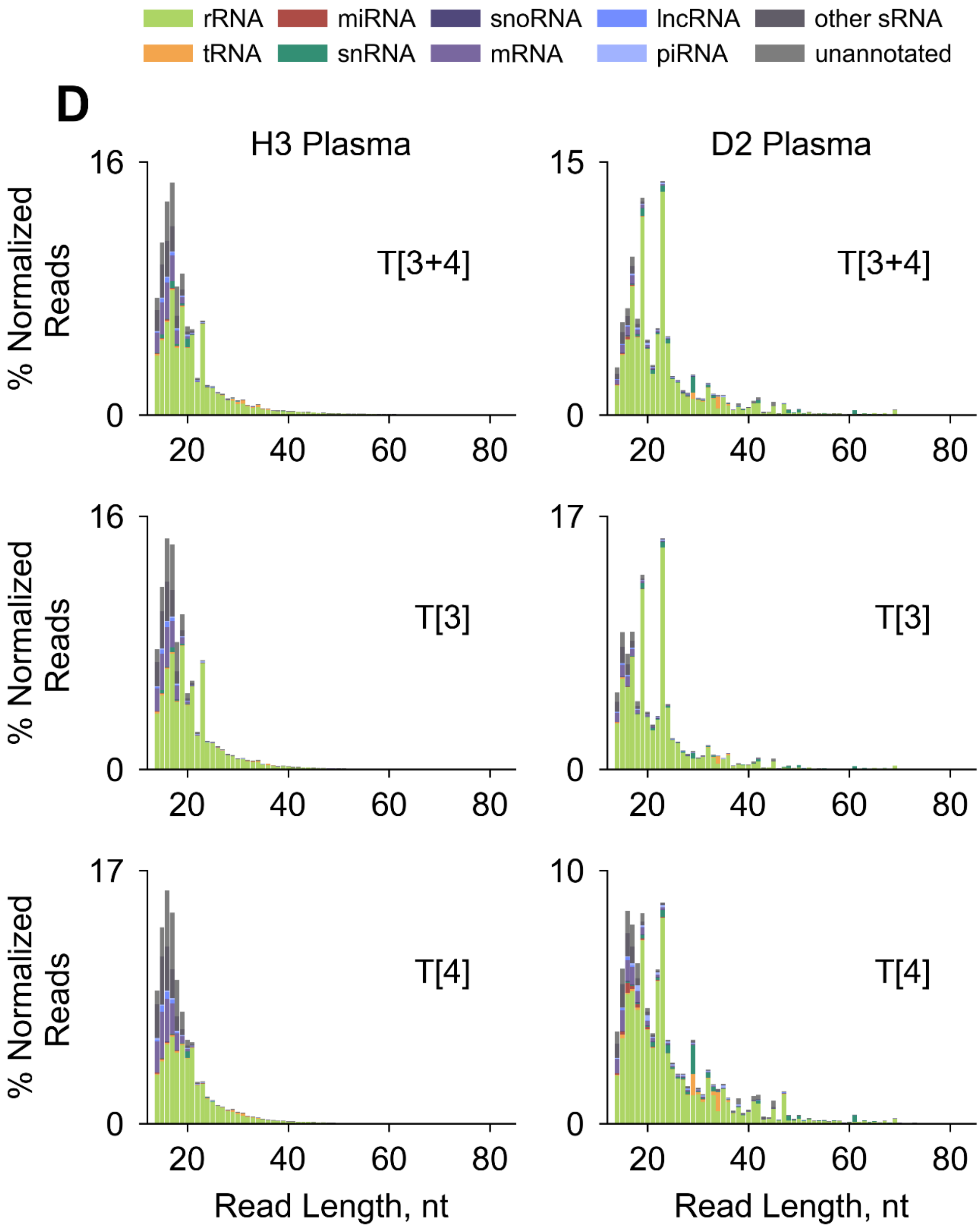

**Supplementary FIGURE S23A-B**

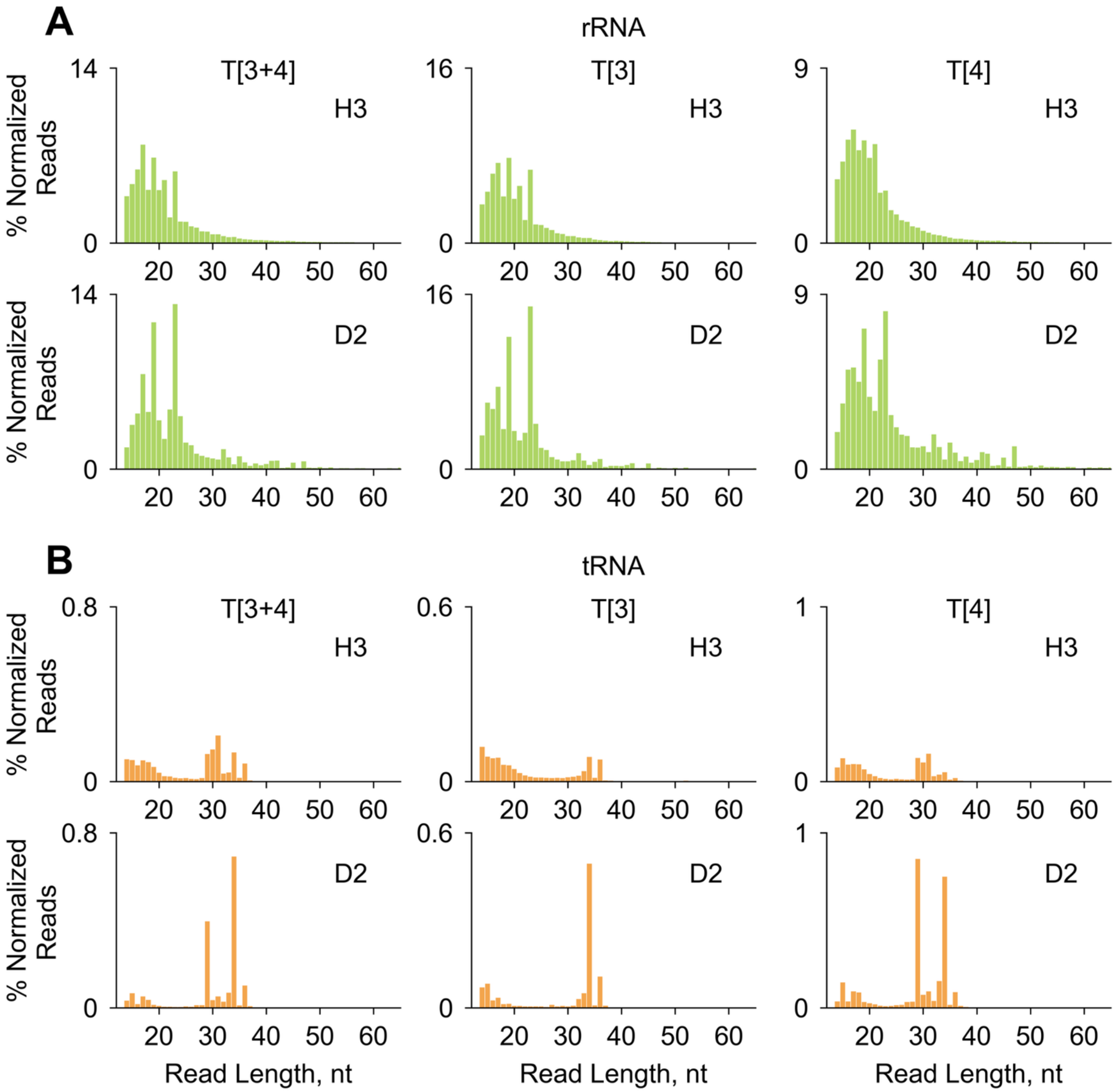

**Supplementary FIGURE S23C-D**

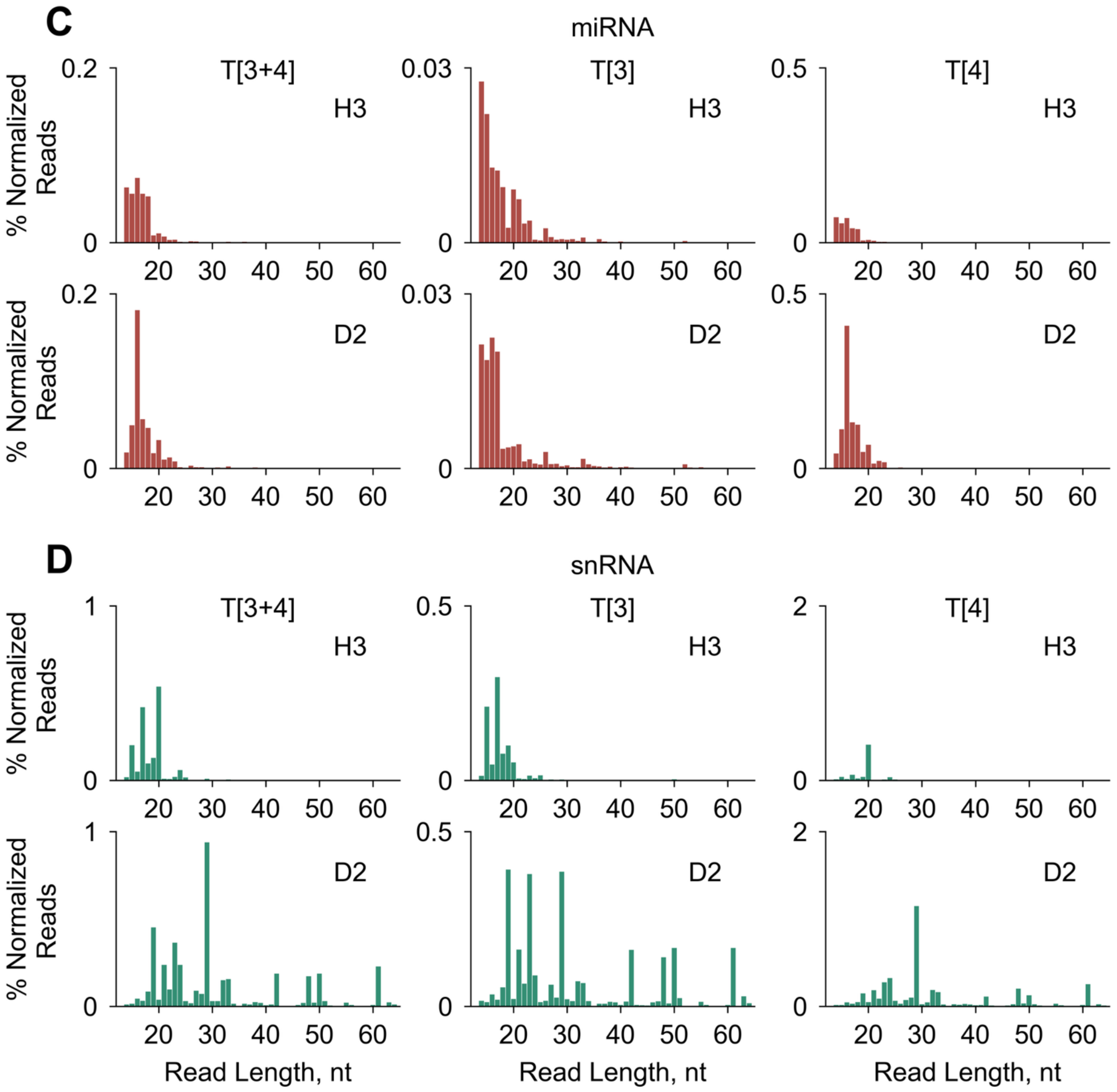

**Supplementary FIGURE S23E-F**

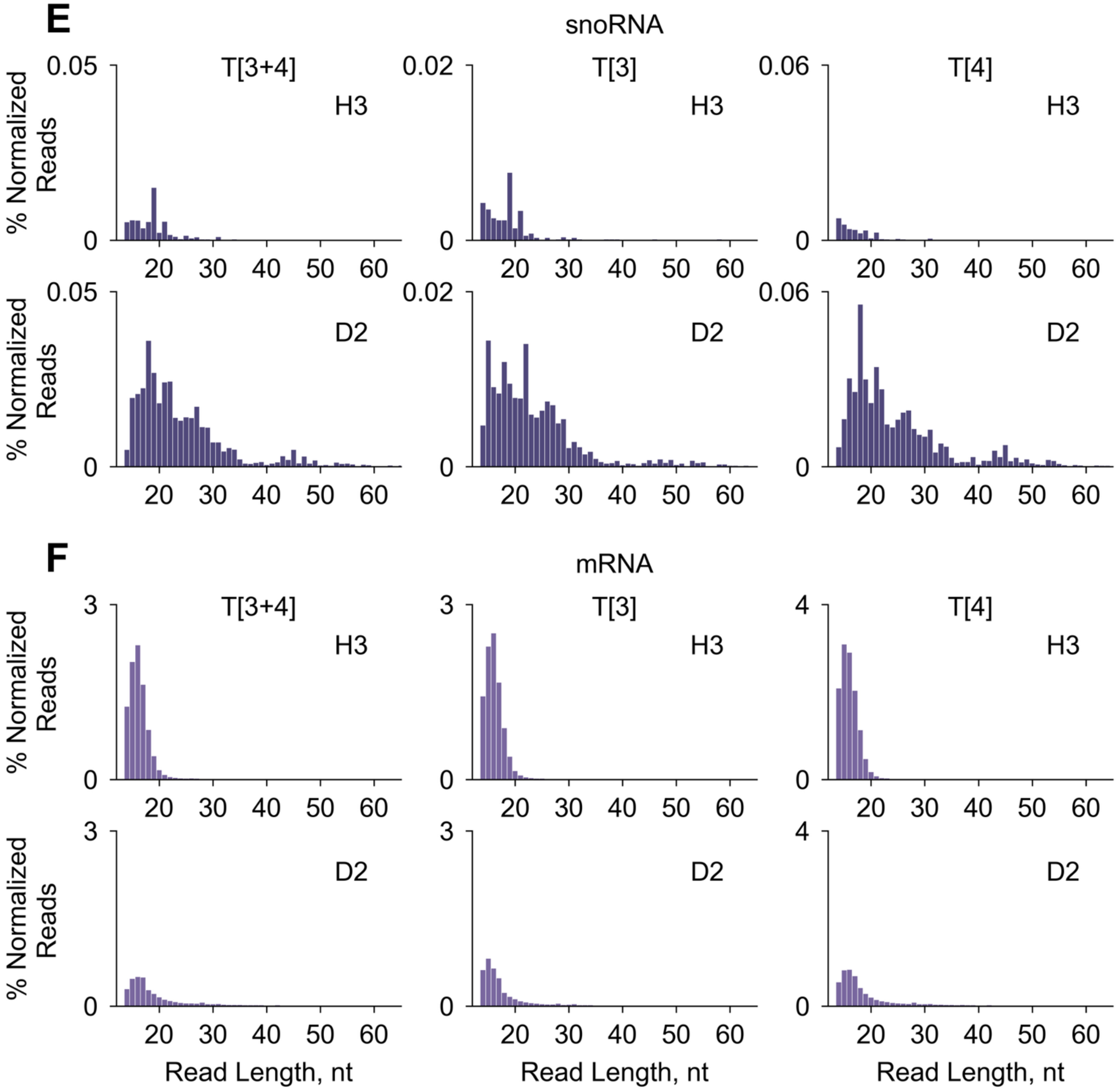

**Supplementary FIGURE S23G-H**

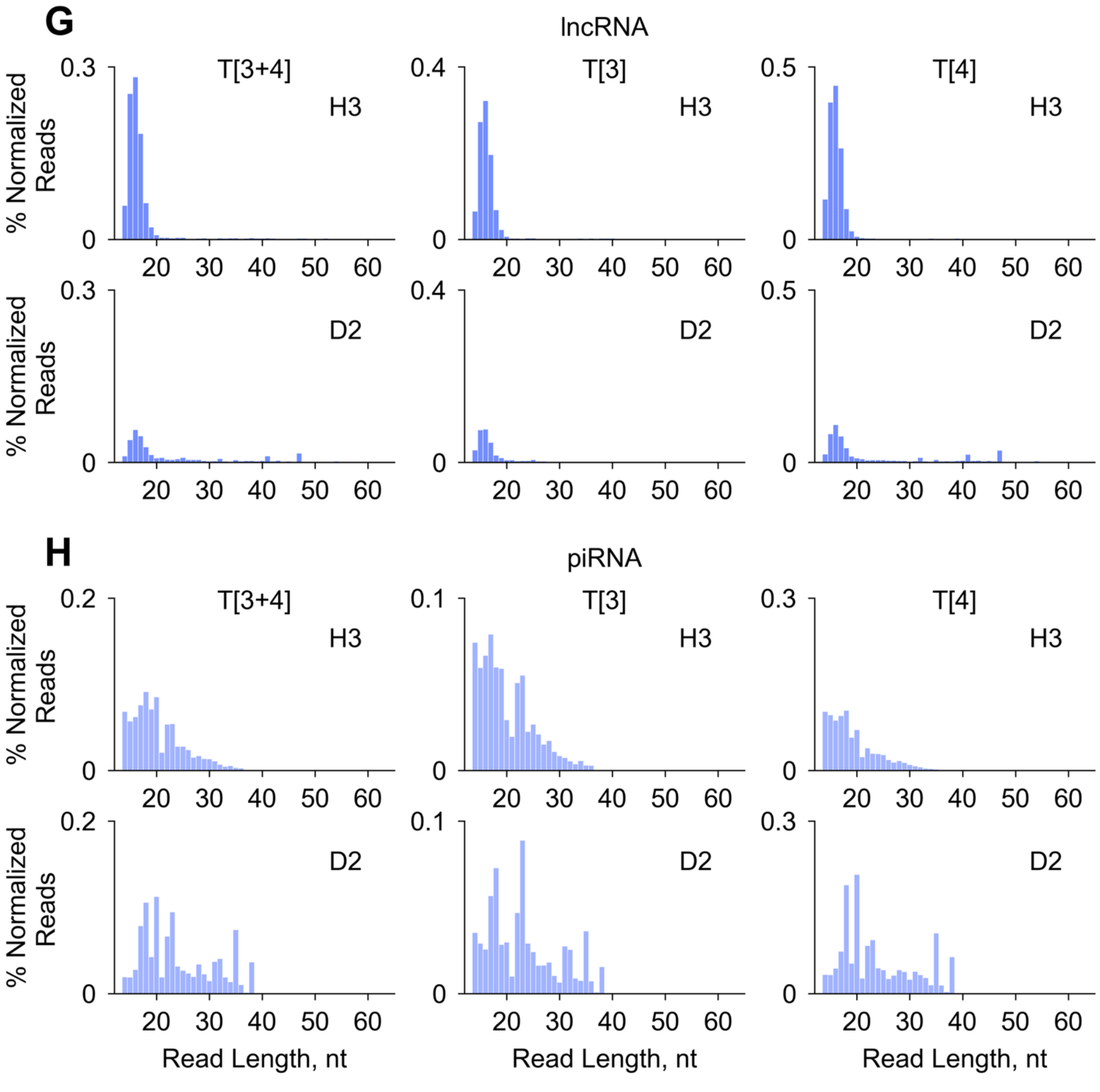

**Supplementary FIGURE S24**

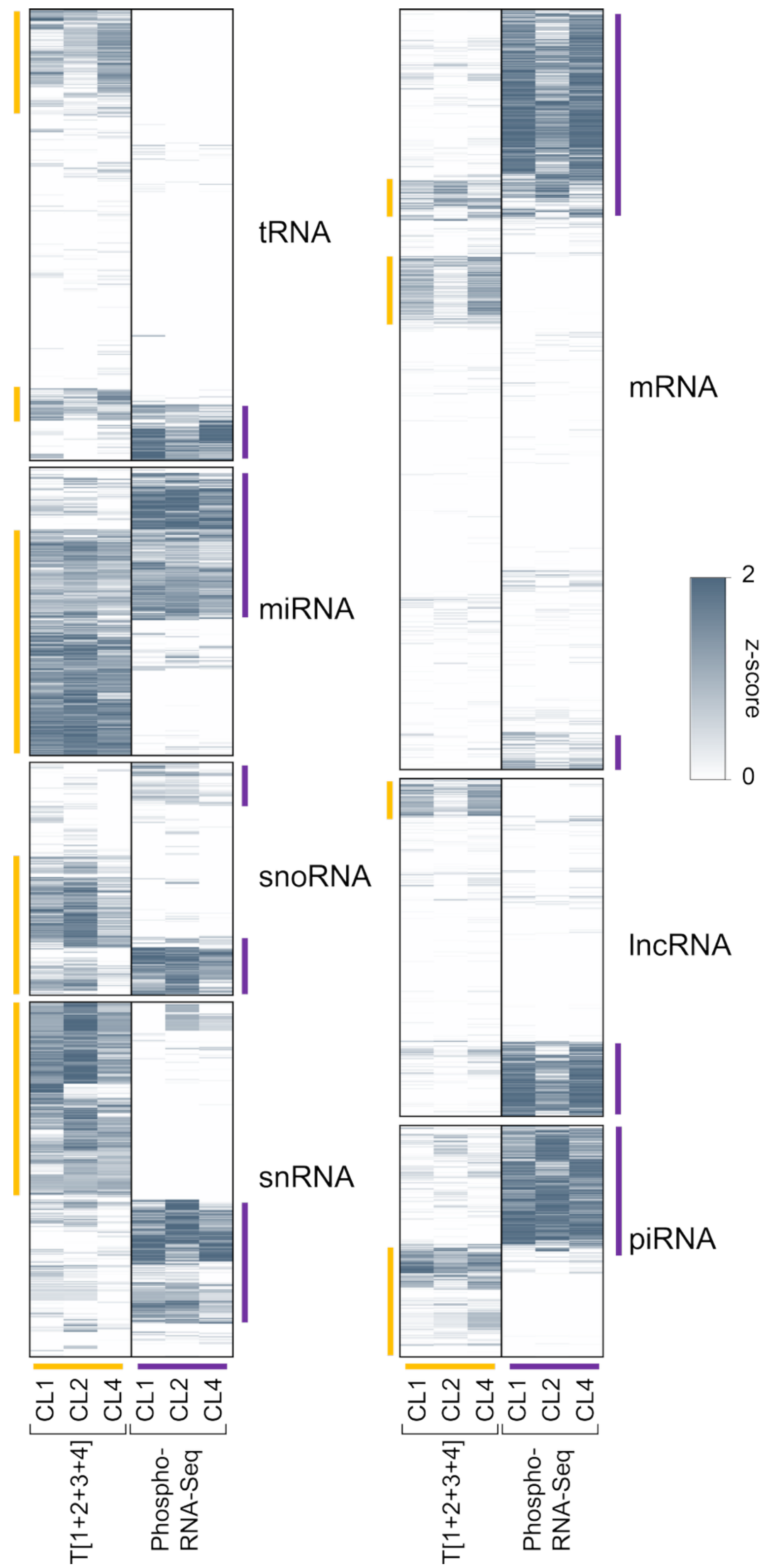

#### Supplementary FIGURE S25

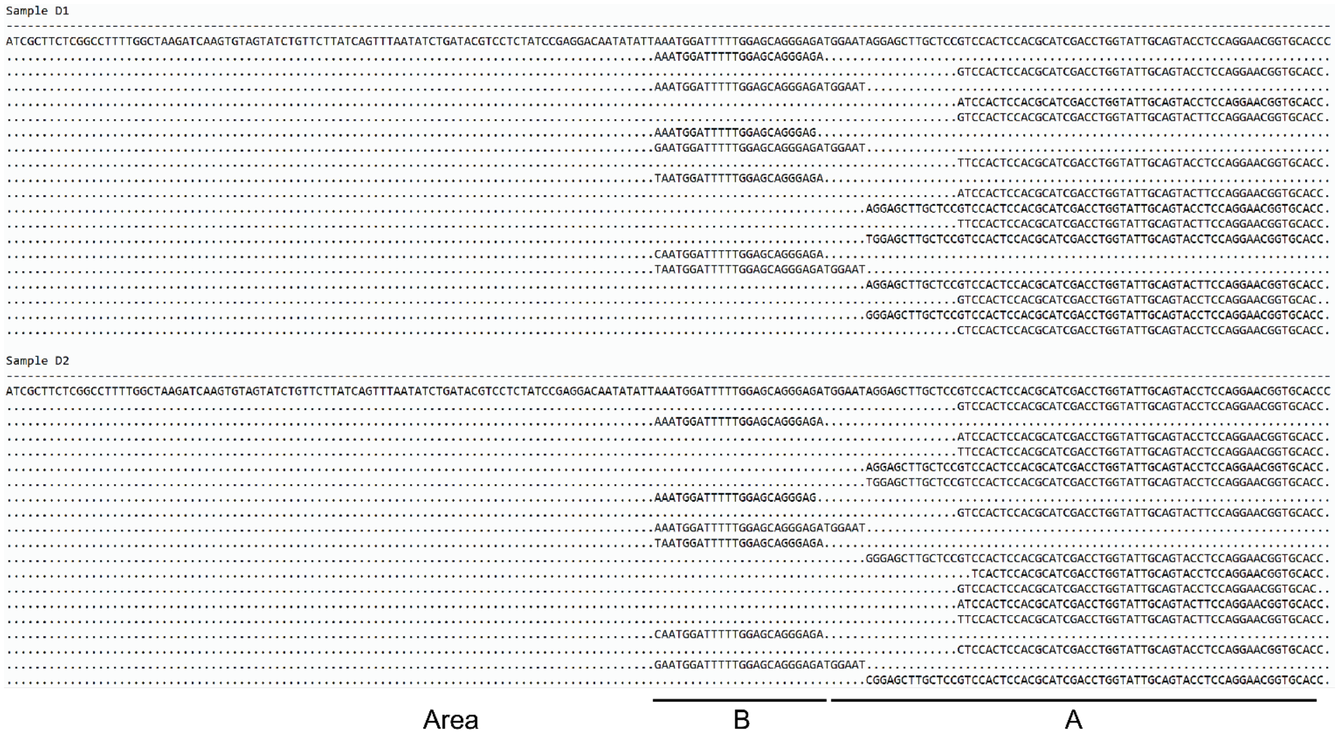
